# Display functions of dinosaur proto-wings before powered flight

**DOI:** 10.64898/2026.04.05.714230

**Authors:** Jinseok Park, Minyoung Son, Woojoo Kim, Yuong-Nam Lee, Sang-im Lee, Piotr G. Jablonski

**Author notes:** Corresponding authors: YL, SL, PGJ.

## Abstract

Pennaceous feathers are fundamental to avian flight, yet their early function in non-volant dinosaurs remains unknown. Early-diverging pennaraptorans had simple pennaceous feathers on proto-wings and tails, which were unsuitable for flight but may have enhanced visual signals. However, the visual display hypothesis has not been empirically tested. To address this, we used computer animations of early pennaraptoran displays to measure responses in a well-established animal model of a visually sensitive neural pathway. We show that pennaceous proto-wings and tails enhance the efficiency of motion-based displays across a range of anatomically plausible movements. Integrating these results with comparative and paleontological evidence, we suggest that early pennaceous feathers functioned in diverse signaling contexts and were subsequently exapted for aerodynamic use.

## Main

Pennaceous feathers are characterized by a stiff central rachis with vanes forming a broad surface. Wing feathers of modern birds are pennaceous and are adaptively modified for flight, featuring aerodynamic asymmetric vanes^1^. Early pennaceous feathers were present in early pennaraptoran dinosaurs before the origin of flight^4^ in the Middle Jurassic^15–17^. The best-preserved early pennaraptorans—*Caudipteryx* and *Protarcheopteryx*—show early pennaceous feathers on their tails and forelimbs, where they form proto-wings^2^. The small proto-wings^18^ in pennaraptora were distally located on forelimbs, prompting hypotheses about their potential use for flap-running^5^, drag/lift-based maneuvering^19,20^, wing-assisted inclined running^21^, or leaping takeoff^22^. However, early pennaraptorans had shoulder joint ranges of motion that were too narrow to generate sufficient aerodynamic forces^3^. Other functions have also been discussed^23–25^, including motion-based displays with pennaceous-feathered surfaces^6–8,17^ towards conspecifics, predators or prey. The signaling function is especially likely because the plesiomorphic simple filamentous feathers likely already functioned as visual signals^26^. The pennaceous feathers may have enhanced these early displays, especially on the limbs and on the tail, which can be used in faster and more extensive movements compared to other body parts. The stiff central rachis with wide vanes could effectively increase the display area and help maintain shape during rapid movement. This visual display hypothesis^8^ has not been directly tested, presumably because strictly speaking it concerns responses from extinct organisms that produce and receive display signals. However, using modern analogs—the visual signals of modern birds (extant Pennaraptora)—may offer fresh insights.

One of the visual signaling contexts concerns displays towards prey, i.e., flush-displays^9^, observed in diverse birds—such as the [greater roadrunner](https://www.youtube.com/watch?v=qmXGDWkZSek) (*Geococcyx californianus*), the [northern mockingbird](https://macaulaylibrary.org/asset/200776531) (*Mimus polyglottos*)^27^ or the [rufous-tailed scrub robin](https://www.youtube.com/watch?v=pS31OI4YNdw) (*Cercotrichas galactotes*). The birds [innate](https://youtu.be/TVVVRIZIEaM)ly^28^ move their wings and tail in species-specific manners to visually flush the prey (hence the term “flush-displays”^9^). This may increase foraging success by startling and exposing prey to the predator (thus increasing the number of prey available for pursuit), or by influencing the focal prey’s escape timing or direction for easier capture (Video S1)^29,30^. The functional role of flush-displays has been experimentally reproduced by a robot imitating mockingbird behaviours^31^. Expanding from this approach, experiments with a robotic early pennaraptoran revealed that the presence of pennaceous feathers may have enabled effective motion-based flush-displays in early pennaraptorans^9^, despite their relatively narrow range of anatomically feasible display movements^9,32,33^ compared to birds. Hence, proto-wings and tail feathers may have enhanced the foraging success of the hypothetical pennaraptorans that pursue the flushed prey. However, a complete evaluation of various movement patterns in the flush-display was not possible due to the mechanical limitations of the robot. Examining responses of prey escape neurons to animations of dinosaurian flush displays offers an alternative to recording prey jumping in response to robotic dinosaurs and overcomes the robot’s limitations. The orthopteran LGMD/DCMD (Lobula Giant Movement Detector/Descending Contralateral Movement Detector) pathway is a well-studied model of neural pathways sensitive to looming stimuli and to accelerated edge translation^34–40^. DCMD neural responses can predict the probability of escape behaviour^41^ and provide a more quantifiable measure of display efficacy than simple binary (escape or non-escape) behaviors. This neuronal pathway serves not only as a model circuit for escape responses to approaching objects but also to displays involving looming and acceleration^9^, thus enabling quantification of the effects of the display on the receiver.

Here, we created computer animations of hypothetical flush-displays in early-diverging pennaraptorans. The animations are based on the flush-display motions of three modern flush-pursuing birds (see Methods and SI Part 1). We recorded DCMD responses in locusts to quantify the effectiveness of proto-wing and tail displays in stimulating the receivers. Based on our experimental results and an overview of visual signaling in extant Sauropsida across diverse contexts, we propose a comprehensive framework for the hypothetical roles of diverse visual displays in the origin and subsequent evolution of pennaceous feathers in the proto-wings and tails of pennaraptoran dinosaurs.

### Effects of the proto-wing and the feathered tail in flush-displays

Pilot experiments using animations of an early pennaraptoran model (Video S2; SI Part 2), as well as previous experiments using robots^9,31^, confirmed the expected link between the occurrence of the orthopteran escape jumps and their DCMD responses to animations of displays (Extended Data Fig. 1; SI Part 2). Subsequently, we used the peak size and the frequency of reaching the escape-preparation threshold in locust’s DCMD spiking rates as proxies of the overall effect of the visual stimulus on orthopteran escape behaviour. Three different hypothetical flush displays by an early pennaraptoran were simulated on the computer screen, either with or without feathered proto-wings and tail, at short versus long distances from the locust (detailed Methods are in SI Part 1). The full set of results is presented in SI Parts 3, 4 5.

In response to animations based on the wing displays of the greater roadrunner (Exp. 1; Video S3; SI Part 1D), DCMD activity showed three distinct peaks in spiking rates corresponding to the three display phases (Figs. 1a, b, d and S3; Extended Data Fig. 2). Peak spiking rates were consistently higher when the feathers were present and when the display occurred at a closer distance (Fig. 1c; Table 1a), and the frequency of reaching the escape threshold was higher with the feathers but it was unaffected by the distance (Fig. 1e–f; Table 1b). In animations based on the wing and the tail displays of the rufous-tailed scrub robin (Exp. 2; Video S4; SI Part 1E), six display phases were included, with clear DCMD peaks for later four phases across the four animations (Figs. 1h, i, k and S4; Extended Data Fig. 3). Presence of feathers increased peak spiking rates and the frequency of reaching the escape threshold, and these responses were stronger at a close distance (Fig. 1j, l–n; Table 1c, d). Presentation of animations based on the wing display of the northern mockingbird (Exp 3; Video S5; SI Part 1F) resulted in four distinct spiking rate peaks corresponding to the four display phases (Fig. 1o, p, r and S5; Extended Data Fig. 4). A significant interaction between feather presence and distance indicates that the feathers significantly increased peak spiking rates only when the display was distant (Fig. 1q; Table 1e). The frequency of reaching threshold remained higher at close distance, regardless of feather presence (Fig. 1s–u; Table 1f). When we added a ‘walking’ movement at the beginning of the animation before the flush display (Exp. 4; Video S6; the same display as used in Exp. 1), the feathers still increased the peak spiking rates (Extended Data Fig. 5a, b, c; Table S9), but the walking movement caused an overall decrease in the peak spiking rates (Extended Data Fig. 5c; Table S9; SI Part 4). Taken altogether, all these results indicate that the presence of proto-wings and tails with simple pennaceous feathers enhances the effectiveness of various types of flush-displays.

**Figure 1.**
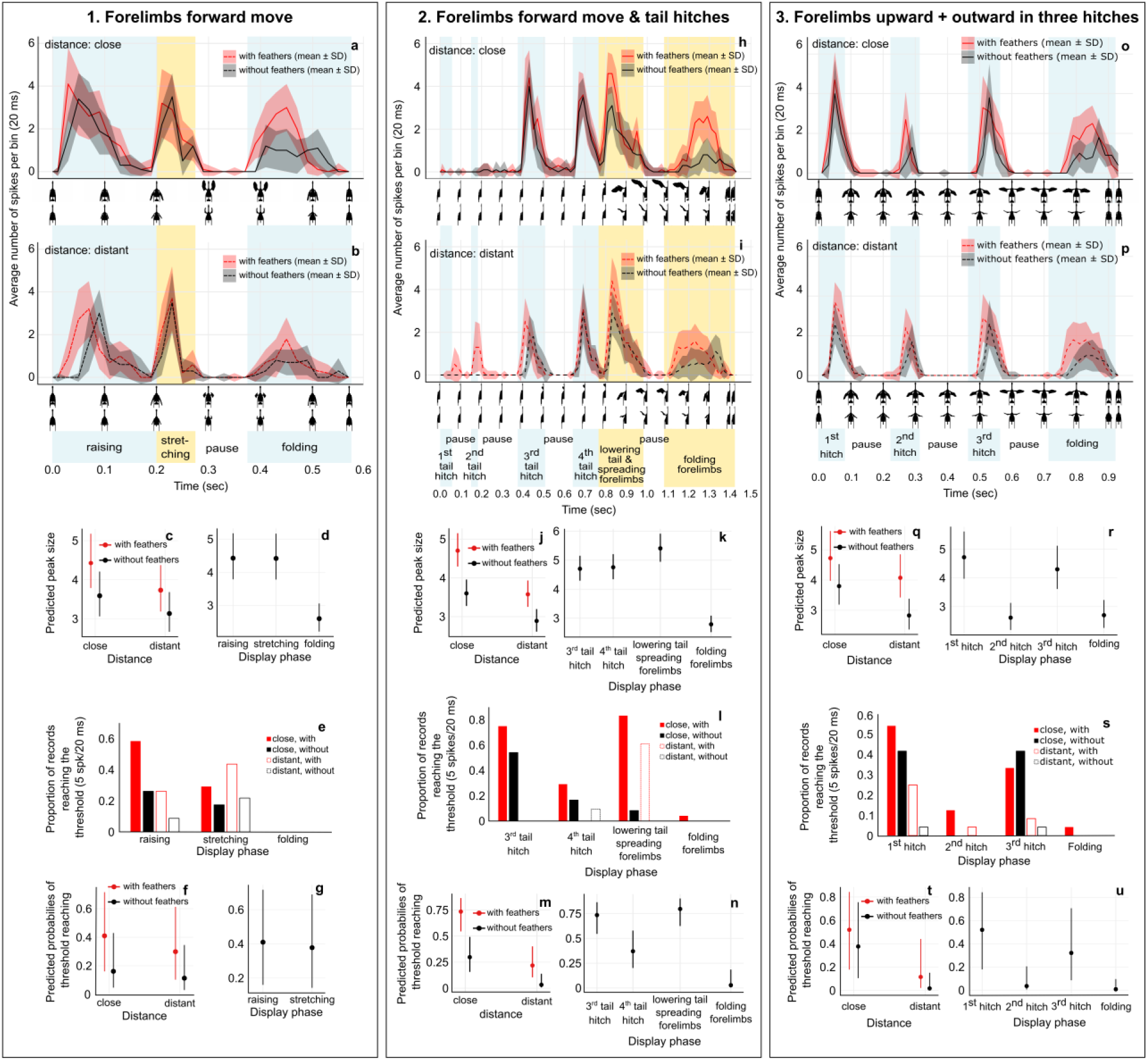
Response of locust DCMD to three types of hypothetical dinosaurian flush-displays based on displays of *Geococcyx californianus* (a–g), *Cercotrichas galactotes* (h–n), and *Mimus polyglottos* (o–u). **a**, **b**, **h**, **i**, **o**, **p**, DCMD responses during animations with (red) and without feathers (black) at close (60 cm; **a**, **h**, **o**) and distant (120 cm; **b**, **i**, **p**) simulated locations (n = 24: four recordings from each of six individuals). Blue or yellow shading indicates display phases, while a lack of shading represents pauses. **c**, **j**, **q**, Effect of presence of feathers and distance on the peak spiking rate (no. spikes/20 ms). **d**, **k**, **r**, Effect of display phase on the peak spiking rate (no. spikes/20 ms). **e**, **l**, **s**, Effect of feather presence and distance on the proportion of records (n = 24) reaching the threshold value (5 spikes/20 ms^41^) for jump preparation at each display phase. **f**, **m**, **t**, Effect of presence of feathers and distance on the probability of reaching the threshold (5 spikes/20 ms^41^) for jump preparation. **g**, **n**, **u**, Effect of display phase on the probability of reaching the threshold for jump preparation. In **c**, **d**, **f**, **g**, **j**, **k**, **i**, **m**, **n**, **t**, **u**, dots indicate predicted values, while vertical lines represent 95% confidence intervals derived from the statistical models presented in Table 1. The ‘folding’ phase is not presented in **g**, because no records reached the threshold. For **h** and **I**, DCMD responses to the first and second phases were visible only in the distant/feathers-present treatment. Details on DCMD responses are shown in SI Part 3, with corresponding animations in Videos S2–S4.

**Table 1.** Results of generalized mixed-effects models for the three experiments. **a, b**, Animation based on *Geococcyx californianus* in exp.1; **c**, **d**, animations based on *Cercotrichas galactotes* in exp. 2; **e**, **f**, animation based on *Mimus polyglottos* in exp. 3. For the under-dispersed no. spikes per 20 ms, we utilized the Conway-Maxwell poisson family with a log link function. For the binary threshold (reached = 1, unreached = 0), we used the Binomial distribution with a logit link function. The fixed effects include ‘feather’ (*with* or *without*, the former is the reference level), ‘distance’ (*close* or *distant*, the former is the reference level), and the ‘feather’ x ‘distance’ interaction. The additional fixed effect of ‘display type’ had levels corresponding to the number of display phases in each experiment: 3 levels in exp. 1 *(raising*, *stretching*, *folding*), 4 levels in exp. 2 (*third tail hitch, fourth tail hitch, lowering the tail and spreading forelimbs, folding forelimbs*), and 3 levels exp. 3 (*first hitch, second hitch, third hitch,* wing *folding*). In the analyses of ‘threshold’ in exp. 1, the display type had only 2 levels (*raising*, *stretching*) because no threshold-reaching event occurred in the final phase. Random effects include individual ID and record ID nested within individual ID. P-values below 0.05 are denoted in bold, except for the intercept.

| Response variable | Fixed effects | Estimate | SE | Z value | P-value |
| --- | --- | --- | --- | --- | --- |
| a) Exp. 1:<br>Peak size<br>Fig. 2c, d | Intercept | 1.488 | 0.080 | 18.693 | < 0.001 |
|  | **Feather** ‘without’ | -0.210 | 0.044 | -4.744 | < **0.001** |
|  | **Distance** ‘distant’ | -0.171 | 0.044 | -3.894 | < **0.001** |
|  | Feather ‘without’:distance ‘distant’ | 0.035 | 0.066 | 0.532 | 0.595 |
|  | Display phase ‘stretching’ | -0.002 | 0.037 | -0.049 | 0.961 |
|  | **Display phase** ‘folding’ | -0.533 | 0.044 | -12.235 | < **0.001** |
| b) Exp. 1:<br>Threshold<br>Fig. 2f, g | Intercept | -0.365 | 0.659 | -0.554 | 0.580 |
|  | **Feather** ‘without’ | -1.286 | 0.508 | -2.529 | **0.011** |
|  | Distance ‘distant’ | -0.488 | 0.476 | -1.026 | 0.305 |
|  | Feather ‘without’:distance ‘distant’ | 0.078 | 0.745 | 0.104 | 0.917 |
|  | Display phase ‘stretching’ | -0.134 | 0.364 | -0.368 | 0.713 |
| c) Exp. 2:<br>Peak size<br>Fig. 2j, k | Intercept | 1.548 | 0.046 | 33.311 | < 0.001 |
|  | **Feather** ‘without’ | -0.267 | 0.035 | -7.663 | < **0.001** |
|  | **Distance** ‘distant’ | -0.274 | 0.035 | -7.727 | < **0.001** |
|  | Feather ‘without’:distance ‘distant’ | 0.054 | 0.055 | 0.980 | 0.327 |
|  | Display phase ‘fourth tail hitch’ | 0.011 | 0.036 | 0.307 | 0.758 |
|  | **Display phase** ‘lowering the tail and spreading forelimbs’ | 0.139 | 0.035 | 3.986 | < **0.001** |
|  | **Display phase** ‘folding forelimbs’ | -0.523 | 0.043 | -12.182 | < **0.001** |
| d) Exp. 2:<br>Threshold<br>Fig. 2m, n | Intercept | 1.021 | 0.427 | 2.389 | 0.017 |
|  | **Feather** ‘without’ | -1.879 | 0.408 | -4.605 | < **0.001** |
|  | **Distance** ‘distant’ | -2.297 | 0.437 | -5.254 | < **0.001** |
|  | Feather ‘without’:distance ‘distant’ | -0.229 | 0.886 | -0.259 | 0.796 |
|  | **Display phase** ‘fourth tail hitch’ | -1.550 | 0.444 | -3.491 | **0.0004** |
|  | Display phase ‘lowering the tail and spreading forelimbs’ | 0.347 | 0.374 | 0.929 | 0.353 |
|  | **Display phase** ‘folding forelimbs’ | -4.552 | 1.068 | -4.264 | < **0.001** |
| e) Exp. 3:<br>Peak size<br>Fig. 2q, r | Intercept | 1.553 | 0.088 | 17.592 | < 0.001 |
|  | **Feather** ‘without’ | -0.219 | 0.048 | -4.575 | < **0.001** |
|  | **Distance** ‘distant’ | -0.148 | 0.047 | -3.143 | **0.002** |
|  | **Feather** ‘without’:distance ‘distant’ | -0.147 | 0.071 | -2.074 | **0.038** |
|  | **Display phase** ‘second hitch’ | -0.592 | 0.044 | -13.305 | < **0.001** |
|  | **Display phase** ‘third hitch’ | -0.093 | 0.038 | -2.482 | **0.013** |
|  | **Display phase** ‘folding’ | -0.559 | 0.044 | -12.732 | < **0.001** |
| f) Exp. 3:<br>Threshold<br>Fig. 2t, u | Intercept | 0.079 | 0.820 | 0.096 | 0.924 |
|  | Feather ‘without’ | -0.583 | 0.557 | -1.048 | 0.295 |
|  | **Distance** ‘distant’ | -2.115 | 0.688 | -3.076 | **0.0021** |
|  | Feather ‘without’:distance ‘distant’ | -1.399 | 1.092 | -1.281 | 0.200 |
|  | **Display phase** ‘second hitch’ | -3.397 | 0.732 | -4.638 | < **0.001** |
|  | Display phase ‘third hitch’ | -0.830 | 0.446 | -1.860 | 0.063 |
|  | **Display phase** ‘folding’ | -4.958 | 1.161 | -4.271 | < **0.001** |

Looming-sensitive fast-escape circuits with properties similar to the LGMD/DCMD pathway are known across a wide range of invertebrate and vertebrate taxa^42–46^ including prey of flush-pursuing birds, suggesting that various animal taxa may have been similarly affected by the visual flush-displays of early pennaraptorans possessing proto-wings and tails (Fig. 3a–c). As expected from the properties of the LGMD/DCMD pathway^37, 47^, the strongest alignments in both timing and magnitude of neural responses were found with the acceleration of white-to-black pixel transitions (viewed on a log scale in Fig. 2; Extended Data Figs. 6–9; see SI Part 3B–E for other visual variables). These results indicate that accelerated movements of forelimbs may have been sufficient for flush-displays in early pennaraptorans, even with their small proto-wings and limited motion ranges compared to modern birds (see SI Parts 6A, B for details). Thus, we hypothesize that flush-displays could be more effective in later-diverging pennaraptorans with their increased pectoral range of motion, especially in paravians with laterally oriented glenoid fossae, and finally in Ornithothoraces where the glenoid fossa is oriented dorsally as in modern birds^48^.

**Figure 2.**
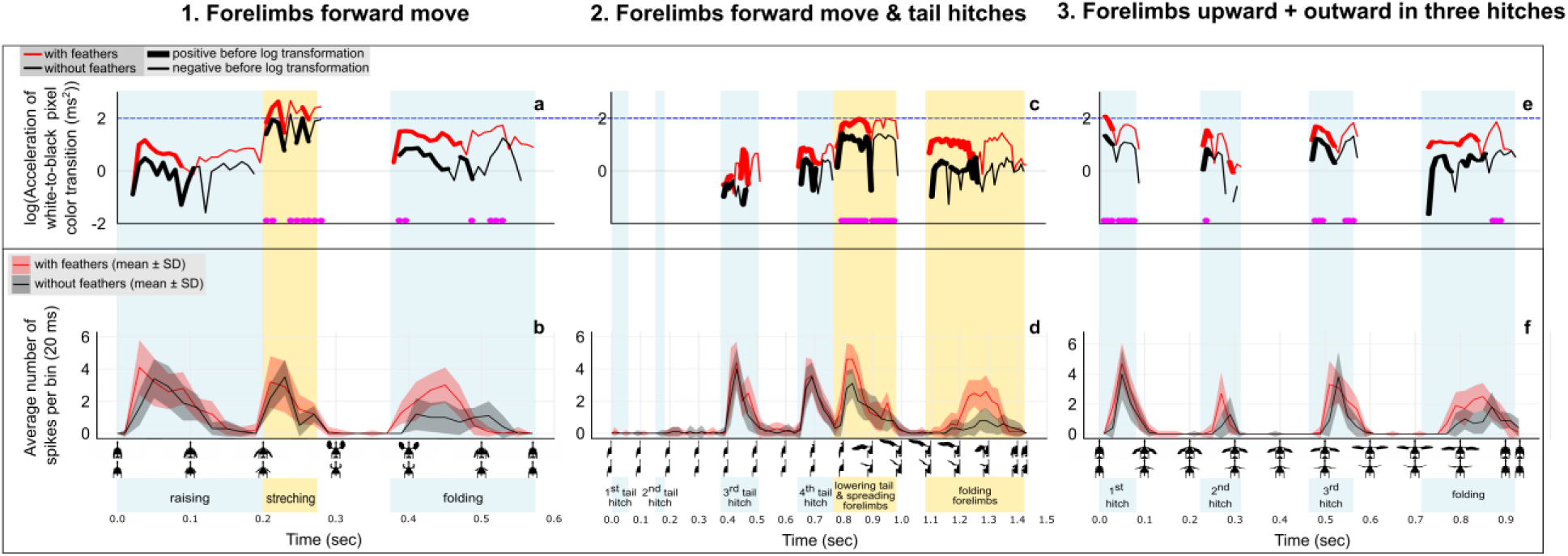
Profiles of acceleration in the change of the numbers of pixels with color transitions in dinosaur animations, and locust DCMD responses to those animations. **a**, **c**, **e**, Acceleration of the change in the number of pixels with color transition from white to black in dinosaur animations with (red) and without (black) feathers at a close distance, illustrating difference between the display types. The acceleration axis uses a log-scale, with positive values (before applying the log scale) shown as thick lines and negative values as thin lines. This is part of a comprehensive comparison between DCMD responses and profiles of several variables extracted from the movements of the animated dinosaur (Extended Data Figs. 6–8; SI Part 3D). To illustrate differences between the three display types in triggering high visual acceleration in the prey field of view, the blue dashed line and the bottom set of pink dots mark acceleration values of 2 and 1.5, respectively. In **c**, the first and second tail hitches are invisible in close distance animations. **b**, **d**, **f**, DCMD responses during animations with (red) and without (black) feathers at a close simulated location (n = 24: four recordings from each of six individuals). Blue or yellow shading indicates display phases, while a lack of shading represents pauses. Details on the animations and DCMD responses are shown in SI Part 1 and Part 3, respectively.

**Figure 3.**
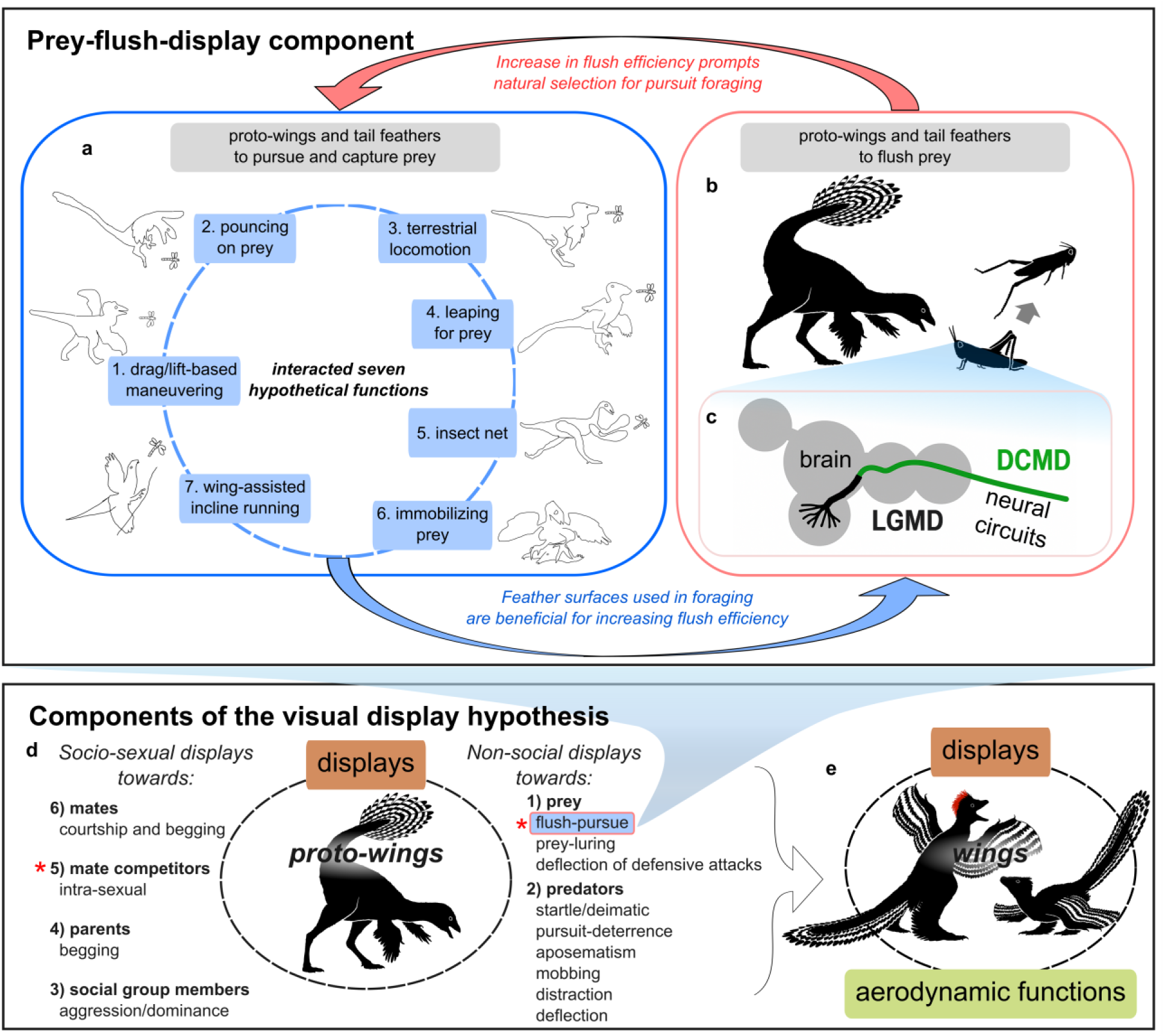
Schematic illustration of the flush-pursue visual-display component (a–c) among various other components comprising the visual display hypothesis (d) and their hypothetical link to the evolution of flight (e). **a**, The role of feathers in pursuit and capture of prey by prey-pursuing predators^9^: 1. drag- and lift-based maneuvering during terrestrial pursuits^19^, 2. drag-based maneuvering while pouncing on prey^20^, 3. enhancement of running^5^, 4. enhancement of leaping^22^ after prey^3^, 5. use as insect nets^24^, 6. immobilization of prey^23^, 7. running after prey on inclined substrates rather than running from predators, as originally proposed^21^. Recently the functions 3, 4, and 7 were determined as less likely before Paraves stage^3^. Increasing feather surface and stiffness for one function also enhances others, leading to the concerted evolution of traits crucial for a cursorial prey pursuer. **b**, The role of feathers in flushing prey: increasing the area or contrast in the proto-wing and the tail enhances the visual stimulation of prey escape neurons (**c**), leading to more frequent prey escapes. This leads to more frequent use of prey pursuits (**a**), which are also aided by increased feather surfaces and limb movement capabilities. Hence, the development of feathers for signaling function could lead to improvements in prey-pursue function, and vice versa (blue and red arrows indicate this reinforcing mechanism of proto-wing and forelimb evolution). **d,** The visual display hypothesis proposes that early pennaceous feathers were initially used for displays in various signaling contexts. An increase in visual signal in one context may have enhanced its effectiveness in other contexts. Red asterisks indicate contexts where proto-wing/forelimb movements may also play non-signaling functions intrinsically linked with visual signaling: pursuing prey flushed by flush-displays or attacking opponents and running away from opponents in visual-display contests. **e,** Once relatively large, feathered areas had evolved, they may have later been exapted for aerodynamic functions. Drawings in (a) by J. Park; other illustrations by Choi Yu-sik. Relevant literature review and discussion are provided in SI Part 6.

### Motion-based displays and the function of early pennaceous feathers

Our empirical results illustrated that the orthopteran escape circuits respond to key motion stimuli, such as acceleration and looming, present in the flush-displays. Similar stimuli are present in diverse displays used by animals in other signaling contexts, and neuro-sensory circuits for visual motion processing are generally sensitive to those stimuli^49–51^. Therefore, our empirical results from the specific flush-display context are also relevant to the proposed hypothetical diverse signaling functions of pennaraptoran proto-wings and tails^6–8,17^. These signals could include displays directed at conspecifics, predators, or prey. Therefore, the visual display hypothesis (Fig. 3d; SI Part 6C) proposes that multiple signaling functions (Fig. 3d) of proto-wings and tails are observed, and these functions can interact in shaping feathered display surfaces and forelimb/tail movements during evolution. Such interactions could contribute to the observed diversity^3^ and inferred multifunctionality of exaggerated, motion-based displays by proto-wing and tail feathers across Pennaraptora with diverse ecological niches^52,53^.

Motion-based displays, including tail and limb movements, are common among extant Sauropsida across diverse signaling contexts as indicated by our review of literature (Table S10; SI Parts 6D–F). Use of tail and limb movements in multiple signaling contexts is especially well described in lizards and birds (SI Part 6D-F). Hence, based on phylogenetic bracketing^54^, it is likely that dinosaurs also used motion-based displays. Bite marks observed in tyrannosauroids^13^, and ground scrape trace fossils^14^ led to speculations about signaling behavior on leks suggesting that theropods might have used visual displays similar to those by extant lekking birds (examples in SI Part 6G). These speculations about the behavior are based on the relatively weak evidence that could have alternative explanations. However, the prevalence of anatomical display structures in dinosaurs supports the idea that visual signaling was a potent selective context for the evolution of integumentary innovations^55^. For example, conspicuous integuments in the form of monofilamentous bristles were present in ceratopsians^56^, and cranial ornaments—likely used in various display functions^55^—were widespread among Ornithodira, including Ornithischians, non-paravian theropods and pterosaurs^55, 59^. While maniraptoriform theropods generally lacked cranial ornaments, they possessed feather-like filaments and feathers^57^, which developed into elaborate pennaceous surfaces on the forelimbs and tail in Pennaraptora^2, 6^ and seem to have taken over the visual signaling functions^60^. Coloration patterns in the pennaceous plumage of *Caudipteryx*^2^ and *Anchiornis*^6^, together with a noticeable increase in melanosome diversity at the base of Pennaraptora^58^, also support the signaling function of colored pennaceous feathers.

Evolutionary trends among pennaraptorans leading to birds also support the view that early pennaceous proto-wings and tails might have initially served diverse signaling functions. Fossil evidence shows that pennaceous feathers and even wing-like large surfaces evolved before the appearance of distinct postcranial anatomical features that may have allowed powered flight^24^, consistent with non-aerodynamic roles, such as visual signaling. Additionally, the absence of a consistent trend in wing area in non-avian pennaraptoran lineages is incongruent with aerodynamic functions^3^ but aligns well with display functions of early pennaceous feathers, because different mixtures of diverse signaling contexts are expected to produce different sizes of visual signals. This scenario is further mirrored in the ontogeny of *Microraptor*, *Archaeopteryx*, and some living birds, in which large wing surfaces develop before the formation of the strong pectoral muscles^61^ required for flight. Correspondingly, anatomical changes in the pectoral girdle^48^ occurred before any aerodynamic advantages emerged, and changes in wrist anatomy in Pennaraptora^62^ possibly allowed bird-like “automated” flexion of the wrist and elbow before the acquisition of flight. These anatomical transformations of forelimb are also generally consistent with components of the display hypothesis because they could enable faster and more elaborate motion-based displays. The adaptations to motion-based displays with proto-wings and tails might have been later exapted for aerodynamic advantages. We suggest that some specific components of the visual display hypothesis (Fig. 3d) are especially likely to have played a role in this process. The flush-display context (Fig. 3a–c) has been proposed^9^ to uniquely combine signaling (Fig 3b) and non-signaling functions (Fig. 3a) of the same phenotypic features. These functions are intrinsically linked with each other owing to the nature of this signaling component of the hypothesis (Fig. 3a, b; SI Part 6B). As some of the non-signaling functions (Fig. 3a) involve drag and possibly lift (e.g., 1, 2, 4 in Fig. 3a), the flush-display component of the visual display hypothesis might have prompted the process towards the exaptation of proto-wings and tails for aerodynamic functions important in flight (Fig. 3e; SI Part B). Similar set of intrinsically linked display functions of proto-wings and their nonsignaling functions is likely in the context of terrestrial displays on leks suggested by behavior of modern birds, where the use of wings for displays is associated with their use for accelerated movements and maneuvering during chases and leaps (SI Part 6G).

In summary, the results of our experiments and the review of existing evidence suggest that the early function and the subsequent early evolution of pennaceous proto-wings and tails in pennaraptoran dinosaurs likely occurred in the context of diverse and multifunctional motion-based displays (Fig. 3d). These signaling surfaces may have been later exapted for aerodynamic advantages (Fig. 3e) in paravians^48,63^, potentially facilitated by reinforcing mechanisms of proto-wing and forelimb evolution in those signaling contexts, in which the signaling and non-signaling functions are intrinsically linked such as in the flush-display component of the hypothesis. Future studies similar to our experiments could empirically evaluate the effect of proto-wing-like surfaces in motion-based socio-sexual displays on the behavioral or neuronal responses of extant sauropsid receivers (e.g., lizards^64^), further providing evidence relevant to the visual display hypothesis.

## Methods

### Pilot experiments

To confirm that the experimental setup targeted the DCMD neuron and triggered escape responses in the subjects, we generated a black circle looming stimulus (*l*/|*v*| = 5 ms) and tested whether this stimulus elicited escape jumps in our subjects (n = 5 locusts). For two locusts that reliably jumped in response to the stimulus, we recorded DCMD activity to observe the single spike shapes and the spiking rate profile in response to the looming stimulus. To verify that the relatively complex dinosaur animations trigger escape behaviour in locusts, we recorded the behavioural reactions of six locusts to dinosaur animations involving movements of feathered forelimbs and tail, loosely based on the rufous-tailed scrub robin’s flush displays. These animations were presented at two distances: 70 and 120 cm. We then assessed whether animations induced higher peak spiking rates and more frequent reaching of the hypothetical threshold DCMD spiking rate for jump preparation^41^ (see Data analysis section), resulting in more frequent behavioural escape reactions. Details are described in SI Part 1B.

### Computer animations of hypothetical flush displays by early pennaraptoran dinosaurs

#### Generating animations

We used Blender software (version 3.2.0, Blender Foundation) to generate a model demonstrating a hypothetical early pennaraptoran flush display. This 3D dinosaur model was based on *Caudipteryx* because of the existence of multiple fossil specimens and relatively well-documented morphology. Detailed justification for the choice of *Caudipteryx*, covering fossil specimen selection, reconstruction of proto-wing and tail feather morphology, and estimation of anatomical constraints on display movements, is presented in Part 1 of the Methods section in Park et al^9^ and SI Part 1A. We generated animations of three types of flush displays, based on three avian flush-pursuers: forelimbs forward move (based on the greater roadrunner, *Geococcyx californianus*), forelimbs forward move with additional tail hitches (based on the rufous-tailed scrub robin, *Cercotrichas galactotes*), forelimbs upward and outward move in three hitches (based on the northern mockingbird, *Mimus polyglottos*). The details are provided in SI Part 1.

#### Experiment 1

This experiment involved animations (Video S3) imitating the display of the greater roadrunner, characterized by wing movements (Links 2 and 3) in SI Part 1C. This species either extended its wings to the side or lifted and twisted during a flush display. Considering that experiment 2 focused on a sideways display, we chose to imitate the lift-twist case. The animation details are described in SI Part 1D.

#### Experiment 2

This experiment involved animations (Video S4) imitating the display of the rufous-tailed scrub robin, which spreads its wings sideways and moves its tail vertically in a flush display (Link 4 in SI Part 1C). The animation details are described in SI Part 1E.

#### Experiment 3

This experiment involved animations (Video S5) imitating the northern mockingbird’s display characterized by wing hitches (Links 5–10 in SI Part 1C, for example: 5, 6, 7). As displays with one to three hitches show similar frequencies^27^, we aimed to optimize foraging efficiency by imitating a display with three hitches. The animation details are given in SI Part 1F.

#### Experiment 4

We assessed the effect of dinosaur walking just before the flush display in the dinosaur animations (Video S6), which were based on the greater roadrunner’s display (as in *Experiment 1*). The animation details are described in SI Part 1G.

### General experimental design

In experiments 1–3, we generated four animations using combinations of feather presence (with or without) and distance (relatively close or distant). The animations simulated distances of 60 (“close”) and 120 cm (“distant”) between the prey and the virtual dinosaur. In Experiment 4, we focused on the presence or absence of walking, rather than considering two distances. Similarly, animations were generated for each walking option, both with feathers and without feathers, resulting in a total of four animations.

For each of the four experiments, the four animations were each presented to six female farm-bred migratory locusts (*Locusta migratoria*). Each animation was repeated four times, resulting in 16 recordings per individual and 96 recordings per experiment. The order of the animations was randomized for each individual to mitigate the potential influence of previously presented animations on subsequent ones. A 1-minute pause was given between animations. Further details are provided in SI Part 1H.

### Stimulus properties

To assess the links between proto-wing and tail movements and neural responses, we considered several variables. For each frame of the dinosaur animations, we determined the angular size (unit: degree) between the tips of the left and right forelimbs, as well as the left and right edges of the tail, from the prey’s ground-level view. From these measurements, we calculated the angular velocity (deg/ms) based on frame-by-frame changes in angular size. Using MATLAB (version R 2022a, MathWorks, USA), we counted the number of black pixels in each frame of the dinosaur animations and estimated the change rate of black surface area (no. of pixels/ms). For each pair of consecutive frames, we also calculated the change rate in the number of pixels transitioning from black to white and white to black (no. of pixels/ms). The latter variable represents the movement of edges and/or changes in total edge length. Additionally, for each pair of consecutive rates, we calculated the acceleration in the change rate of the number of pixels with a white-to-black color transition (no. of pixels/ms). For the acceleration values, we used a logarithmic scale as suggested in the previous study^36^. We also calculated the relative speed of pixel color transitions from white to black by dividing the number of white pixels transitioning to black by the number of black pixels in the previous frame. Subsequently, we calculated the relative acceleration for each pair of consecutive relative color transition speeds. The MATLAB code is available in SI Part 1I.

### Extracellular recordings

We fixed the locust on a wooden holder, ventral side up. We carefully tilted the head backward using a pin to expose the neck. Beeswax was applied to both sides of the neck as a barrier to prevent the saline solution from escaping. The right eye was covered with beeswax to block visual input, while the left eye faced the monitor screen. Using established methods^9^ (also described in SI Part 1J), we recorded spiking activity in the right DCMD.

### Data analysis

We used Spike2 software (version 5, Cambridge Electronic Design, Cambridge) to analyze the neural spike data (SI Part 1J). In each neural recording, we observed several peaks in the spiking rate corresponding to the movements of the animated dinosaur. Therefore, the number of peaks depended on the type of animation used. For each peak, we measured its size (no. of spikes/20 ms) and determined whether it reached five spikes per 20 ms, the estimated hypothetical value when, on average, hind leg muscle co-contraction occurs in free-moving locusts^41^. Muscle co-contraction is the initial stage of jump preparation.

We ran a generalized linear mixed-effects model to examine the effect of feather presence and distance on peak size (continuous variable determined for each animation phase; no. spikes/20 ms) and the occurrence of reaching the hypothetical threshold for initiating jump preparation (binary variable determined for each animation phase; reached = 1, unreached = 0). The fixed effects included feather (binary variable: with or without), distance (binary variable: close or distant), and an interaction term (feather * distance). Display phase (3 or 4 levels, depending on the animation type) was also included as a fixed factor. When analyzing the data from experiment 4, walking (binary variable: walk or stationary) was used as a fixed effect (the experiment imitated only one distance). Individual ID and record ID nested within individual ID were used as random effects. As peak size is an under-dispersed count, we used the Conway-Maxwell Poisson family with the log link function, employing the *glmmTMB* function from the *glmmTMB* package^65^. Since the variable ‘threshold’ is binary, we assumed a Binomial distribution with the logit link function, using the *glmer* function from the *lme4* package^66^. To assess over- or under-dispersion, we applied the *testDispersion* function from the *DHARMa* package^67^ and found no significant concerns. All statistical analyses were conducted in *R* version 4.2.2^68^.

In experiment 1 with animations imitating the display of the greater roadrunner, no records reached the threshold value of 5 spikes per 20 ms in response to the ‘folding’ display phase (Table S5). Consequently, these data points were omitted from the analysis of the ‘threshold’ variable. For experiment 4, with animations imitating the display and walking of the greater roadrunner, in response to the ‘raising’ display phase, only three records reached the threshold (5 spikes/20 ms), all of which responded to animations with feathers and stationary treatments (Table S8). Moreover, none of the responses to walking and non-feathered animations reached the threshold in the ‘folding’ display phase (Table S8). Consequently, when analyzing the occurrence of reaching the hypothetical threshold for initiating jump preparation, we only considered records responding to the animation of the ‘stretching’ display phase without walking.

### Data availability

Authors can confirm that all relevant data are included in the paper and/or its supplementary information files.

## Acknowledgement

This work was supported by National Research Foundation of Korea grants (2019R1A6A1A10073437 to Y.-N. Lee, RS-2023-00247087 to S.-i. Lee, RS-2024-00343461 to W. Kim, and RS-2025-00514508 to J. Park). This work is a part of Jinseok Park’s doctoral dissertation at the School of Biological Sciences, Seoul National University.

## Author contributions

Y.L., S.L., and P.G.J. conceptualized the project and supervised the overall experiments. J.P. and P.G.J. developed the methodology. J.P. conducted animation production, neurophysiological experiments, data curation and analysis with W.K., and visualization, and drafted the manuscript with M.S. J.P., W.K., Y.L., and S.L. acquired funding. All authors reviewed and commented on the manuscript.

## Extended Data

**Extended Data Fig. 1.**
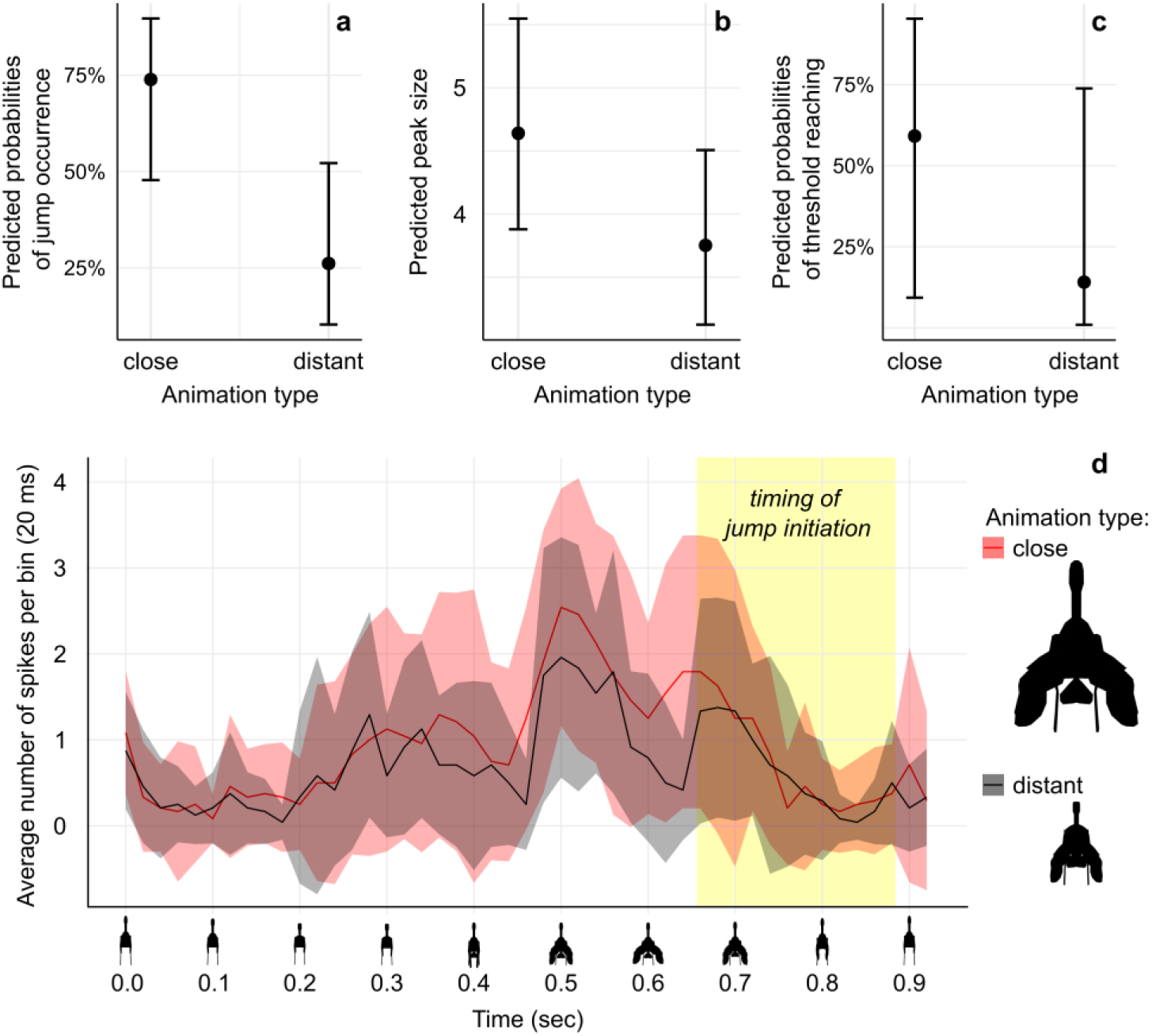
Results of the pilot experiment testing behavioral and neural responses in the same individual locusts. **a**, Probability of jump occurrence, **b**, Predicted size of peak spiking rate (no. of spikes per 20 ms), and **c**, Probability of reaching the threshold (5 spikes/20 ms) for jump preparation^41^ in response to a hypothetical flushing dinosaur animation imitating close and distant displays. Dots indicate predicted values, and vertical lines represent 95% confidence intervals from the statistical model presented in Table S4. **d**, Spiking rate of the locust’s LGMD/DCMD escape pathway (mean ± SD; n = 24, i.e., four recordings from each of six individuals) in response to animations imitating close (red) and distant (black) scenarios in the pilot experiments. Time (sec) is plotted on the X-axis, with corresponding screenshots below. In the behavioral experiment using the same animals and animations, most jumps were initiated during the animatic dinosaur folding its forelimbs, and the latency to jump initiation ranged from 0.656 to 0.884 sec (yellow box). The animation can be viewed in Video S2.

**Extended Data Fig. 2.**
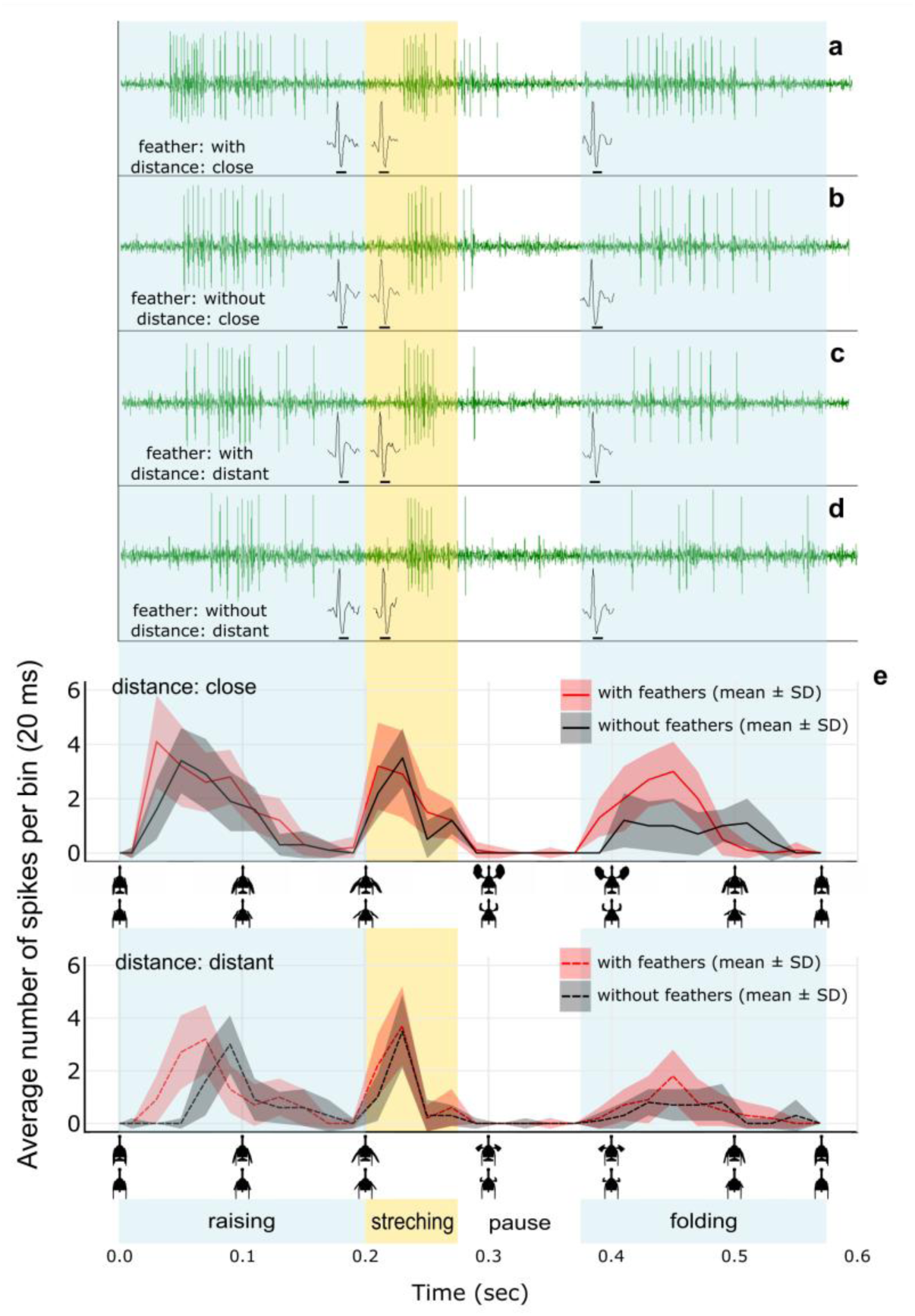
Examples of raw neural recordings in response to animations imitating the display of *Geococcyx californianus* in Experiment 1. “with” (**a**, **d**) or “without” (**b**, **d**) feathers at either “close” (**a**, **b**) or “distant” (**c**, **d**) location. Single spike shapes within each colored box represent an example from the recording corresponding to that display phase. The four recordings in a–d are from the same individual. **e**, Average spiking rate of the locust’s LGMD/DCMD escape pathway (no. spikes/20 ms; mean ± SD; n = 24, i.e., four recordings from each of six individuals) in response to animations “with” (red) and “without” feathers (black). The upper panel in E represents the imitation of a “close” situation (60 cm to prey), while the lower panel represents a “distant” case (120 cm to prey). Time (sec) is plotted on the X-axis, with corresponding screenshots of animations below the X-axis. Display phases are indicated by color-shaded semitransparent vertical boxes.

**Extended Data Fig. 3.**
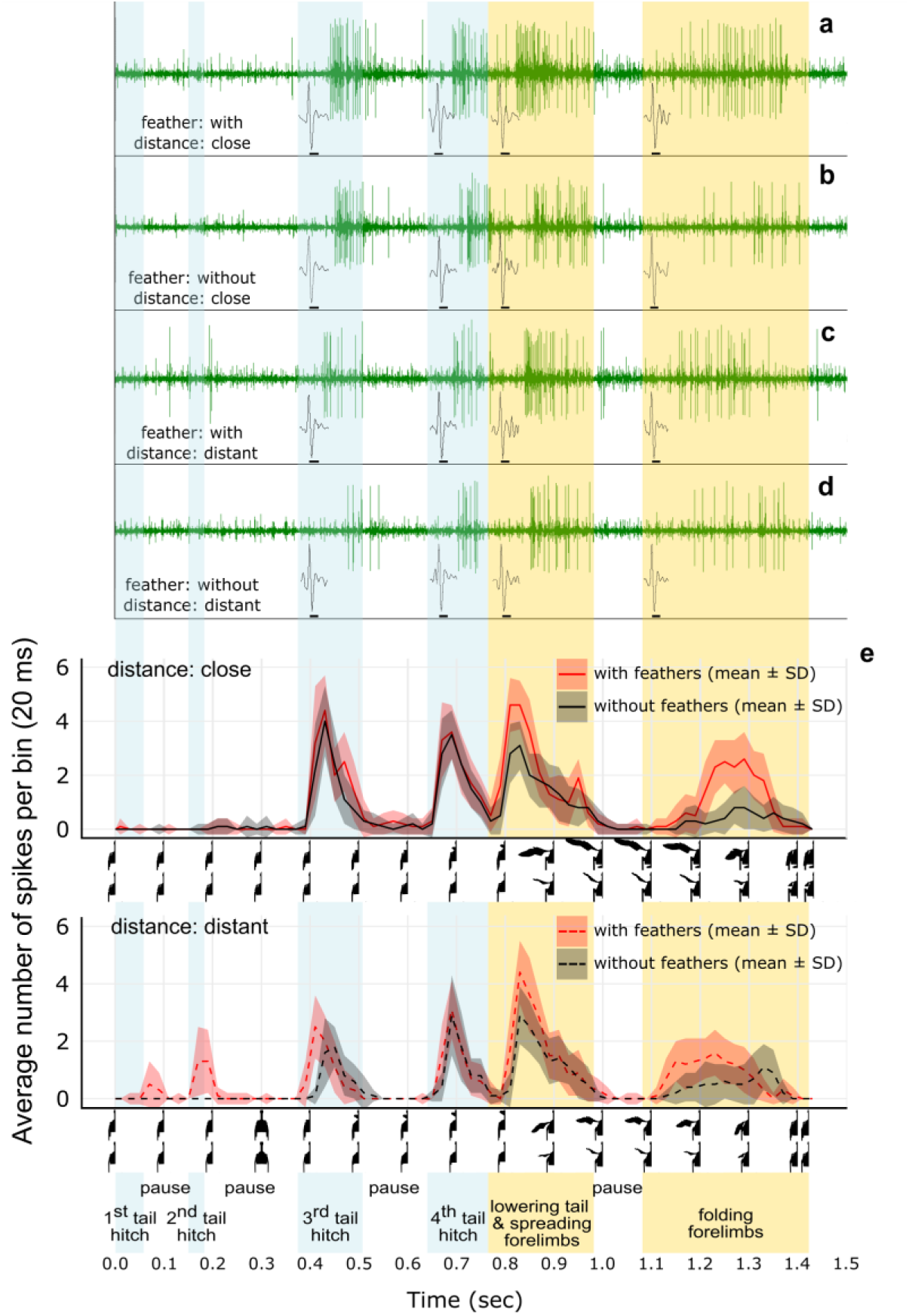
Examples of raw neural recordings in response to animations imitating the display of *Cercotrichas galactotes* in Experiment 2. “with” (**a**, **d**) or “without” (**b**, **d**) feathers at either “close” (**a**, **b**) or “distant” (**c**, **d**) location. Single spike shapes within each colored box represent an example from the recording corresponding to that display phase. The four recordings in a–d are from the same individual. **e**, Average spiking rate of the locust’s LGMD/DCMD escape pathway (no. spikes/20 ms; mean ± SD; n = 24, i.e., four recordings from each of six individuals) in response to animations “with” (red) and “without” feathers (black). The upper panel in E represents the imitation of a “close” situation (60 cm to prey), while the lower panel represents a “distant” case (120 cm to prey). Time (sec) is plotted on the X-axis, with corresponding half-screenshots of animations below the X-axis. Display phases are indicated by color-shaded semitransparent vertical boxes.

**Extended Data Fig. 4.**
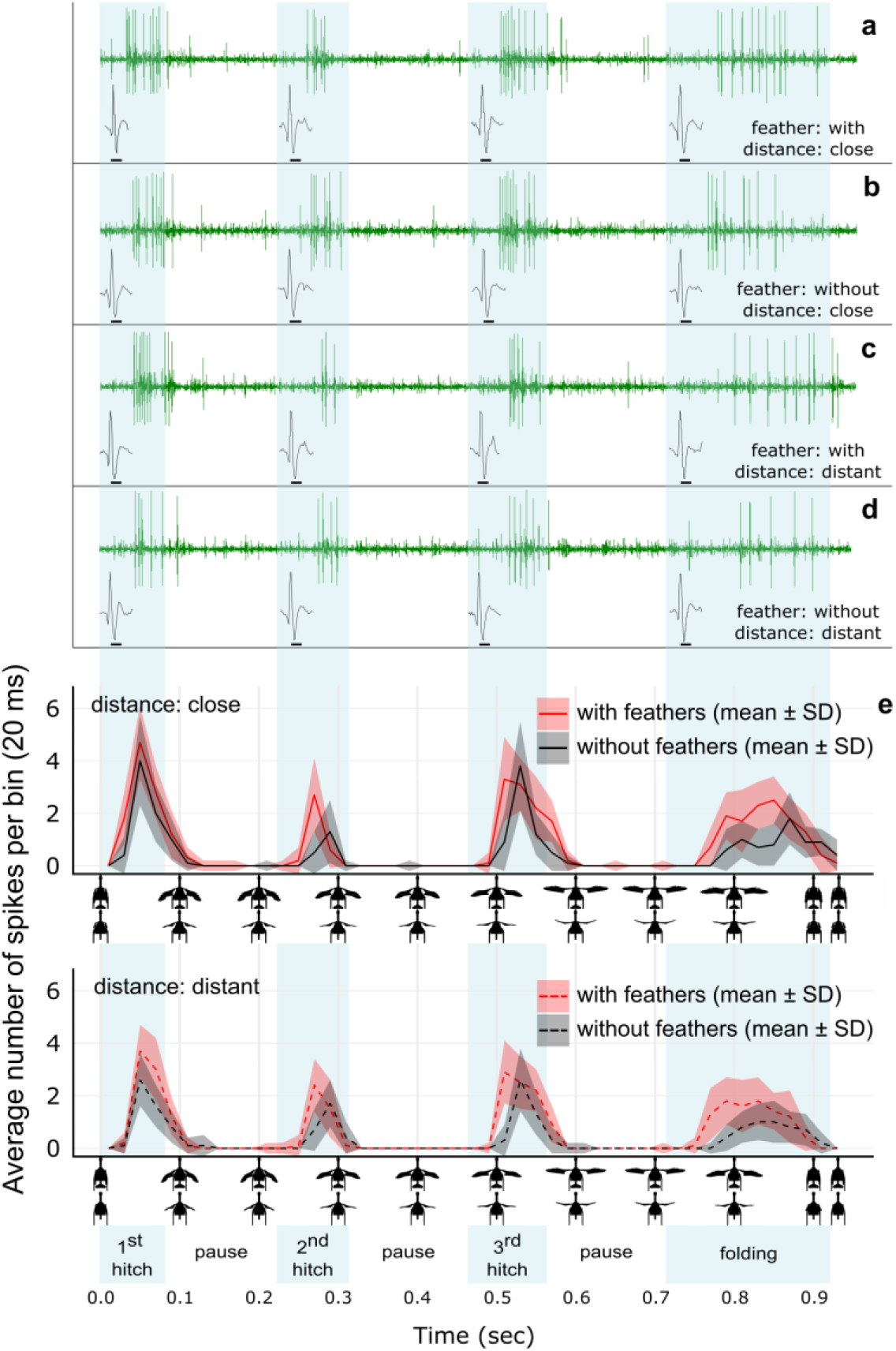
Examples of raw neural recordings in response to animations imitating the display of *Mimus polyglottos* in Experiment 3. “with” (**a**, **d**) or “without” (**b**, **d**) feathers at either “close” (**a**, **b**) or “distant” (**c**, **d**) location. Single spike shapes within each colored box represent an example from the recording corresponding to that display phase. The four recordings in a–d are from the same individual. **e**, Average spiking rate of the locust’s LGMD/DCMD escape pathway (no. spikes/20 ms; mean ± SD; n = 24, i.e., four recordings from each of six individuals) in response to animations “with” (red) and “without” feathers (black). The upper panel in E represents the imitation of a “close” situation (60 cm to prey), while the lower panel represents a “distant” case (120 cm to prey). Time (sec) is plotted on the X-axis, with corresponding half-screenshots of animations below the X-axis. Display phases are indicated by color-shaded semitransparent vertical boxes.

**Extended Data Fig. 5.**
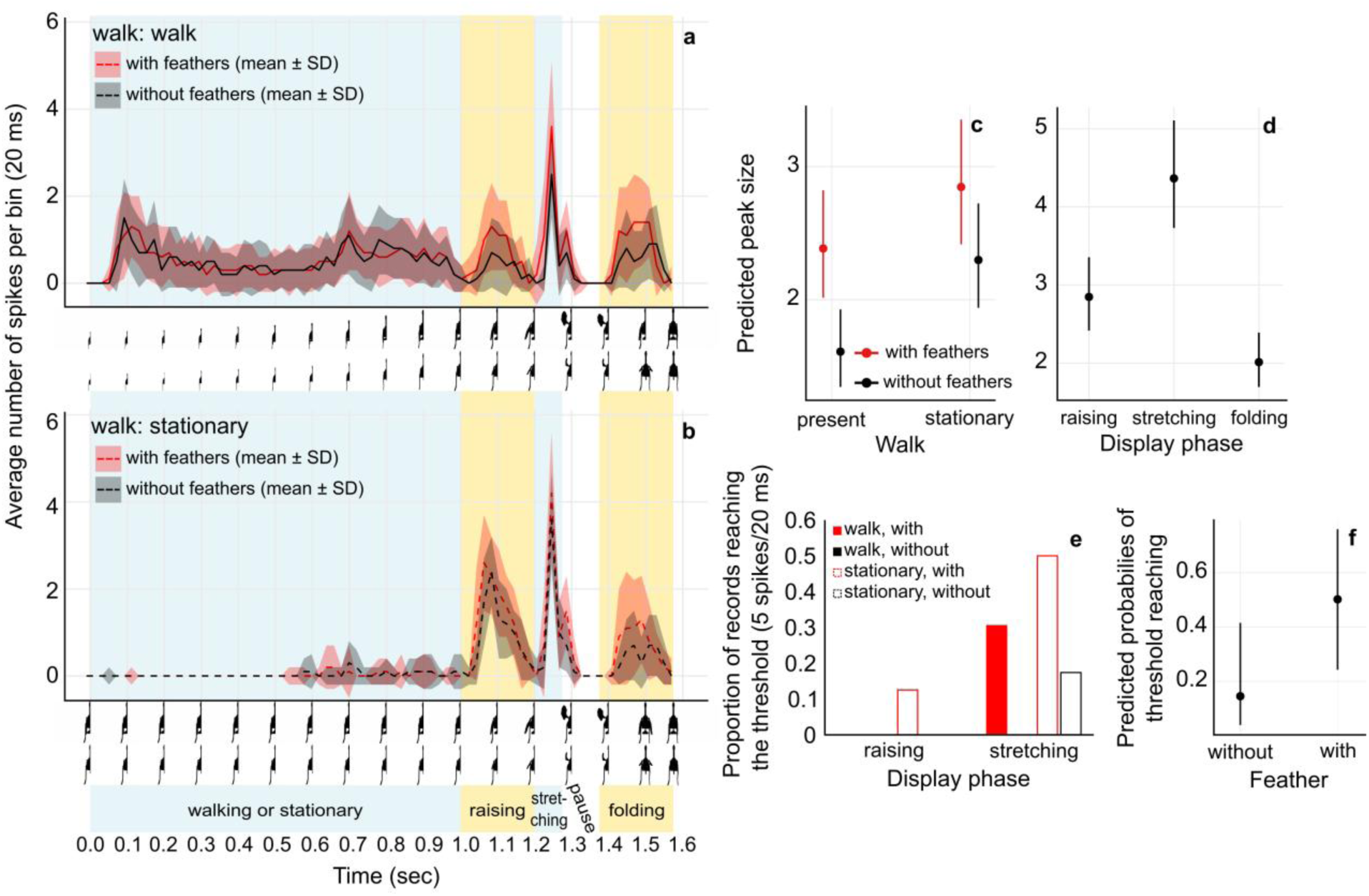
Results of the experiment involving animations based on the wing display of *Geococcyx californianus* and a walking stimulus of 1 m/sec in Experiment 4. **a**, **b**, Spiking rate of the locust’s LGMD/DCMD escape pathway (no. spikes/20 ms; mean ± SD; n = 24, i.e., four recordings from each of six individuals) in response to animations with (red) and without feathers (black). Panel A represents the case where walking stimulation is included, while Panel b represents the case where the dinosaur model remains stationary until it begins the forelimb display. Time (sec) is plotted on the X-axis, with corresponding half-screenshots below. Display phases are indicated by semitransparent vertical boxes shaded in blue for walking or stationary and stretching phases and yellow for raising and folding forelimb displays. **c**, Predicted size of peak spiking rate (no. spikes/20 ms^41^) for jump preparation in response to walking (left) and stationary (right) dinosaur animations, which begin their forelimb display with (red) and without (black) feathers. **d**, Predicted size of peak spiking rate (no. spikes/20 ms^41^) for jump preparation for each forelimb display phase: raising, stretching, and folding. **e**, Proportion of records (n = 24, i.e., four recordings from each of six individuals) reaching the threshold value (5 spikes/20 ms^41^) for jump preparation across different display phases (X-axis) and animation types. Animation type is differentiated by color (red – with feathers, black – without feathers) and box filling (filled – walk, empty – stationary). **f**, Probability of reaching the threshold (5 spikes/20 ms^41^) in response to dinosaur animation with (red) and without (black) feathers. Because most records did not reach the threshold (Table S8; see Methods), we only considered records responding to the ‘stretching’ phase of the stationary animation. In c, d, and f, dots indicate predicted values, while vertical lines represent 95% confidence intervals derived from the statistical model presented in Table S9. The animation can be viewed in Video S6.

**Extended Data Fig. 6.**
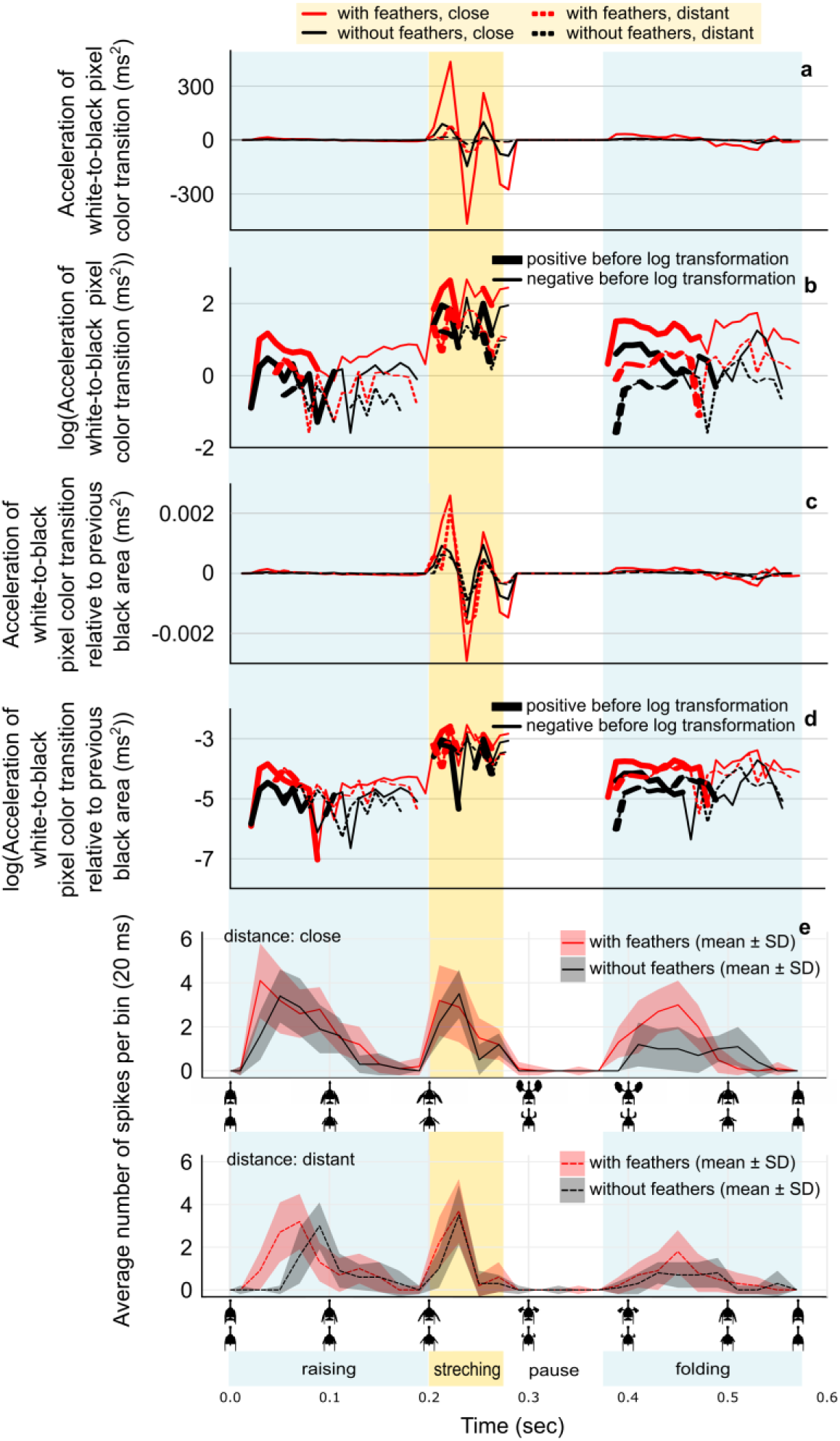
Profiles of the acceleration of white-to-black transition i.e., the change in the number pixels with color transition per pair of consecutive frames during animation (a–d), and the responses of DCMD (e) to those animations imitating the display of *Geococcyx californianus* in Experiment 1 by dinosaurs “with” (red line) or “without” (black line) feathers at “close” (solid line) or “distant” (dashed line) locations. **a**, Acceleration of the white-to-black color transition (no. pixels/ms^2^) based on Fig. S9C. **b**, same as (a) but with log-transformed Y-axis. **c**, “Relative Acceleration” of the white-to-black color transition represented as a proportion of the total black area (i.e., no. of white-to-black transition pixels for each pair of frames divided by the total number of black pixels in the first of the two frames). **d**, same as (c) but with log-transformed Y-axis. **e**, Average spiking rate per bin (no. spikes/20 ms; mean ± SD; n = 24, i.e., four recordings from each of six individuals) of the locust’s LGMD/DCMD escape pathway in response to animations “with” (red) and “without” feathers (black). Upper panel in E represents the imitation of a close situation (60 cm to prey), while lower panels represent a distant case (120 cm to prey). Time (sec) is plotted on the X-axis, with corresponding half-screenshots below. Display phases are indicated by color-shaded vertical boxes.

**Extended Data Fig. 7.**
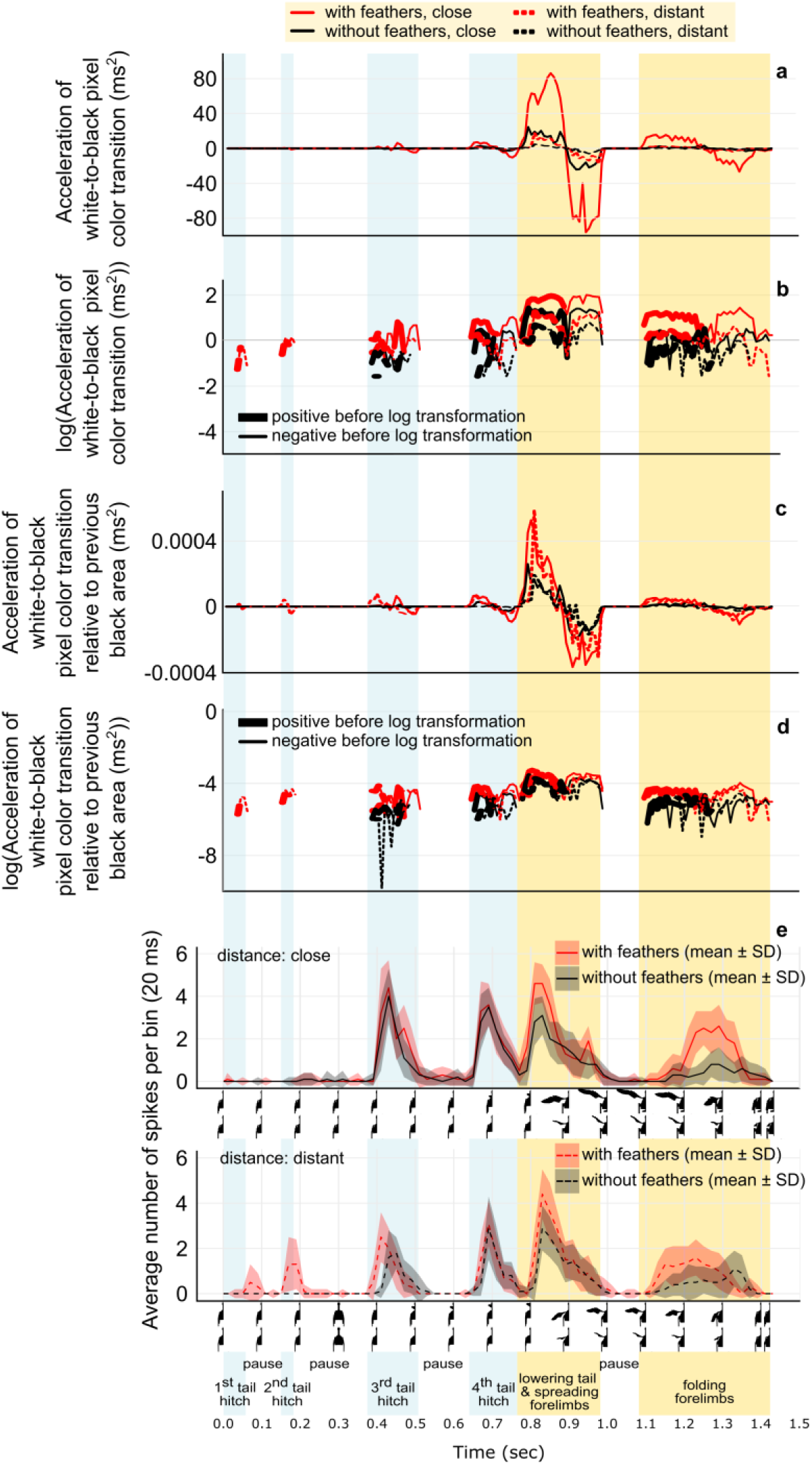
Profiles of the acceleration of white-to-black transition i.e., the change in the number pixels with color transitions per pair of consecutive frames during animation (a–d), and the responses of DCMD (e) to those animations imitating the display of *Cercotrichas galactotes* in Experiment 2 by dinosaurs “with” (red line) or “without” (black line) feathers at “close” (solid line) or “distant” (dashed line) locations. **a**, Acceleration of the white-to-black color transition (no. pixels/ms^2^) based on Fig. S10C. **b**, same as (a) but with log-transformed Y-axis. **c**, “Relative Acceleration” of the white-to-black color transition represented as a proportion of the total black area (i.e., no. of white-to-black transition pixels for each pair of frames divided by the total number of black pixels in the first of the two frames). **d**, same as (c) but with log-transformed Y-axis. **e**, Average spiking rate per bin (no. spikes/20 ms; mean ± SD; n = 24, i.e., four recordings from each of six individuals) of the locust’s LGMD/DCMD escape pathway in response to animations “with” (red) and “without” feathers (black). Upper panel in E represents the imitation of a close situation (60 cm to prey), while lower panels represent a distant case (120 cm to prey). Time (sec) is plotted on the X-axis, with corresponding half-screenshots below. Display phases are indicated by semitransparent vertical boxes.

**Extended Data Fig. 8.**
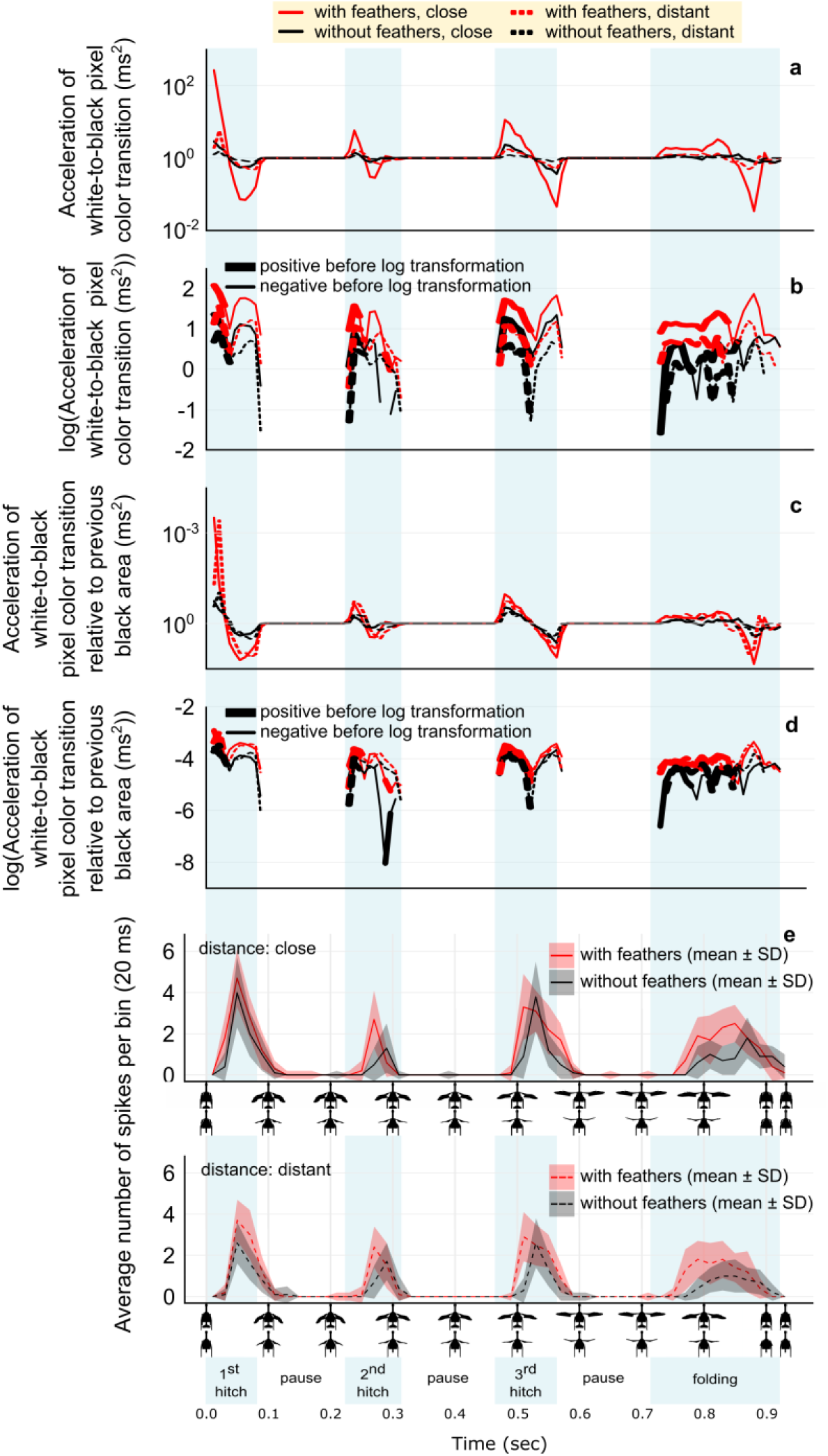
Profiles of the acceleration of white-to-black transition i.e., the change in the number pixels with color transition per pair of consecutive frames during animation (a–d), and the responses of DCMD (e) to those animations imitating the display of *Mimus polyglottos* in Experiment 3 by dinosaurs “with” (red line) or “without” (black line) feathers at “close” (solid line) or “distant” (dashed line) locations. **a**, Acceleration of the white-to-black color transition (no. pixels/ms^2^) based on Fig. S11C. **b**, same as (a) but with log-transformed Y-axis. **c**, “Relative Acceleration” of the white-to-black color transition represented as a proportion of the total black area (i.e., no. of white-to-black transition pixels for each pair of frames divided by the total number of black pixels in the first of the two frames). **d**, same as (c) but with log-transformed Y-axis. **e**, Average spiking rate per bin (no. spikes/20 ms; mean ± SD; n = 24, i.e., four recordings from each of six individuals) of the locust’s LGMD/DCMD escape pathway in response to animations “with” (red) and “without” feathers (black). Upper panel in E represents the imitation of a close situation (60 cm to prey), while lower panels represent a distant case (120 cm to prey). Time (sec) is plotted on the X-axis, with corresponding half-screenshots below. Display phases are indicated by semitransparent vertical boxes.

**Extended Data Fig. 9.**
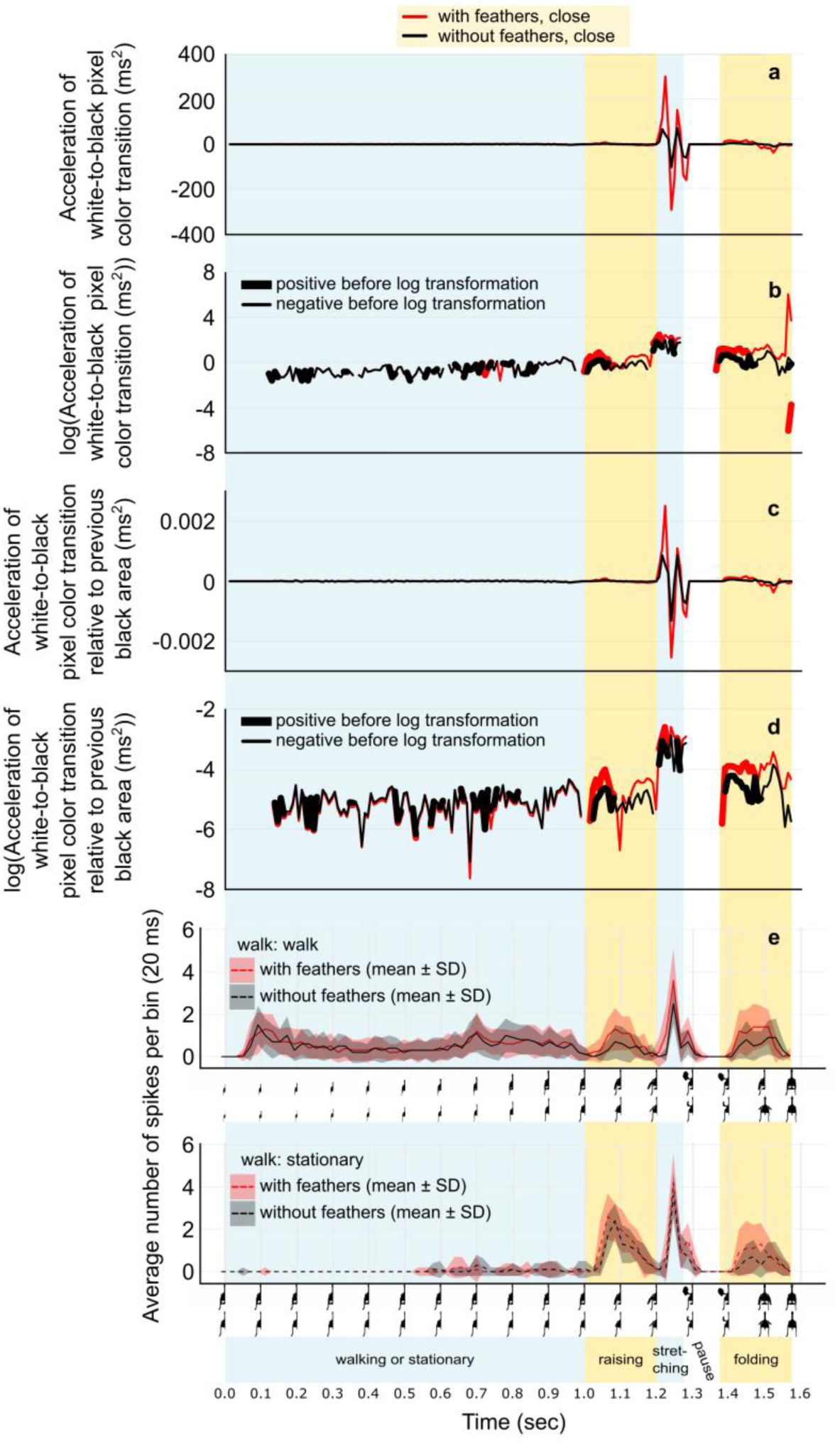
Profiles of the acceleration of white-to-black transition i.e., the change in the number pixels with color transition per pair of consecutive frames during animation (a–d), and the responses of DCMD (e) to the animations imitating the display of *Geococcyx californianus* in Experiment 4 by dinosaurs “with” (red line) or “without” (black line) feathers including a “walk” before display or presenting a “stationary” display. **a**, Acceleration of the white-to-black color transition (no. pixels/ms^2^) based on Fig. S15C. **b**, same as (a) but with log-transformed Y-axis. **c**, “Relative Acceleration” of the white-to-black color transition represented as a proportion of the total black area (i.e., nr of white-to-black transition pixels for each pair of frames divided by the total nr of black pixels in the first of the two frames). **d**, same as (c) but with log-transformed Y-axis. **e**, Average spiking rate per bin (no. spikes/20 ms; mean ± SD; n = 24, i.e., four recordings from each of six individuals) of the locust’s LGMD/DCMD escape pathway in response to animations “with” (red) and “without” feathers (black). Upper panel in E represents the imitation with walking, while lower panel represents the stationary animation. Time (sec) is plotted on the X-axis, with corresponding half-screenshots below. Display phases are indicated by color-shaded vertical boxes.

