## Supplementary material for "Display functions of dinosaur proto-wings before powered flight"

Jinseok Park[1, 2], Minyoung Son[3], Woojoo Kim[1, 2], Yuong-Nam Lee[4*], Sang-im Lee[5*], and Piotr, G. Jablonski[6*]
*[1]Research Institute of Basic Sciences, Seoul National University; Seoul, South Korea; [2]School of Biological Sciences, Seoul National University; Seoul, South Korea; [3]Department of Earth and Environmental Sciences, University of Minnesota; Minneapolis, U.S.A.; [4]School of Earth and Environmental Sciences, Seoul National University; Seoul, South Korea; [5]Department of New Biology, DGIST; Daegu, South Korea; [6]Museum and Institute of Zoology, Polish Academy of Sciences; Warsaw, Poland*

**PART 1: Supplementary Methods**
1A) Angles in avian flush-displays and in animations considering pennaraptoran dinosaur anatomy (Fig. S1)
1B) Pilot experiments: Methods
1C) Analysis of videos of flush-pursuing birds for animation production
1D) Animations in Experiment 1: forelimb forward move
1E) Animations in Experiment 2: forelimb forward move & tail hitches
1F) Animations in Experiment 3: forelimb upward + outward in three hitches
1G) Animations in Experiment 4: forelimb forward-move preceded by walking
1H) Experimental set-up
1I) MATLAB codes for extracting variables from animations
1J) Extracellular recordings

**PART 2: Results of Pilot Experiments**
2A) Text
2B) Figure (Fig. S2)
2C) Tables (Tables S2–S4)

**PART 3: Results of the three main experiments**
3A) Profiles of average spiking rates for each individual (Figs. S3–S5)
3B) DCMD responses and profiles of angular size and angular velocity based on forelimb/tails tips in animations (Figs. S6–S8)
3C) DCMD response and profiles of visual expansion of dinosaur silhouette and edge movements (Figs. S9–S11)
3D) Text – commentary to Extended Data Figs. 6–9 and Figs S6–S11 and S14–S15

**PART 4: Results of Experiment 4 with walking before display** (Figs. S12–S15)

**PART 5: Supplementary Tables with results** (Tables S5–S9)

**PART 6: Details of visual display hypothesis and review of sauropsid visual displays**
6A) Hypothetical flush-displays in Pennaraptora
6B) Reinforcing natural selection mechanisms of the flush-display component of the visual display hypothesis
6C) Visual display hypothesis (Fig. S16)
6D) Brief overview of motion-based signals in Sauropsida (Table S10)
6E) Examples of various descriptions of display motions in Sauropsida, excluding birds.
6F) Examples of papers on Sauropsida with descriptions of visual displays
6G) Examples of terrestrial agonistic contests associated with visual-displays in lekking birds that also use of wings for non-display functions inextricably linked to the visual-display behavior

**PART 7: Description of Supplementary Videos**

**PART 8: Supplementary References**

### Part 1: Supplementary Methods

#### 1A) Angles in avian flush-displays and in animations considering pennaraptoran dinosaur anatomy

The reasons for choosing *Caudipteryx* fossils as the basis for the animations are the same as the reasons for the robot pennaraptoran in our earlier study[1] being based on *Caudipteryx* (See Methods in Park et al[1] for a full explanation). Briefly, the resting posture was based on Senter and Robins[2], and the possible motion ranges were inferred using the phylogenetic bracketing approach[3] based on the nearest species *Acrocanthosaurus*[4] and *Bambiraptor*[5] (see Table S8 in Part et al[1]).

Diet ecology of *Caudipteryx* model[1]: Finally, although *Caudipteryx* is often presented as herbivorous, it may have been omnivorous, as were the early basal pennaraptora[6]. The preservation of gastroliths in several *Caudipteryx* fossils[7,8] indicates inconclusively that the diet may have included hard plant materials[9]. However, gastroliths may also indicate a diet of arthropods and other prey with hard exoskeletons (as shown in extant lizards[10,11] and birds[12]), suggesting an omnivorous diet. Indeed, while the later derived oviraptorosaurs showed functional mandibular characters indicating specialization for herbivory that requires handling of particularly hard plant materials, these functional morphological characters are less pronounced in the caudipterid clade within Oviraptosauria[13], which is also consistent with omnivory rather than exclusive herbivory. Assuming omnivory, it is feasible to hypothesize that—similar to omnivorous flush-pursuing birds such as the greater roadrunner (*Geococcyx californianus*[14]) or the northern mockingbird (*Mimus polyglottos*[15])—these early basal omnivorous pennaraptorans may have incorporated flush-displays into their foraging methods, in addition to using their proto-wings and tails in other displays. Therefore, testing the efficacy of proto-wing and tail feathers in a hypothetical flush-display by a pennaraptoran based on *Caudipteryx* is, in our view, a reasonable approach to evaluating the role of these structures in enhancing motion-based flush-displays in early Pennaraptora.

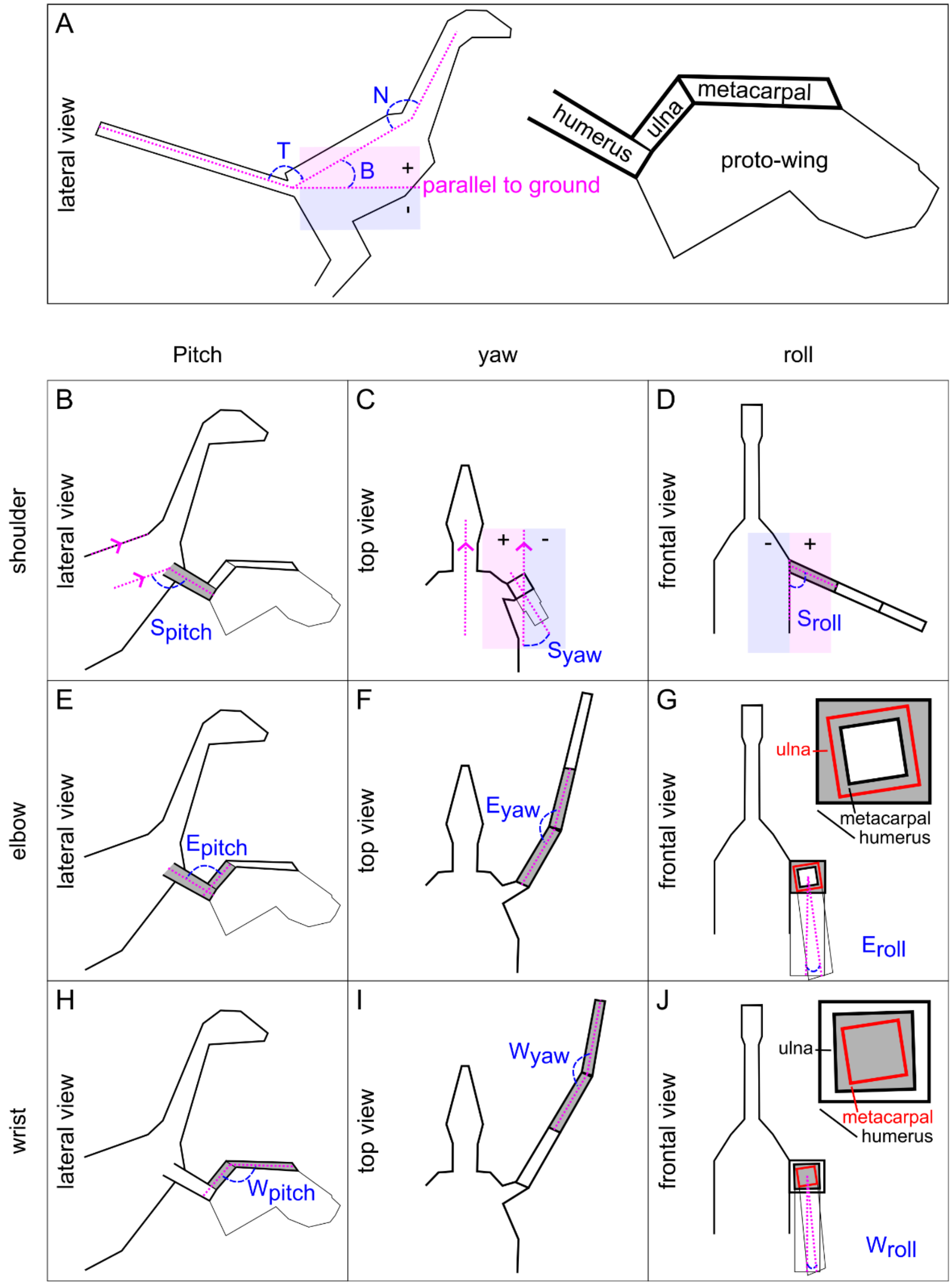


**Figure S1.** Definitions of angles used in building dinosaur animations

(A) Three angles in lateral view. ***Body angle*** (**B**) is defined as the angle between a line parallel to the horizontal ground and a line parallel to the dinosaur's back, extending from the base of the tail to the base of the neck. The angle value is positive when the base of the neck is higher than the base of the tail. The angle value is negative when the base of the neck is lower, indicating a downward bend. ***Neck angle*** (**N**) is defined as the angle between a straight line parallel to the back and a straight line following the contour of the neck, extending from the base of the neck to the top of the head. ***Tail angle*** (**T**) is defined as the angle between a straight line parallel to the back and a straight line following the contour of the tail, extending from the base to the tip of the tail.

Forelimb involves three sections corresponding to humerus, ulna, and metacarpal.

(B) Shoulder joint pitch (**$S_{pitch}$**) is defined in the lateral view as the angle between a straight line parallel to the back and the humerus.

(C) Shoulder joint yaw (**$S_{yaw}$**) is defined in the top view as the angle between a line parallel to the sagittal plane of the main body and the plane perpendicular to the humerus. Zero value represent humerus oriented vertically. Positive values represent humerus closer to the body and negative values represent humerus moved away from the body.

(D) Shoulder joint roll (**$S_{roll}$**) is defined in the frontal view as the angle between the side of the body (a line perpendicular to the ground and tangent to the body side) and the humerus. Positive values represent the humerus tip lifted up away from body; negative values indicate humerus positioned closer to the body axis.

(E) Elbow joint pitch (**$E_{pitch}$**) is defined in the later view as the angle between the humerus and the ulna.

(F) Elbow joint yaw (**$E_{yaw}$**) is defined in the top view as the angle between the humerus and the ulna.

(G) Elbow joint roll (**$E_{roll}$**) is defined as the angle between the planes of feather surfaces on the humerus and ulna, viewed in the frontal plane. This measures how much the ulna rotates along its axis relative to the unrotated position, where the feather attachment line on the ulna aligns with that on the humerus. Positive values indicate feather tips moving away from the body, while negative values indicate rotation toward the body.

(H) Wrist joint pitch (**$W_{pitch}$**) is defined in lateral view as the angle between the ulna and the metacarpal.

(I) Wrist joint yaw (**$W_{yaw}$**) is defined in the top view as the angle between the ulna and the metacarpal.

(J) Wrist joint roll (**$W_{roll}$**) is defined as the angle between the planes of feather surfaces on the ulna and metacarpal, viewed in the frontal plane. This measures how much the metacarpal rotates along its axis relative to the unrotated position, where the feather attachment line on the metacarpal aligns with that on the ulna. Positive values indicate feather tips moving away from the body, while negative values indicate rotation toward the body.

#### 1B) Pilot experiments: Methods

To confirm that the experimental setup targeted the DCMD neuron, we generated a circular looming stimulus previously used in numerous studies on orthopteran DCMD. The stimulus comprised a black approaching circle characterized by $l/|v| = 5$ ms, where l represents a radius of 3 cm and v is a constant approaching speed of 6 m/s, similar to stimuli commonly utilized in neurophysiological studies of the LGMD/DCMD pathway[16–20]. The animation simulated the approach from a starting point when the circle subtended 0.74° on the retina, to a final point when it subtended 56.48° of the field of view on the retina of the locust. We presented this animation to five female farm-bred migratory locusts and observed their behavioral responses 2–5 times, with 1-min intervals between observations. Additionally, for two individuals that reliably responded by jumping, we recorded their neural responses to the stimulus three times, with 1-min intervals between recordings, one day after the behavioral test.

To verify if relatively complex dinosaur animations indeed trigger escape behavior in locusts and to establish a link between behavioral and neural responses, we tested three male and three female locusts using a pilot flushing dinosaur animation with proto-wings and tail feathers. The animation simulated two scenarios: relatively close (70 cm) and distant (130 cm). These animations (Video S2) served as preliminary versions, which approximately mimicked the display of the rufous-tailed scrub robin (*Cercotrichas galactotes*; Link 4 in SI Part 1C), i.e., the angles at each animation step were visually matched to the video clip without precise measurements of the forelimb and tail angles (which was done in Exp. 2). Therefore, the pilot animation differs in small details from the more precise rufous-tailed scrub robin's animation in Experiment 2. The aim of the pilot experiment was to determine within an individual locust a link between the observed escape behavior and the recorded neural response (n = 6 individuals). The following angles characterize the pilot animation presented to the locusts:

Phase 0: The dinosaur model in a resting posture ($S_{pitch} = 33°$, $S_{roll} = 12°$, $E_{pitch} = 106°$, $W_{pitch} = 131°$, T = 160°, B = 0°, N = 147°).

Phase 1: Leaning of the main body downward and tail upward ending with T = 180°, B = -21° at the end of the phase 1 duration of 0.25 sec.

Phase 2: Lowering the tail and stretching the forelimbs forward ending up with $S_{pitch} = 65°$, $S_{yaw} = 7°$, $S_{roll} = -6°$, $E_{pitch} = 138°$, $W_{pitch} = 178°$. Phase 2 lasted 0.167 sec.

Phase 3: Moving the forelimbs to the sides, which results in $S_{pitch}$ = 107°, $S_{yaw}$ = 47°, $S_{roll}$ = 5°. The movements in phase 3 lasted for 0.133 sec, followed by maintaining the final posture for 0.1 sec.
Phase 4: Returning to Phase 0 during 0.25 sec.

First, we presented the animations to six freely moving individual locusts and recorded whether they jumped or not, with 1-min intervals between presentations. Each locust was tested four times for both close and distant treatments. Three locusts experienced the close-distant-close-distant order, while the other three experienced the distant-close-distant-close order. After one day, we recorded neural responses to the same animations, with 1-min intervals between observations. Each locust was tested four times for both close and distant treatments. Locusts that experienced the close-first order in the behavioral experiment were tested in the distant-first order in neural recordings, and vice versa. We then analyzed the effect of animation distance on jump behavior (using a *glmer* model with a binomial family), as well as on the peak spiking rate and the probability of reaching the hypothetical spiking rate threshold for initiating jump preparation (see the Methods 'Data analysis' section). Animation distance and sex were considered as fixed factors, while individual ID was included as a random effect.

#### 1C) Analysis of videos of flush-pursuing birds for animation production

We searched YouTube and the Macaulay Library media (Cornell Lab of Ornithology) for videos containing flush-pursue foraging of the greater roadrunner (*Geococcyx californianus*), rufous-tailed scrub robin (*Cercotrichas galactotes*), and northern mockingbird (*Mimus polyglottos*), and found a small number of videos: one for the robin, two for the roadrunner, and eleven for the mockingbird. We used the following videos to determine the timing and angles of forelimb and tail movements in the birds:

**Greater roadrunner**
Link 1) https://www.youtube.com/watch?v=qmXGDWkZSek © Kat Avila
Link 2) https://www.youtube.com/watch?v=ctm4PNNNo4M © 4GUESTS USA
Link 3; Video S1) https://www.youtube.com/watch?v=u6RSPk7y1us © 1986, licensed by Albatross World Sales

**Rufous-tailed scrub robin**
Link 4) https://www.youtube.com/watch?v=pS31OI4YNdw, © Canal Natura

**Northern mockingbird**
Link 5) https://macaulaylibrary.org/asset/200776531 © Joe Angseesing
Link 6) https://www.youtube.com/watch?v=fe1QXveO4aw © Colette Micallef
Link 7) https://www.youtube.com/watch?v=bMsa0T8fZpY © Linzy's Vids
Link 8) https://www.youtube.com/watch?v=OgX_BLkxe6I © Sport Cards Brass Trains VideoGames
Link 9) https://www.youtube.com/watch?v=znkKWFhMnac © Jim Emery
Link 10) https://macaulaylibrary.org/asset/200776491 © Joe Angseesing

We extracted the time durations of the flush display phases from these videos and took screenshots when the camera angle was nearly parallel to the bird's side view. The averaged values for time durations may not precisely align with the average value because the frames per second (fps) in the bird clip differ from those in dinosaur animations (120 fps). In such cases, we used 'based on' rather than 'averaged from' to indicate the distinction. The time unit was rounded to the fourth decimal place, and the angle unit was rounded to the first decimal place.

Using ImageJ (version 1.53e), we measured body angles, neck angles, and stride lengths from these screenshots. We then calculated stride speeds based on stride lengths and time durations. The number of video clips is small and may not fully represent the average flush display characteristics of a species. However, given the observed variation in flush behavior among bird species (see links above and Supplementary Information in Park et al.[1]), we believe that this approach is sufficient to assess the role of proto-wings and tail feathers across different flush displays observed in modern birds, and adjusted to the anatomy of the pennaraptoran dinosaurs.

#### 1D) Animations in Experiment 1: forelimb forward move

*Experiment 1. Imitating the display of the greater roadrunner*
The animations (Video S3) were based on two roadrunner clips (Links 1 and 2 in SI Part 1C). We found that this species either extended its wings to the side (n = 2; example in Link 1) or lifted and twisted its wings (n = 2; example in Link 2) when making a flush display. Since the robin experiment described below presents a sideway display, we adopted the twist case.

In Phase 0, the dinosaur model takes a resting posture[2]. Moving to Phase 1, the model raises its forelimbs. In Phase 2, the model stretches its forelimbs and involves two sub-phases. In Phase 1, the model rotates forward its forelimbs. Subsequently, in Phase 2, the model spreads more and twists its forelimbs, lasting 0.1 sec. Finally, in Phase 3, the model folds its forelimbs. Details are provided below:

Phase 0. Resting posture

- Forelimbs: the resting posture speculated by senter & robins[2] ($S_{pitch}$ = 33°, $S_{yaw}$ = 0°, $S_{roll}$ = 0°, $E_{pitch}$ = 106°, $E_{yaw}$ = 0°, $E_{roll}$ = 0°, $W_{pitch}$ = 131°, $W_{yaw}$ = 0°, $W_{roll}$ = 0°)
- Body: 23° (averaged from three cases: 14°, 22°, and 32°)
- Neck: 134° (averaged from three cases: 104°, 134°, and 164°)

Phase 1. Raising forelimbs for 0.2 sec (based on four cases: 0.267, 0.167, 0.167, and 0.133 sec).

- Forelimbs: The dinosaur model displays a slight shoulder joint elevation ($S_{roll}$ = 20°). The roadrunner spreads its primary feathers and alulae, whereas the resting posture of *Caudipteryx* already exhibits spread feathers. The roadrunner slightly rotates the shoulder forward; however, the shoulder is already rotated in the resting posture of *Caudipteryx*. The roadrunner slightly raises the shoulder, but the extent of the elevation is uncertain. Consequently, we have adjusted the dinosaur model by slightly raising its shoulders.

Phase 2. Stretching forelimbs
Phase 2-1. Rotating forelimbs forward for 0.042 sec (averaged from four cases: 0.033, 0.033, 0.033, and 0.067 sec).

- Forelimbs: The roadrunner spreads its forelimbs and rotates the shoulders forward ($S_{pitch}$ = 113°, $S_{yaw}$ = -11°, $S_{roll}$ = 41°, $E_{pitch}$ = 126°, $E_{yaw}$ = 5°, $E_{roll}$ = 16°, $W_{pitch}$ = 164°, $W_{yaw}$ = 28°, $W_{roll}$ = -15°). Here, we attempted the first display of link 1 (between 0–2 sec).

Phase 2-2. Spreading and twisting forelimbs for 0.033 sec (averaged from four cases: 0.033, 0.033, 0.033, and 0.033 sec)

- Forelimbs: The forelimbs are spread more and twisted in a manner that the upper parts rotate inward. We have attempted to imitate a roadrunner's posture ($S_{pitch}$ = 123°, $S_{yaw}$ = 29°, $S_{roll}$ = 67°, $E_{pitch}$ = 106°, $E_{yaw}$ = 0°, $E_{roll}$ = 0°, $W_{pitch}$ = 131°, $W_{yaw}$ = 12°, $W_{roll}$ = -2°). Here, we attempted the first display of link 1 (between 0–2 sec).

The posture lasts for 0.1 sec.

Phase 3. Folding forelimbs for 0.2 sec (averaged from four cases: 0.2, 0.2, 0.2, and 0.2 sec).

- Forelimbs: the resting posture speculated by senter & robins[2] ($S_{pitch}$ = 33°, $S_{yaw}$ = 0°, $S_{roll}$ = 0°, $E_{pitch}$ = 106°, $E_{yaw}$ = 0°, $E_{roll}$ = 0°, $W_{pitch}$ = 131°, $W_{yaw}$ = 0°, $W_{roll}$ = 0°).

#### 1E) Animations in Experiment 2: forelimb forward move & tail hitches

*Experiment 2. Imitating the display of the rufous-tailed scrub robin*
This species uses both its wings and tail in a flush display (Link 4 in SI Part 1C).

In the initial phase (phase 0) of animation (Video S4), the dinosaur model takes a resting posture[2]. Moving on to the next phase (Phase 1–4), the model raises its tail with four hitches, and the body tilts downwards: First hitch (Phase 1) with a pause of 0.092 sec, second hitch (Phase 2) with a pause of 0.192 sec, third hitch (Phase 3) and with a pause of 0.133 sec, and fourth hitch (Phase 4). In the fifth phase (Phase 5), the model lowers its tail and spreads its forelimbs, maintaining this posture for 0.1 sec. Finally (Phase 6), the model folds its forelimbs. Details are provided below:

Phase 0. Resting posture
- Forelimbs: the resting posture speculated by senter & robins[2] ($S_{pitch}$ = 33°, $S_{yaw}$ = 0°, $S_{roll}$ = 0°, $E_{pitch}$ = 106°, $E_{yaw}$ = 0°, $E_{roll}$ = 0°, $W_{pitch}$ = 131°, $W_{yaw}$ = 0°, $W_{roll}$ = 0°)
- Tail (T): 106° (averaged from four cases: 96°, 96°, 115°, and 118°)
- Body (B): 33° (averaged from two cases: 27° and 38°)
- Neck (N): 155° (averaged from two cases: 152°, 152°, 156°, and 158°)

Phase 1 to 4. Raising the tail, consisting of four hitches and three pauses between hitches. The robin exhibits variations in the body angle at the end of this phase, including tilting slightly upward, downward, and parallel to the ground. The parallel case appears relatively frequently in the bird clip we adopt (3 out of 6 cases). Therefore, we focused on the parallel case. In the animation, changes in the body angle (from 33° to 0°) are distributed based on the duration of each hitch.

Phase 1. First tail hitch for 0.058 sec (based on three cases: 0.033, 0.067, and 0.067 sec)
- Tail (T): 113° (up 7°, averaged from three cases: 3°, 7°, and 10°)
- Body (B): 28° (down 5°)

The posture lasts for 0.092 sec (based on three cases: 0.033, 0.033, and 0.2 sec)

Phase 2. Second tail hitch for 0.033 sec (averaged from three cases: 0.033, 0.033, and 0.033 sec)
- Tail (T): 120° (up 7°, averaged from three cases: 1°, 7°, and 11°)
- Body (B): 25° (down 3°)

The posture lasts for 0.192 sec (based on three cases: 0.167, 0.167, and 0.233 sec)

Phase 3. Third tail hitch for 0.133 sec (based on three cases: 0.1, 0.133, and 0.167 sec)
- Tail (T): 137° (up 17°, averaged from three cases: 11°, 15°, and 25°)
- Body (B): 12° (down 13°)

The posture lasts for 0.133 sec (based on three cases: 0.1, 0.132, and 0.167 sec)

Phase 4. Fourth tail hitch for 0.125 sec (based on three cases: 0.1, 0.1, and 0.167 sec)
- Tail (T): 157° (up 20°, averaged from three cases: 18°, 18°, and 24°)
- Body (B): 0° (down 12°)

Phase 5. Lowering the tail and spreading forelimbs for 0.217 sec (averaged from six cases: 0.133, 0.167, 0.2, 0.217, 0.233, 0.233, and 0.333 sec)
- Forelimbs: Spread them sidewise ($S_{pitch}$ = 122°, $S_{yaw}$ = -15°, $S_{yaw}$ = 81°, $E_{pitch}$ = 136°, $E_{yaw}$ = 0°, $E_{roll}$ = 0°, $W_{pitch}$ = 134°, $W_{yaw}$ = 2°, $W_{roll}$ = 3°). Due to the wing bones being covered with feathers, it is challenging to estimate the angles accurately. Therefore, we attempted to imitate the bird's posture from a viewpoint similar to that in the bird video. The angles are within the possible motion range of *Caudipteryx* (see Table S8 in Part et al[1]).

- Tail (T): 141° (estimated from a case; among the cases where the bird's side view is parallel to the camera in the clip, we chose a case in which the tail is lowered to ensure visibility to the prey, which is located on the ground.)
- Body (B): 47° (averaged from two cases: 35° and 58°)
- Neck (N): 130° (averaged from two cases: 129° and 130°)

The posture lasts for 0.1 sec.

Phase 6. Folding forelimbs for 0.342 sec (based on six cases: 0.433, 0.267, 0.2, 0.367, 0.633, and 0.167 sec).

- Forelimbs: the resting posture speculated by senter & robins[2] ($S_{pitch}$ = 33°, $S_{yaw}$ = 0°, $S_{roll}$ = 0°, $E_{pitch}$ = 106°, $E_{yaw}$ = 0°, $E_{roll}$ = 0°, $W_{pitch}$ = 131°, $W_{yaw}$ = 0°, $W_{roll}$ = 0°).

#### 1F) Animations in Experiment 3: forelimb upward + outward in three hitches

*Experiment 3. Imitating the display of the northern mockingbird*

This species uses its wings in a flush display. This flush display involves wing hitches, ranging from 1 to 5 hitches[21]. The frequency of displays featuring one to three hitches appears similar[21]. Given that pauses in a display can induce multiple spiking rate peaks, increasing the frequency of prey escape[22], a display with three hitches is expected to be more efficient for foraging than displays with one or two hitches. Therefore, we decided to imitate the display with three hitches. The duration of each phase is calculated based on 34 flush displays (Table S1; Links 5 and 10 in SI Part 1C).

**Table S1. Duration of hitches and pauses in clips involving the *Mimus polyglottos* display with three hitches in Experiment 3.** The first column shows the Clip ID and Display ID, while the remaining columns indicate the time duration (unit: sec) for each hitch and pause. The two bottom-most rows present the average and standard deviation (SD) values. Clip IDs 'links 5–10' are presented in SI Part 1C.

| Clip ID_Display ID | Duration (sec) | | | | | | |
|---|---|---|---|---|---|---|---|
| | Hitch 1 | Pause 1 | Hitch 2 | Pause 2 | Hitch 3 | Pause 3 | Folding |
| mockingbird 01_3 | 0.08 | 0.12 | 0.12 | 0.12 | 0.12 | 0.2 | 0.28 |
| mockingbird 01_4 | 0.16 | 0.08 | 0.12 | 0.12 | 0.12 | 0 | 0.28 |
| mockingbird 01_5 | 0.16 | 0.12 | 0.12 | 0.12 | 0.12 | 0.16 | 0.24 |
| mockingbird 01_7 | 0.16 | 0.08 | 0.12 | 0.12 | 0.12 | 0.16 | 0.2 |
| mockingbird 01_8 | 0.16 | 0.08 | 0.12 | 0.12 | 0.08 | 0.28 | 0.24 |
| DSCF1729Converted.avi_1 | 0.067 | 0.133 | 0.067 | 0.067 | 0.1 | 0.133 | 0.167 |
| DSCF1736Converted.avi_1 | 0.033 | 0.167 | 0.067 | 0.167 | 0.067 | 0.167 | 0.133 |
| DSCF1736Converted.avi_4 | 0.067 | 0.133 | 0.067 | 0.167 | 0.067 | 0.133 | 0.167 |
| DSCF1735Converted.avi_1 | 0.1 | 0.133 | 0.067 | 0.167 | 0.1 | 0 | 0.167 |
| DSCF1735Converted.avi_4 | 0.033 | 0.167 | 0.033 | 0.167 | 0.1 | 0.4 | 0.2 |
| DSCF1735Converted.avi_5 | 0.033 | 0.167 | 0.033 | 0.167 | 0.033 | 0.2 | 0.2 |
| DSCF1735Converted.avi_6 | 0.1 | 0.2 | 0.033 | 0.167 | 0.033 | 0.2 | 0.3 |
| DVD 322.avi_5 | 0.241 | 0.069 | 0.103 | 0.172 | 0.103 | 0 | 0.207 |
| Link 5_3 | 0.03 | 0.07 | 0.03 | 0.1 | 0.24 | 0.21 | 0.21 |
| Link 5_4 | 0.03 | 0.14 | 0.07 | 0.21 | 0.17 | 0.14 | 0.1 |
| Link 5_8 | 0.07 | 0.17 | 0.07 | 0.1 | 0.14 | 0.1 | 0.17 |
| Link 5_10 | 0.07 | 0.1 | 0.14 | 0.14 | 0.1 | 0.14 | 0.17 |
| Link 5_11 | 0.07 | 0.14 | 0.07 | 0.14 | 0.1 | 0.1 | 0.14 |
| Link 5_12 | 0.07 | 0.12 | 0.07 | 0.1 | 0.07 | 0 | 0.17 |
| Link 5_16 | 0.07 | 0.17 | 0.07 | 0.17 | 0.07 | 0.17 | 0.14 |
| Link 6_2 | 0.1 | 0.13 | 0.1 | 0.1 | 0.13 | 0.1 | 0.23 |
| Link 6_3 | 0.07 | 0.17 | 0.17 | 0.07 | 0.1 | 0.1 | 0.3 |
| Link 6_4 | 0.07 | 0.13 | 0.13 | 0.17 | 0.07 | 0.03 | 0.43 |
| Link 6_5 | 0.03 | 0.17 | 0.07 | 0.17 | 0.13 | 0.13 | 0.17 |
| Link 7_32 | 0.03 | 0.1 | 0.07 | 0.4 | 0.03 | 0.24 | 0.13 |
| Link 7_39 | 0.07 | 0.13 | 0.07 | 0.17 | 0.03 | 0.03 | 0.17 |
| Link 7_43 | 0.07 | 0.1 | 0.1 | 0.17 | 0.07 | 0.1 | 0.1 |
| Link 8_1 | 0.03 | 0.1 | 0.2 | 0.1 | 0.13 | 0.13 | 0.17 |
| Link 8_2 | 0.03 | 0.03 | 0.13 | 0.03 | 0.1 | 0.34 | 0.27 |
| Link 9_1 | 0.07 | 0.25 | 0.11 | 0.22 | 0.07 | 0.14 | 0.54 |
| Link 9_2 | 0.07 | 0.22 | 0.07 | 0.22 | 0.07 | 0.25 | 0.25 |
| Link 9_3 | 0.07 | 0.22 | 0.14 | 0.11 | 0.14 | 0.11 | 0.18 |
| Link 10_1 | 0.07 | 0.25 | 0.07 | 0.18 | 0.11 | 0.47 | 0.14 |
| Link 10_2 | 0.07 | 0.25 | 0.07 | 0.18 | 0.11 | 0.14 | 0.25 |
| Average | 0.08 | 0.14 | 0.09 | 0.15 | 0.1 | 0.15 | 0.21 |
| SD | 0.05 | 0.05 | 0.04 | 0.06 | 0.04 | 0.11 | 0.09 |

In Phase 0 of the animation (Video S5), the dinosaur model takes a resting posture[2]. Phase 1 involves lifting its forelimbs and a first pause. Phase 2 also includes lifting its forelimbs and a second pause. Phase 3 consists of stretching and spreading its forelimbs and a third pause. Finally, Phase 4 involves folding forelimbs to return to the resting posture. Details are provided below:

Phase 0. Resting posture

- Forelimbs: the resting posture speculated by senter & robins[2] ($S_{pitch}$ = 33°, $S_{yaw}$ = 0°, $S_{roll}$ = 0°, $E_{pitch}$ = 106°, $E_{yaw}$ = 0°, $E_{roll}$ = 0°, $W_{pitch}$ = 131°, $W_{yaw}$ = 0°, $W_{roll}$ = 0°)
- Body: 38° (averaged from seven cases: 10°, 22°, 37°, 37°, 38°, 56°, and 65°)

- Neck: 164° (averaged from seven cases: 152°, 152°, 158°, 161°, 172°, and 177°)

Phase 1. First hitch for 0.083 sec (based on 34 cases, Table S1)

- Forelimbs: The dinosaur model raises its forelimbs ($S_{roll}$ = 44°). The mockingbird lifts its wings nearly horizontally, with minimal feather spreading. However, three considerations were taken into account: 1) the mockingbird raises its wings further during the second hitch; 2) specific degrees of wing elevation for the first and second hitches cannot be estimated from bird clips; 3) *Caudipteryx* cannot raise its forelimbs to the horizontal level (max set to 88°). Consequently, the dinosaur model elevates its forelimbs to 44° for both the first and second hitches.

The first pause lasts for 0.141 sec (based on 34 cases, Table S1).

Phase 2. Second hitch for 0.091 sec (based on 34 cases, Table S1)

- Forelimbs: The dinosaur model raised and spread its forelimbs ($S_{roll}$ = 88°, $E_{pitch}$ = 121°, $W_{pitch}$ = 154.5°). The mockingbird exhibits diversity in this second hitch. In case A, the bird sometimes slightly spreads its feathers, similar to or less than the resting posture of *Caudipteryx*[2], without fully extending its wings in the subsequent motion (third hitch). Conversely, in case B, the bird appears to spread its feathers more than the resting posture of *Caudipteryx*[2], nearly fully extending its wings in the following motion (third hitch). Imitating the latter case (B) can offer valuable clarity in assessing the effect of feather presence. Considering the limitations of forelimb bones, the elbow joint can extend an additional 30°, and the wrist joint can spread by 47° more. To account for these constraints, we performed the second and third motions halfway.

The second pause lasts for 0.15 sec (averaged from 34 cases, Table S1).

Phase 3. Third hitch for 0.1 sec (averaged from 34 cases, Table S1)

- Forelimbs: the model rotates forward its forelimbs and spreads them ($S_{pitch}$ = 80°, $E_{pitch}$ = 136°, $W_{pitch}$ = 178°). In mockingbirds, the wings are slightly rotated forward and raised to a degree less than 90°. Observing this, we set the S value in the dinosaur model to 80°. Since it appears that birds often spread their wings to the maximum extent possible, we similarly allow the dinosaur model to spread its forelimbs maximally.

The third pause lasts for 0.15 sec (averaged from 34 cases, Table S1).

Phase 4. Folding forelimbs to return to the resting posture for 0.208 sec (based on 34 cases, Table S1)

- Forelimbs: the resting posture speculated by senter & robins[2] ($S_{pitch}$ = 33°, $S_{yaw}$ = 0°, $S_{roll}$ = 0°, $E_{pitch}$ = 106°, $E_{yaw}$ = 0°, $E_{roll}$ = 0°, $W_{pitch}$ = 131°, $W_{yaw}$ = 0°, $W_{roll}$ = 0°).

#### 1G) Animations in Experiment 4: forelimb forward-move preceded by walking

*Experiment 4. Imitating the display and walking of the greater roadrunner* (Video S6)
We aimed to assess the effect of walking in a dinosaur animation, imitating the display of the greater roadrunner. We selected the roadrunner's display for its simplicity compared to the two other birds considered here. The dinosaur model imitated the stride parameters of the greater roadrunner with a stride length of 0.25 m (averaged from five cases: 0.08, 0.21, 0.25, 0.26, 0.28, and 0.40 m) and a stride speed of 0.97 m/s (based on from five cases: 0.42, 0.57, 0.76, 0.85, 1.51, and 1.72 m/s). Stride length is defined as the distance covered during the completion of two steps, one with each foot. The number of steps between flush displays was set to four (based on five cases: 2, 2, 4, 7, and 7 steps), resulting in the dinosaur walking for a total distance of 1 m (0.25 m per step multiplied by four steps). This walking phase was integrated into the front part of the animations, imitating the display of the greater roadrunner in Exp. 1.

Observing variations in head bobbing during the greater roadrunner's walking (https://macaulaylibrary.org/asset/296951221, © Mark A. Brogie), we encountered challenges in replicating this behavior due to the lack of a clear association between head bobbing and leg movements[23]. Consequently, we chose to maintain a consistent head position in the dinosaur model during walking. To simplify the animation, we kept the tail stationary while walking, as tail movements were not prominent in the roadrunner and were not our primary focus. This walking phase was integrated into the front part of the animations, imitating the display of the greater roadrunner.

#### 1H) Experimental set-up

The animations were presented upside down since the locusts were positioned with their ventral side up during the experiments. To synchronize the timing between the neural recording and the animation, we incorporated a full black screen lasting one frame (0.0083 sec), occurring 0.5 sec after the animation onset and 1 sec before the dinosaur initiates the flush display. The dinosaur model was rendered black (R – 000, G – 000, B – 000), contrasting with a white background (R – 255, G – 255, B – 255). For animations, we set the frame rate to 120 frames per second, exceeding the flicker fusion frequency of the locust eye (40–90 Hz[24]).

#### 1I) MATLAB codes for extracting variables from animations

##### *Counting black pixels in each animation frame*

```
% Define the path to the directory containing PNG files
directoryPath = 'C:\Users\location';

% Get a list of PNG files in the directory
pngFiles = dir(fullfile(directoryPath, '*.png'));

% Initialize array to store information
blackPixelCounts = zeros(1, length(pngFiles));

% Iterate through each PNG file
for i = 1:length(pngFiles)
    % Read the image
    imagePath = fullfile(directoryPath, pngFiles(i).name);
    img = imread(imagePath);

    % Count black pixels
    blackPixelCounts(i) = sum(img(:) == 1);
end

% Create a table with the results
resultsTable = table((1:length(blackPixelCounts))', blackPixelCounts', 'VariableNames',
{'FileIndex', 'BlackPixelCount'});

% Save the table to an Excel file
writetable(resultsTable, 'black_pixel_counts.xlsx');
```

##### *Counting pixels undergoing a color transition in each animation frame*

```
% Define the path to the directory containing PNG files
directoryPath = 'C:\Users\location';

% Get a list of PNG files in the directory
pngFiles = dir(fullfile(directoryPath, '*.png'));

% Initialize arrays to store information
blackToWhiteChanges = zeros(1, length(pngFiles) - 1);
whiteToBlackChanges = zeros(1, length(pngFiles) - 1);

% Iterate through each pair of consecutive PNG files
for i = 1:length(pngFiles)-1
    % Read the current and next images
```

```
    currentImagePath = fullfile(directoryPath, pngFiles(i).name);
    nextImagePath = fullfile(directoryPath, pngFiles(i+1).name);

    currentImg = imread(currentImagePath);
    nextImg = imread(nextImagePath);

    % Count changes from black to white and vice versa
    blackToWhiteChanges(i) = sum(currentImg(:) == 1 & nextImg(:) == 255);
    whiteToBlackChanges(i) = sum(currentImg(:) == 255 & nextImg(:) == 1);
end

% Create a table with the results
resultsTable = table((1:length(blackToWhiteChanges))', blackToWhiteChanges',
whiteToBlackChanges', 'VariableNames', {'Event', 'BlackToWhiteChanges', 'WhiteToBlackChanges'});

% Save the table to an Excel file
writetable(resultsTable, 'results.xlsx');
```

#### 1J) Extracellular recordings

We fixed the locust on a wooden holder ventral side up. We carefully tilted the head backward using a pin to expose the neck. Beeswax was applied to both sides of the neck as a barrier to retain the saline solution. Right eye was covered with beeswax to block visual input, while left eye faced the monitor screen. Under stereoscope, we dissected the ventral part of the neck to uncover the ventral nerve cords. Saline solution (NaCl 210 mM, KCl 7.1 mM, CaCl2 9.0 mM, Tris buffered to pH 6.8) was carefully dropped onto the dissected area. An extracellular silver wire hook electrode (127-µm bare diameter, AM systems) in an electrode holder (H-13, Narishige) attached to a micromanipulator (MM-3, Narishige) was connected to the contralateral nerve cord. A pin secured in the abdomen was connected to the ground line. The electrode was connected to the Neuron SpikerBox Pro (Backyard Brains, USA), which was connected to a laptop. The neural activity in response to animations was recorded at a sampling rate of 10 kHz. Animations were presented on a flat-screen monitor (TFG32Q14P IPS QHD 144, Hansung computer; monitor screen size: 32 inch) with a display brightness of 400 cd/m$^2$ and a refresh rate of 120 Hz. The distance between the monitor and the locust was set at 35 cm. To minimize recording noise, cables were used to ground the equipment during the experiments.

We used Spike2 software (version 5, Cambridge Electronic Design, Cambridge) to analyze the neural spike data. The software identified DCMD spikes based on the spike amplitude and general shape. This method aligns with numerous previous studies[16–20]. Subsequently, the spiking rate of the DCMD neuron was calculated using a bin size of 20 ms.

### Part 2: Results of Pilot Experiments

#### 2A) Text

We confirmed that 3 out of 5 tested locusts responded by jumping (Table S2) to the expanding simple circular looming stimulus on the computer screen (an approaching circle; $l/|v|$ = 5 ms, where $l$ represents the radius of 3 cm and $v$ is the constant approaching speed of 6 m/s). Furthermore, in two individuals that jumped during the behavioral tests, we confirmed that the recorded spiking rate, measured a day after, increased as the circle expanded (Fig. S2; Table S3). The spike shapes recorded in these tests (Fig. S2A, B) were similar to those observed in the subsequent experiments (see insets with examples of spike shapes in Extended Data Figs. S2–S4; Fig. S12).

Locusts jumped more frequently (Extended Data Fig. 1a; Table S4), and the peak spiking rate (Extended Data Fig. 1b; Table S4) as well as the frequency of reaching the hypothetical jump preparation threshold (5 spikes/20 ms[16]; Extended Data Fig. 1c; Table S4) were higher for the close animation compared to the distant one (Video S2), confirming that the differences in the DCMD response between close and distant animations were consistent with the behavioral differences in jump responses. Most jumps were initiated when the animated dinosaur folded its forelimbs (yellow shading in Extended Data Fig. 1d).

#### 2B) Figure

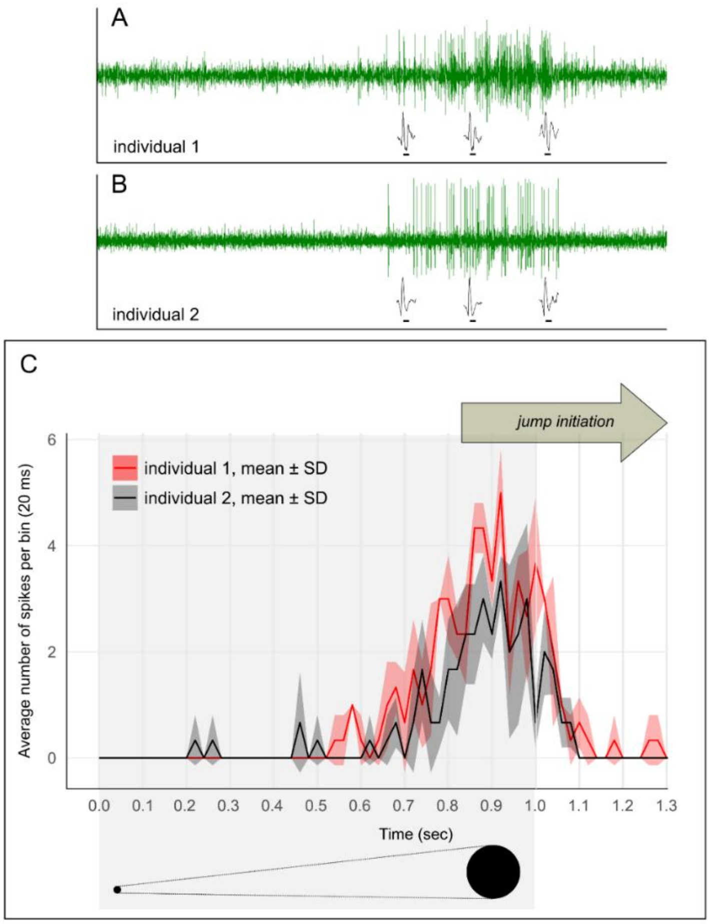


**Figure S2. Locust's LGMD/DCMD response to a looming circle animation.** (A, B) Examples of neural recordings from each individual; zoomed-in spikes are shown to visualize their shapes. (C) Spiking rate of the LGMD/DCMD escape pathway (mean ± SD, bin size = 30 ms) in response to a looming circle animation. The two lines depict profiles of average spiking rate of individual 1 (red; n = 3 recordings) and individual 2 (black; n = 3 recordings) during the animation of a looming circle approach at the locust. In the behavioral experiments, using the same animals and animation, jump initiation was observed in all 3 trials for individual 1 and in 2 trials for individual 2. The timing of jump initiation ranged from 0.83 to 1.98 sec, which would place it after the peak spiking rate.

#### 2C) Tables

**Table S2. Behavioral responses to the simple looming stimulus in five individuals tested in behavioral tests.** Bold text indicates the individuals (1 and 4) from which neural recordings were collected.

| Individual | Trial | Take-off initiation timing (unit: sec) |
|---|---|---|
| **1** | 1 | 0.4 sec after the end of the animation |
| | 2 | 0.73 sec after the end of the animation |
| | 3 | 0.17 sec before the end of the animation |
| 2 | 1 | No jump |
| | 2 | 0.27 sec after the end of the animation |
| | 3 | 0.1 sec after the end of the animation |
| | 4 | No jump |
| 3 | 1 | No jump |
| | 2 | No jump |
| | 3 | No jump |
| | 4 | No jump |
| **4** | 1 | 0.63 sec after the end of the animation |
| | 2 | 0.98 sec after the end of the animation |
| 5 | 1 | No jump |
| | 2 | No jump |
| | 3 | No jump |
| | 4 | No jump |
| | 5 | No jump |

**Table S3. Response of the LGMD/DCMD pathway to an animation of a simple looming stimulus.** The number of recorded DCMD spikes in response to the “looming circle” animation for two individual locusts. The spike numbers are summed up in every bin (20 ms). We recorded the neural response three times for each individual.

| Bin order | Individual 1 | | | Individual 2 | | |
|---|---|---|---|---|---|---|
| (size: 20 ms) | Record1 | Record2 | Record3 | Record1 | Record2 | Record3 |
| 1 | 0 | 0 | 0 | 0 | 0 | 0 |
| 2 | 0 | 0 | 0 | 0 | 0 | 0 |
| 3 | 0 | 0 | 0 | 0 | 0 | 0 |
| 4 | 0 | 0 | 0 | 0 | 0 | 0 |
| 5 | 0 | 0 | 0 | 0 | 1 | 0 |
| 6 | 0 | 0 | 0 | 0 | 0 | 0 |
| 7 | 0 | 0 | 0 | 0 | 1 | 0 |
| 8 | 0 | 0 | 0 | 0 | 0 | 0 |
| 9 | 0 | 0 | 0 | 0 | 0 | 0 |
| 10 | 0 | 0 | 0 | 0 | 0 | 0 |
| 11 | 0 | 0 | 0 | 0 | 0 | 0 |
| 12 | 0 | 0 | 0 | 0 | 0 | 0 |
| 13 | 0 | 0 | 0 | 0 | 1 | 0 |
| 14 | 0 | 0 | 0 | 0 | 1 | 0 |
| 15 | 0 | 0 | 1 | 0 | 1 | 0 |
| 16 | 0 | 0 | 1 | 0 | 0 | 0 |
| 17 | 0 | 0 | 1 | 0 | 0 | 0 |
| 18 | 0 | 2 | 0 | 0 | 0 | 1 |
| 19 | 1 | 0 | 1 | 0 | 0 | 0 |
| 20 | 0 | 1 | 2 | 0 | 1 | 1 |
| 21 | 1 | 2 | 4 | 1 | 0 | 0 |
| 22 | 2 | 0 | 1 | 0 | 0 | 1 |
| 23 | 1 | 3 | 5 | 3 | 1 | 0 |
| 24 | 1 | 3 | 4 | 4 | 0 | 2 |
| 25 | 2 | 5 | 3 | 2 | 1 | 3 |
| 26 | 6 | 3 | 7 | 3 | 3 | 3 |
| 27 | 4 | 5 | 7 | 5 | 3 | 4 |
| 28 | 5 | 6 | 3 | 4 | 6 | 5 |
| 29 | 5 | 6 | 5 | 3 | 4 | 3 |
| 30 | 8 | 3 | 5 | 2 | 3 | 4 |
| 31 | 5 | 6 | 3 | 5 | 3 | 1 |
| 32 | 4 | 5 | 3 | 3 | 2 | 4 |
| 33 | 6 | 3 | 2 | 2 | 2 | 2 |

**Table S4. Generalized mixed-effects model for analyzing jump occurrence, peak size, and threshold variations to prototype dinosaur animations imitating relatively close and distant animations in pilot experiment.** For the binary variables of jump occurrence (jumped = 1, non-jumped = 0) and threshold (reached = 1, unreached = 0), we used the Binomial distribution with a logit link function. For the under-dispersed count of peak size (no. spikes/20 ms), we utilized the Conway-Maxwell poisson family with a log link function. The fixed effects include animation distance (close or distance) and sex (male or female). Individual ID with 6 levels was used as a random effect. The table values correspond to the effect estimate, standard error, Z value, and P-value for each variable. P-values below 0.05 are denoted in bold, except for the intercept. This table concerns Extended Data Fig. 1.

| Response variable | Fixed effect | Estimate | Standard Error | Z value | P-value |
|---|---|---|---|---|---|
| Jump occurrence | Intercept | -1.039 | 0.575 | -1.806 | 0.071 |
| | **Animation distance** *'close'* | 2.077 | 0.678 | 3.066 | **0.002** |
| | Sex *'male'* | -0.655 | 0.671 | -0.977 | 0.329 |
| Peak size | Intercept | 1.322 | 0.094 | 14.138 | < 0.001 |
| | **Animation distance** *'close'* | 0.212 | 0.060 | 3.601 | **< 0.001** |
| | Sex *'male'* | -0.163 | 0.125 | -1.304 | 0.192 |
| Threshold | Intercept | -1.807 | 1.451 | -1.245 | 0.213 |
| | **Animation distance** *'close'* | 2.177 | 1.039 | 2.096 | **0.036** |
| | Sex *'male'* | -1.763 | 1.860 | -0.948 | 0.343 |

### Part 3: Results of the three main experiments

#### 3A) Profiles of average spiking rates for each individual

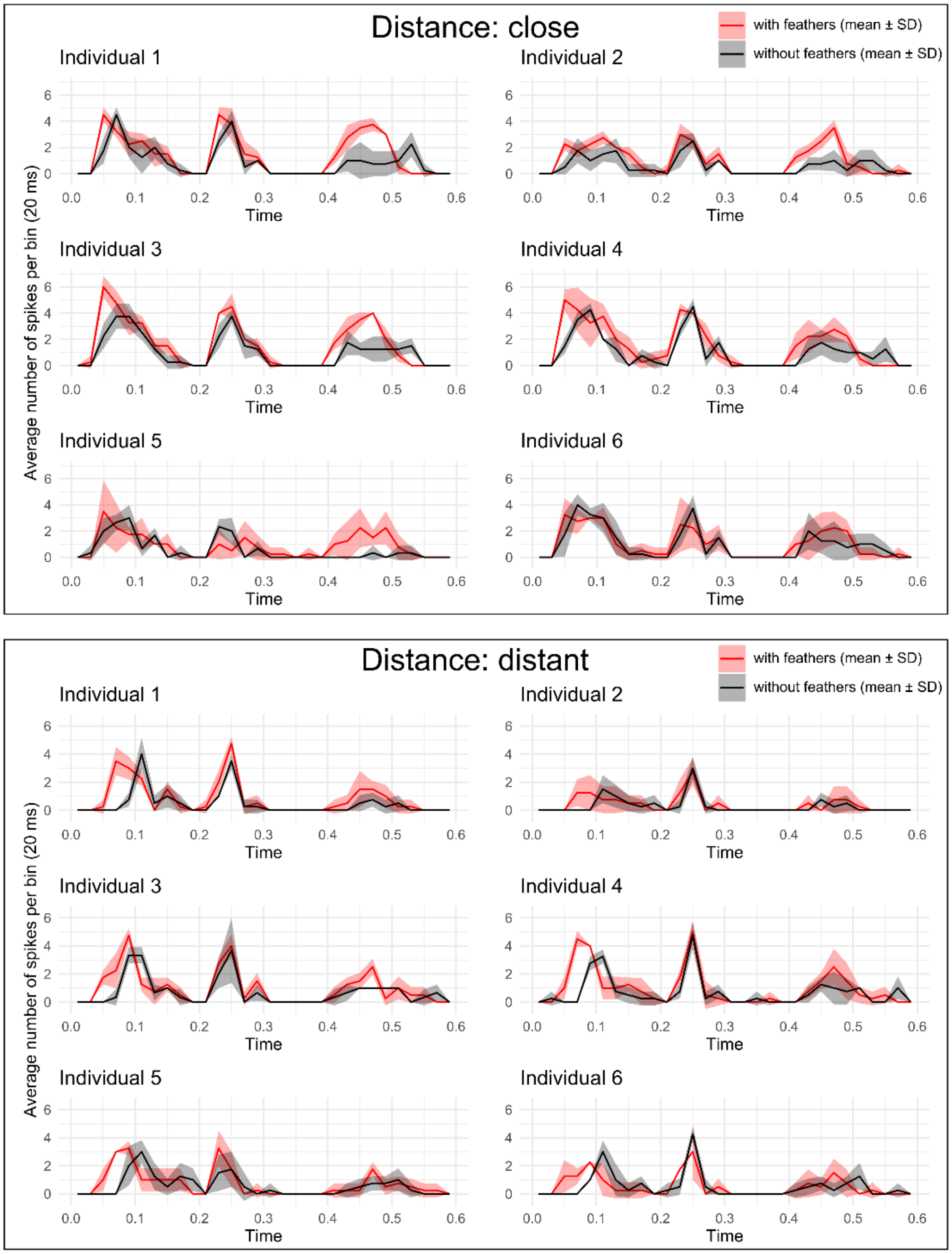


**Figure S3. Average spiking rates per bin (no. spikes/20 ms; mean ± SD) of the locust's LGMD/DCMD escape pathway for each individual in response to animations imitating the display of *Geococcyx californianus* in Experiment 1.** Each small panel represents a profile for the mean spiking rate per bin from n = 4 recordings in response to animations "with" (red line) or "without" (black line) feathers. Shading represents the standard deviation. The X-axis represents time. The upper box represents the spiking rate for animations imitating a close situation (60 cm to prey), while the lower box represents the spiking rate for animations imitating a distant case (120 cm to prey).

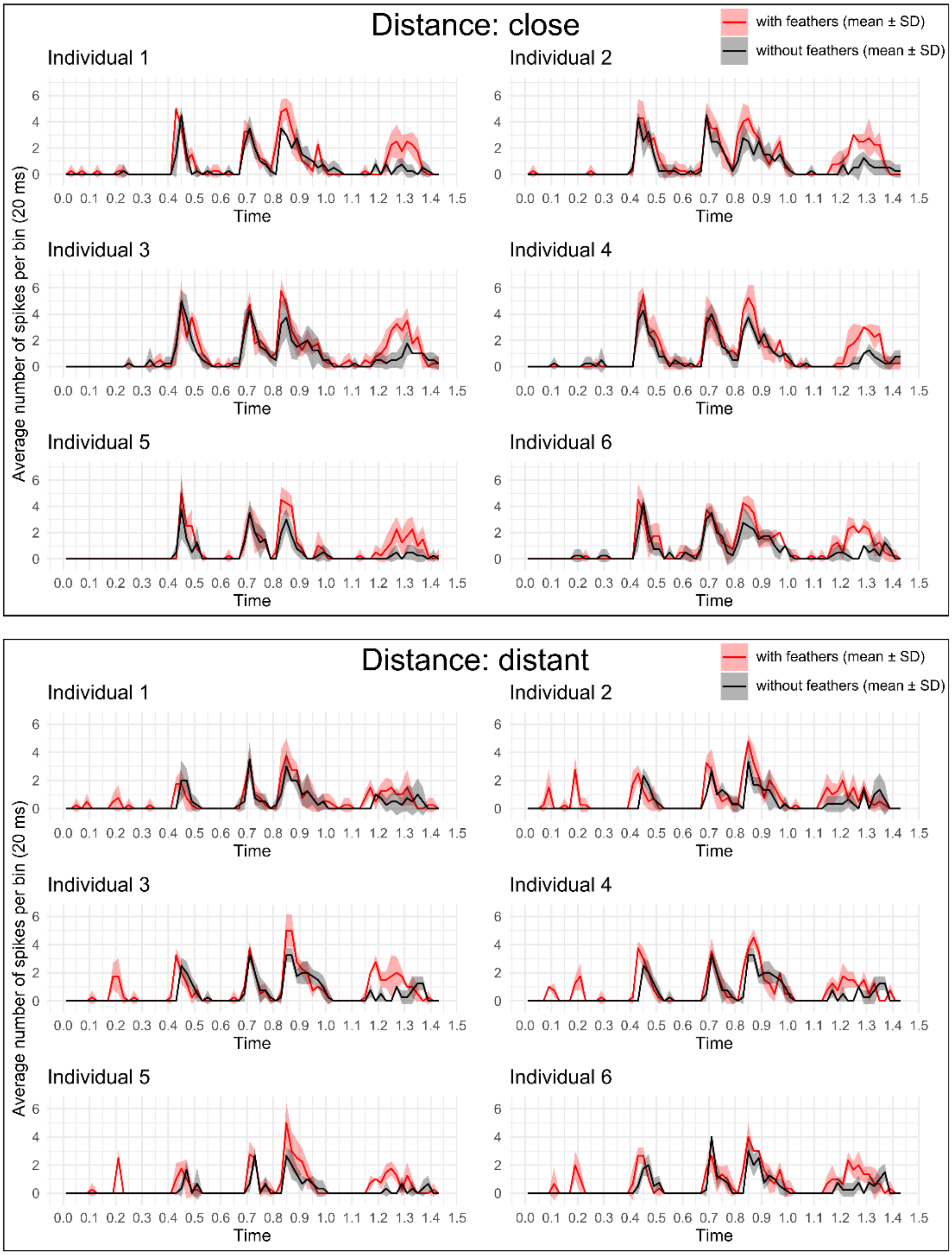


**Figure S4. Average spiking rate per bin (no. spikes/20 ms; mean ± SD) of the locust's LGMD/DCMD escape pathway for each individual in response to animations imitating the display of *Cercotrichas galactotes* in Experiment 2.** Each small panel represents the spiking rate per bin from n = 4 recordings in response to animations "with" (red line) or "without" (black line) feathers. Shading represents the standard deviation. The X-axis represents time. The upper box represents the spiking rate for animations imitating a close situation (60 cm to prey), while the lower box represents the spiking rate for animations imitating a distant case (120 cm to prey).

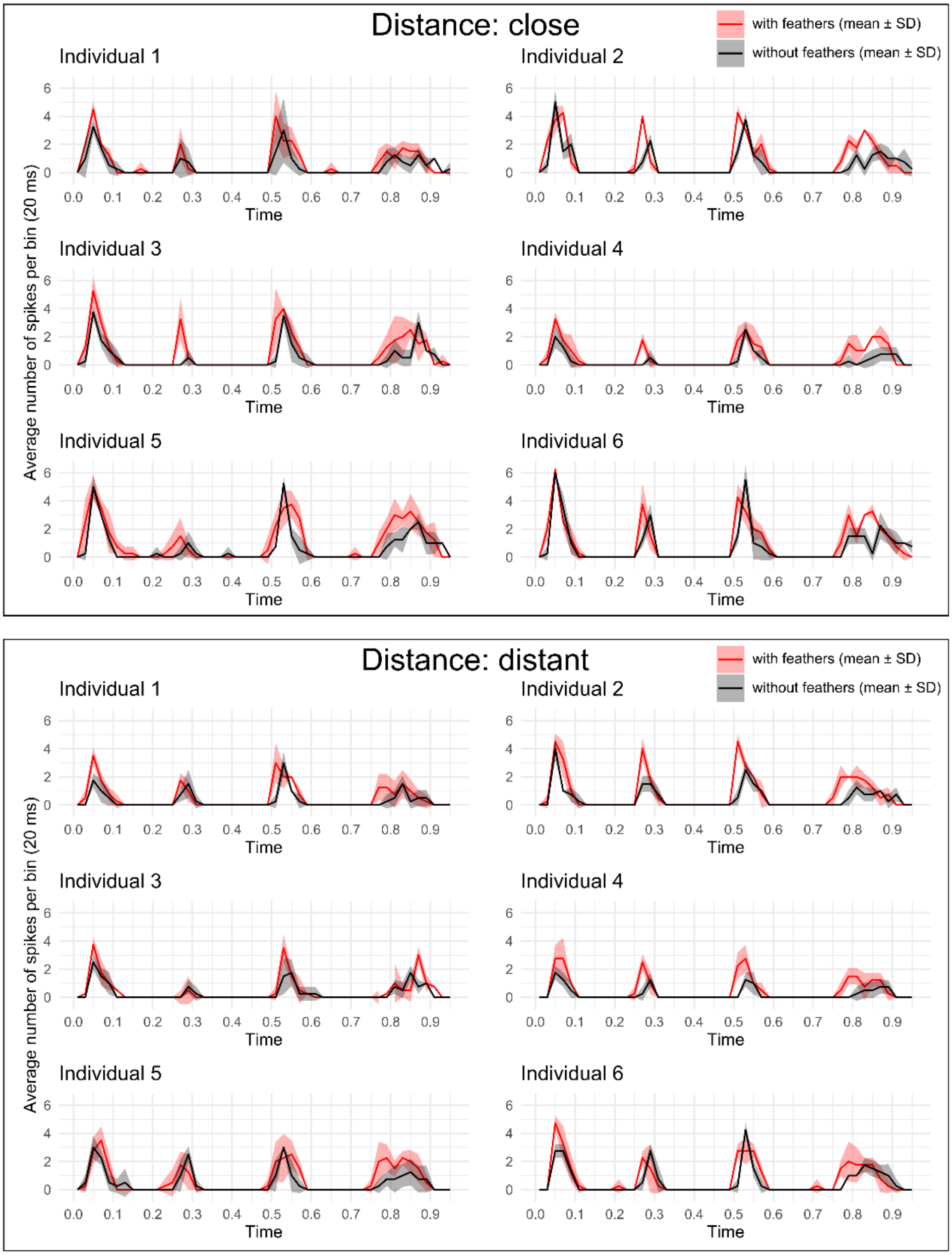


**Figure S5. Average spiking rate per bin (no. spikes/20 ms; mean ± SD) of the locust's LGMD/DCMD escape pathway for each individual in response to animations imitating the display of *Mimus polyglottos* in Experiment 3.** Each small panel represents the spiking rate per bin based on n = 4 recordings in response to animations "with" (red line) or "without" (black line) feathers. Shading represents the standard deviation. The X-axis represents time. The upper box represents the spiking rate for animations imitating a close situation (60 cm to prey), while the lower box represents the spiking rate for animations imitating a distant case (120 cm to prey).

3B) DCMD responses and profiles of angular size and angular velocity based on forelimb/tails tips in animations

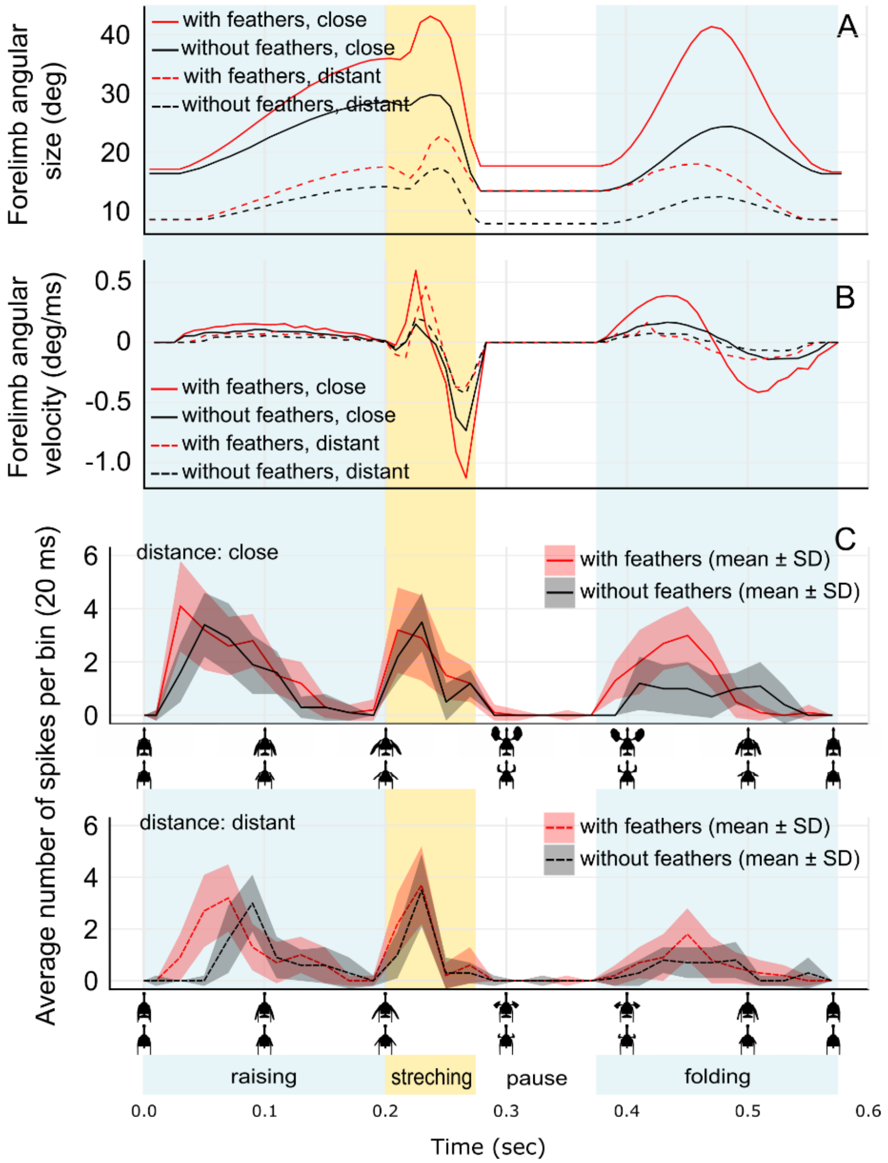


**Figure S6. Profiles of angular size and angular velocity based on forelimb tips in animations, and DCMD responses of locusts to those animations imitating the display of *Geococcyx californianus* in Experiment 1 by dinosaurs "with" (red line) or "without" (black line) feathers at "close" (solid line) or "distant" (dashed line) locations.** (A) Angular size (deg) of the distance between the forelimb tips. (B) Angular velocity (deg/ms) based on (A). (C) Average spiking rate per bin (no. spikes/20 ms; mean ± SD; n = 24, i.e., four recordings from each of six individuals) of the locust's LGMD/DCMD escape pathway in response to animations "with" (red) and "without" feathers (black) at "close" (60cm; upper panel) and "distant" (120cm; lower panel) locations. Shading represents the standard deviation. Time (sec) is plotted on the X-axis, with corresponding screenshots below. Display phases are indicated by color-shaded vertical boxes.

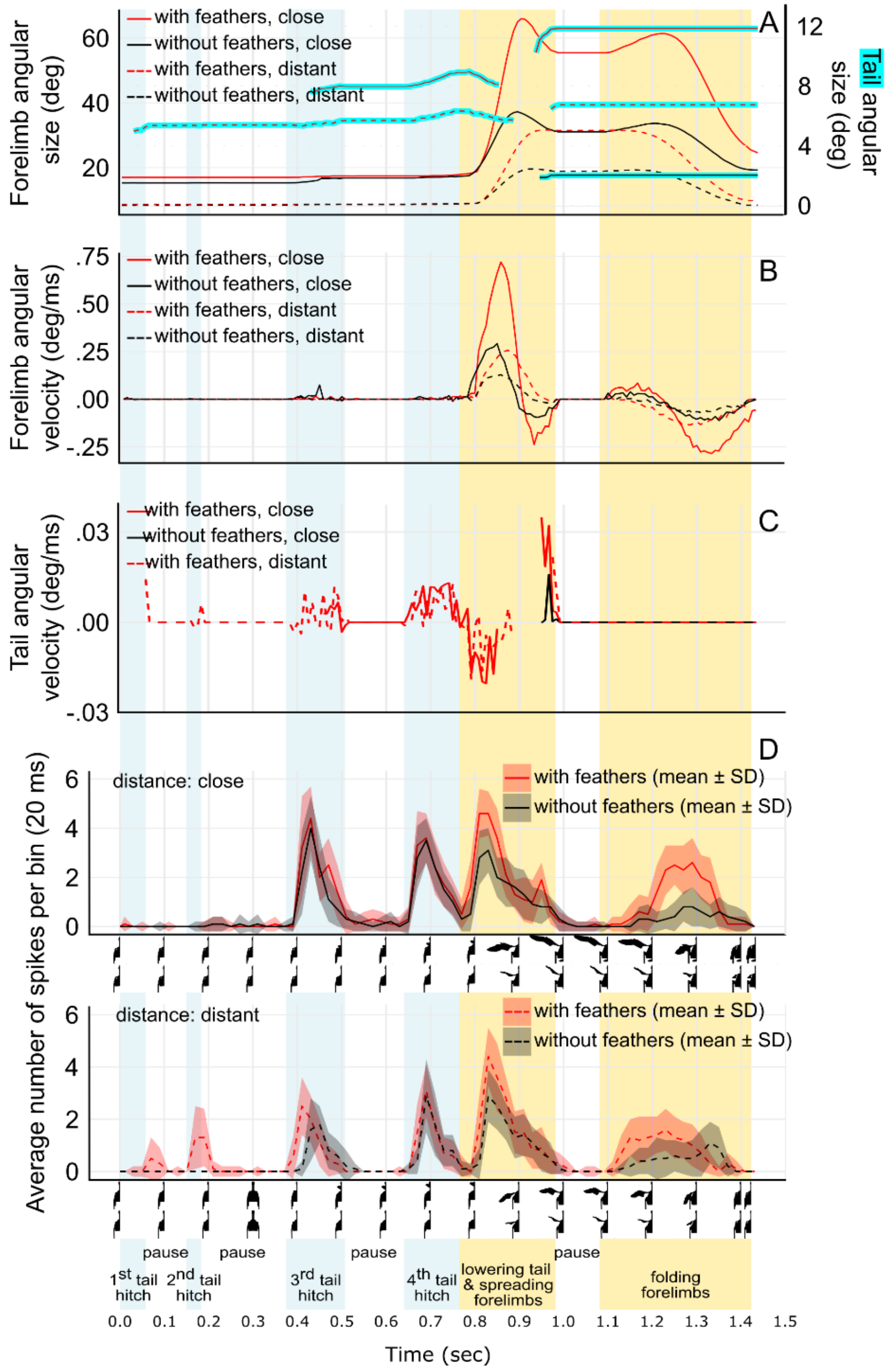


**Figure S7. Profiles of angular size and angular velocity based on forelimb tips and tail corners in animations (A-C), and DCMD responses (D) of locusts to those animations imitating the display of *Cercotrichas galactotes* in Experiment 2 by dinosaurs "with" (red line) or "without" (black line) feathers at "close" (solid line) or "distant" (dashed line) locations.** (A) Angular size (deg) for the distance between the left and right forelimb tips and for the distance between left and right tail corners. (B) Angular velocity (deg/ms) based on angular distance between forelimb tips in (A). (C) Angular velocity (deg/ms) based on angular distance between left and right tail tip in (A). (D) Average spiking rate per bin (no. spikes/20 ms; mean ± SD; n = 24, i.e., four recordings from each of six individuals) of the locust's LGMD/DCMD escape pathway in response to animations "with" (red) and "without" feathers (black) at "close" (60cm; upper panel) and "distant" (120cm; lower panel) locations. Shading represents the standard deviation. Time (sec) is plotted on the X-axis, with corresponding half-screenshots below. Display phases are indicated by color-shaded vertical boxes.

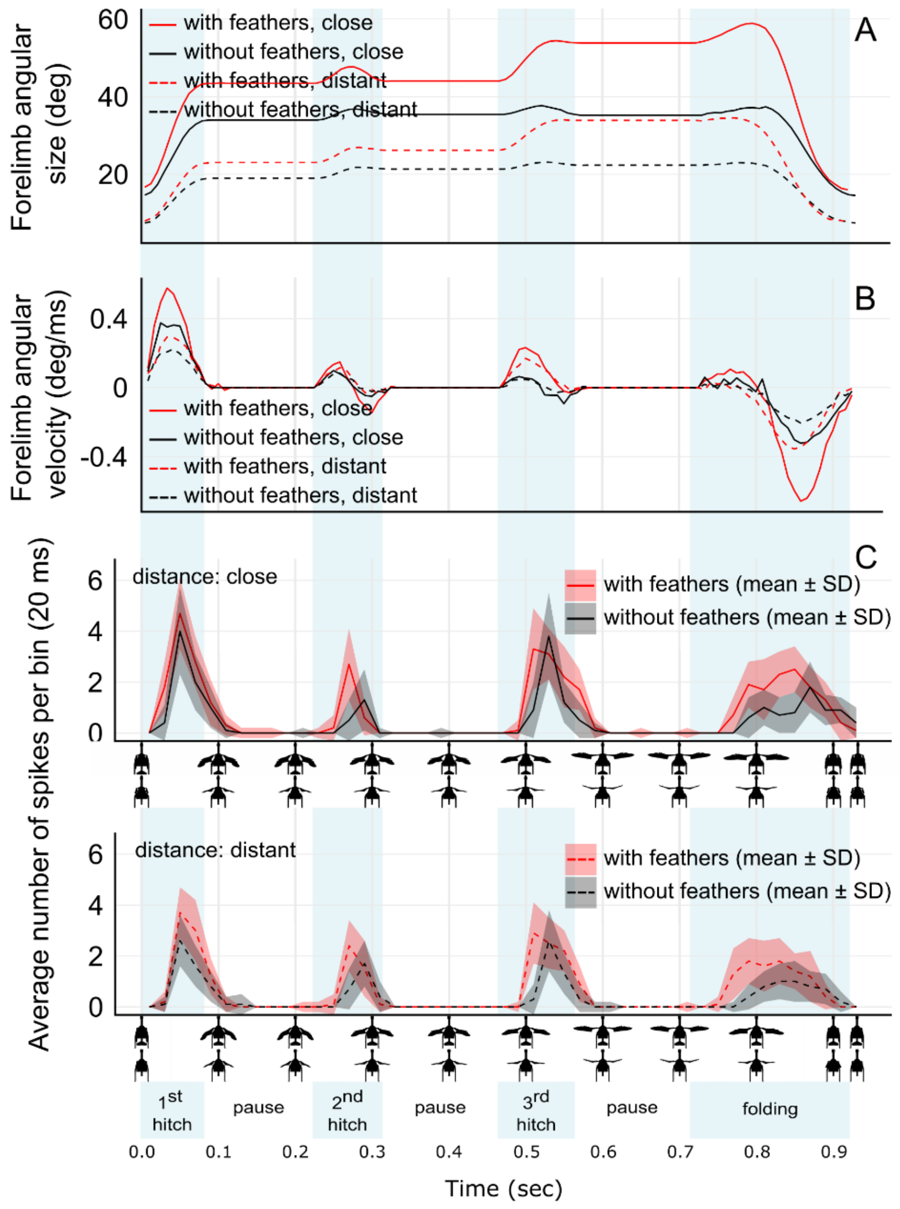


**Figure S8. Profiles of angular size and angular velocity based on forelimb tips in animations (A, B), and DCMD responses (C) of locusts to those animations imitating the display of *Mimus polyglottos* in Experiment 3 by dinosaurs "with" (red line) or "without" (black line) feathers at "close" (solid line) or "distant" (dashed line) locations.** (A) Angular size (deg) of the distance between the forelimb tips. (B) Angular velocity (deg/ms) based on (A). (C) Average spiking rate per bin (no. spikes/20 ms; mean ± SD; n = 24, i.e., four recordings from each of six individuals) of the locust's LGMD/DCMD escape pathway in response to animations "with" (red) and "without" feathers (black) at "close" (60cm; upper panel) and "distant" (120cm; lower panel) locations. Shading represents the standard deviation. Time (sec) is plotted on the X-axis, with corresponding screenshots below. Display phases are indicated by color-shaded vertical boxes.

#### 3C) DCMD response and profiles of visual expansion of dinosaur silhouette and edge movements

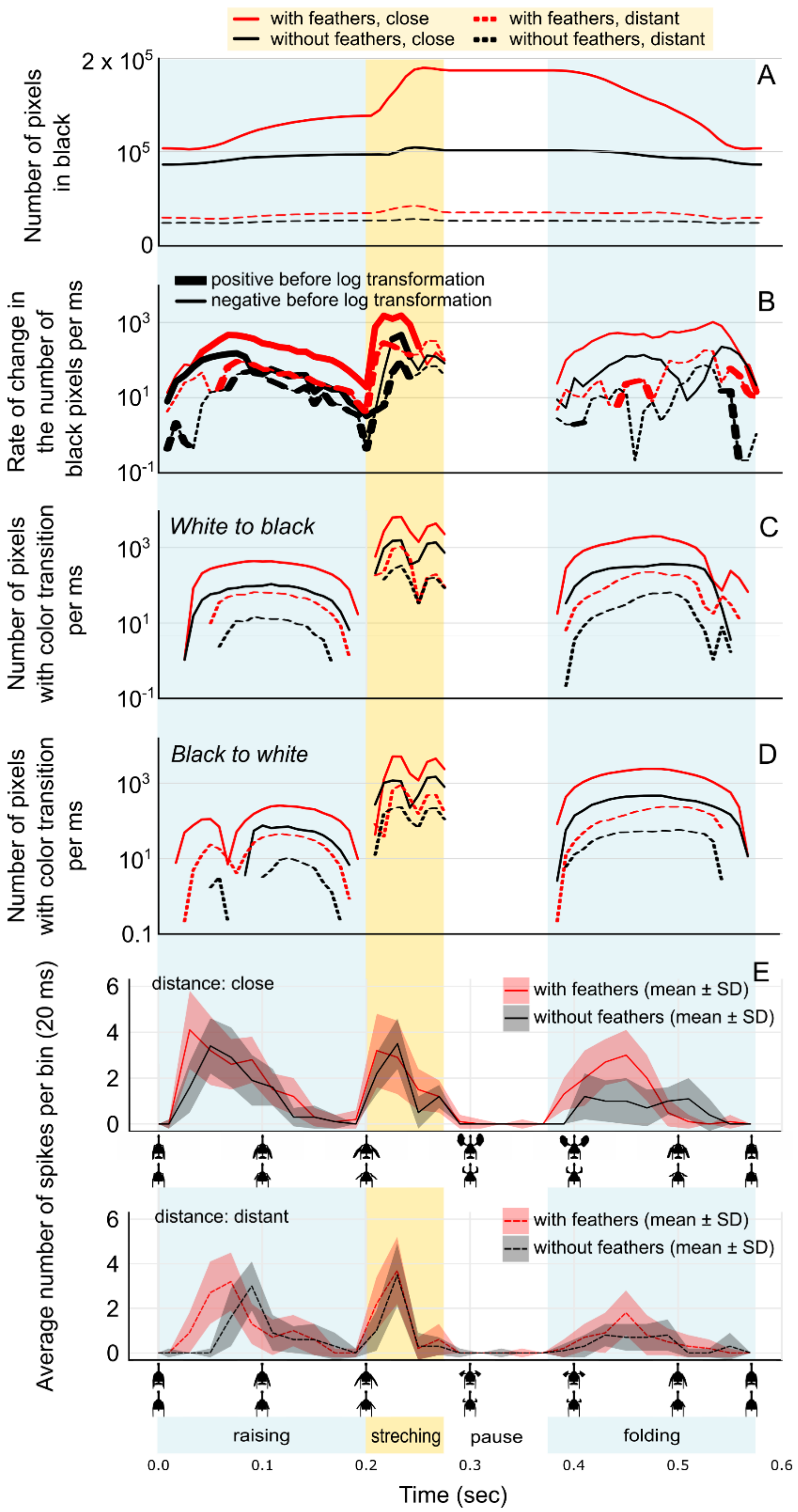


**Figure S9. Profiles of four variables concerning the visual expansion of the dinosaur silhouette and edge movements extracted from the animations (A–D), and DCMD responses (E, F) to animations imitating the display of *Geococcyx californianus* in Experiment 1 by dinosaurs "with" (red line) or "without" (black line) feathers at "close" (solid line) or "distant" (dashed line) locations.** (A) Number of pixels in black per frame. (B) Rate of change in the number of black pixels per ms based on change observed between consecutive frames. Positive values before applying the log scale to the Y-axis are shown as thick lines, while negative values are represented as thin lines. (C) Rate of white-to-black transition in the stimulus represented by the number of pixels with white-to-black color transition per ms based on the number of pixels with color transition observed between two consecutive frames during animation. (D) Rate of black-to-white transition in the stimulus represented by the number of pixels with black-to-white color transition per ms based on the number of pixels with color transition observed between two consecutive frames during animation. (E) Average spiking rate per bin (no. spikes/20 ms; mean ± SD; n = 24, i.e., four recordings from each of six individuals) of the locust's LGMD/DCMD escape pathway in response to animations "with" (red) and "without" feathers (black). Upper panel in E represents the imitation of a close situation (60 cm to prey), while lower panels represent a distant case (120 cm to prey). Time (sec) is plotted on the X-axis, with corresponding screenshots below. Display phases are indicated by color-shaded vertical boxes. For panels B, C, and D, a logarithmic scale was applied to the Y-axis for better visualization.

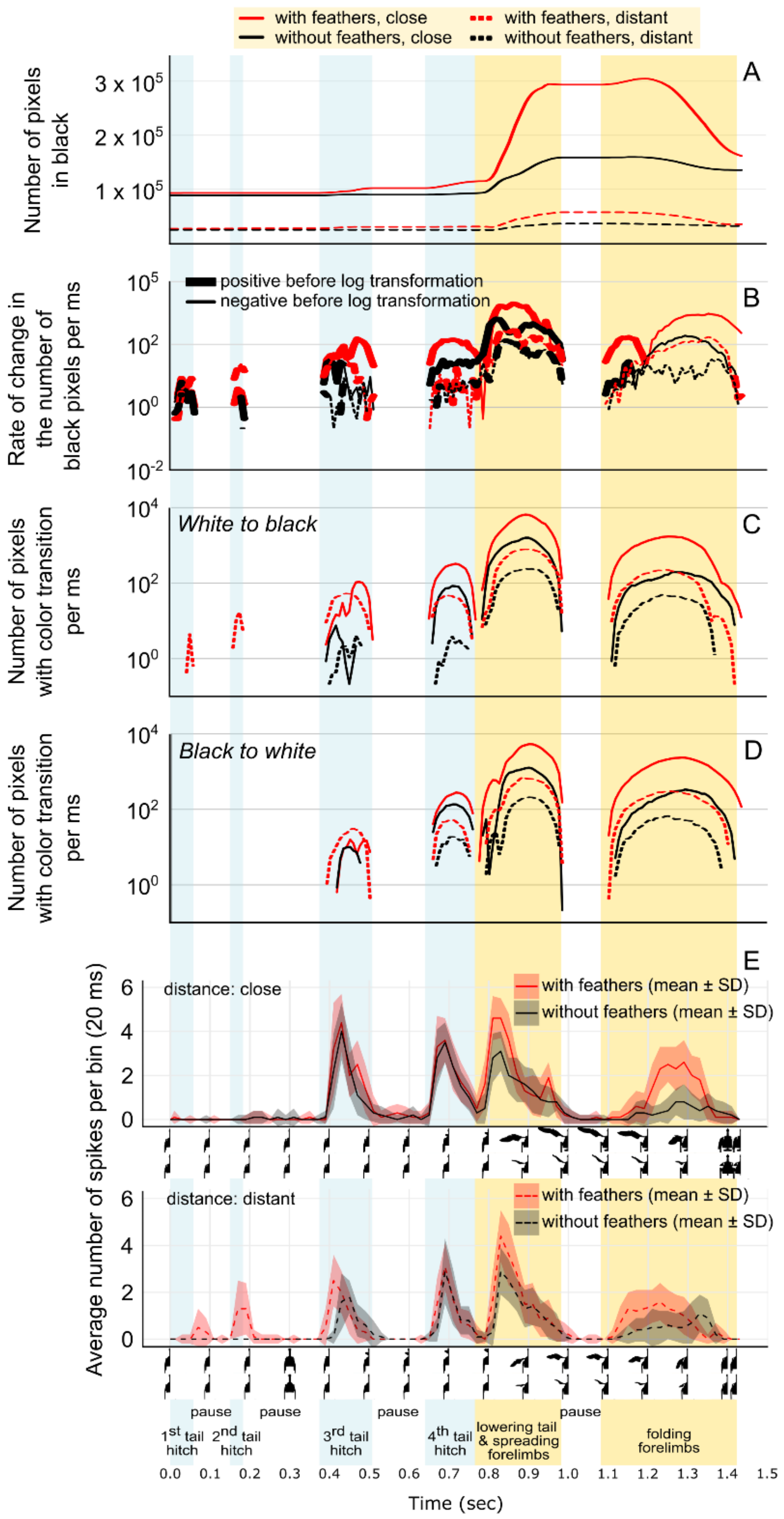


**Figure S10. Profiles of four variables concerning the visual expansion of the dinosaur silhouette and edge movements extracted from the animations (A–D), and DCMD responses (E, F) to animations imitating the display of *Cercotrichas galactotes* in Experiment 2 by dinosaurs "with" (red line) or "without" (black line) feathers at "close" (solid line) or "distant" (dashed line) locations.** (A) Number of pixels in black per frame. (B) Rate of change in the number of black pixels per ms based on change observed between two consecutive frames. Positive values before applying the log scale to the Y-axis are shown as thick lines, while negative values are represented as thin lines. (C) Number of pixels with white-to-black color transition per ms based on the number of pixels with color transition observed between two consecutive frames during animation. (D) Number of pixels with black-to-white color transition per ms based on the number of pixels with color transition observed between two consecutive frames during animation. (E) Average spiking rate per bin (no. spikes/20 ms; mean ± SD; n = 24, i.e., four recordings from each of six individuals) of the locust's LGMD/DCMD escape pathway in response to animations "with" (red) and "without" feathers (black). Upper panel in E represents the imitation of a close situation (60 cm to prey), while lower panels represent a distant case (120 cm to prey). Time (sec) is plotted on the X-axis, with corresponding screenshots below. Display phases are indicated by color-shaded vertical boxes. For panels B, C, and D, a logarithmic scale was applied to the Y-axis for better visualization.

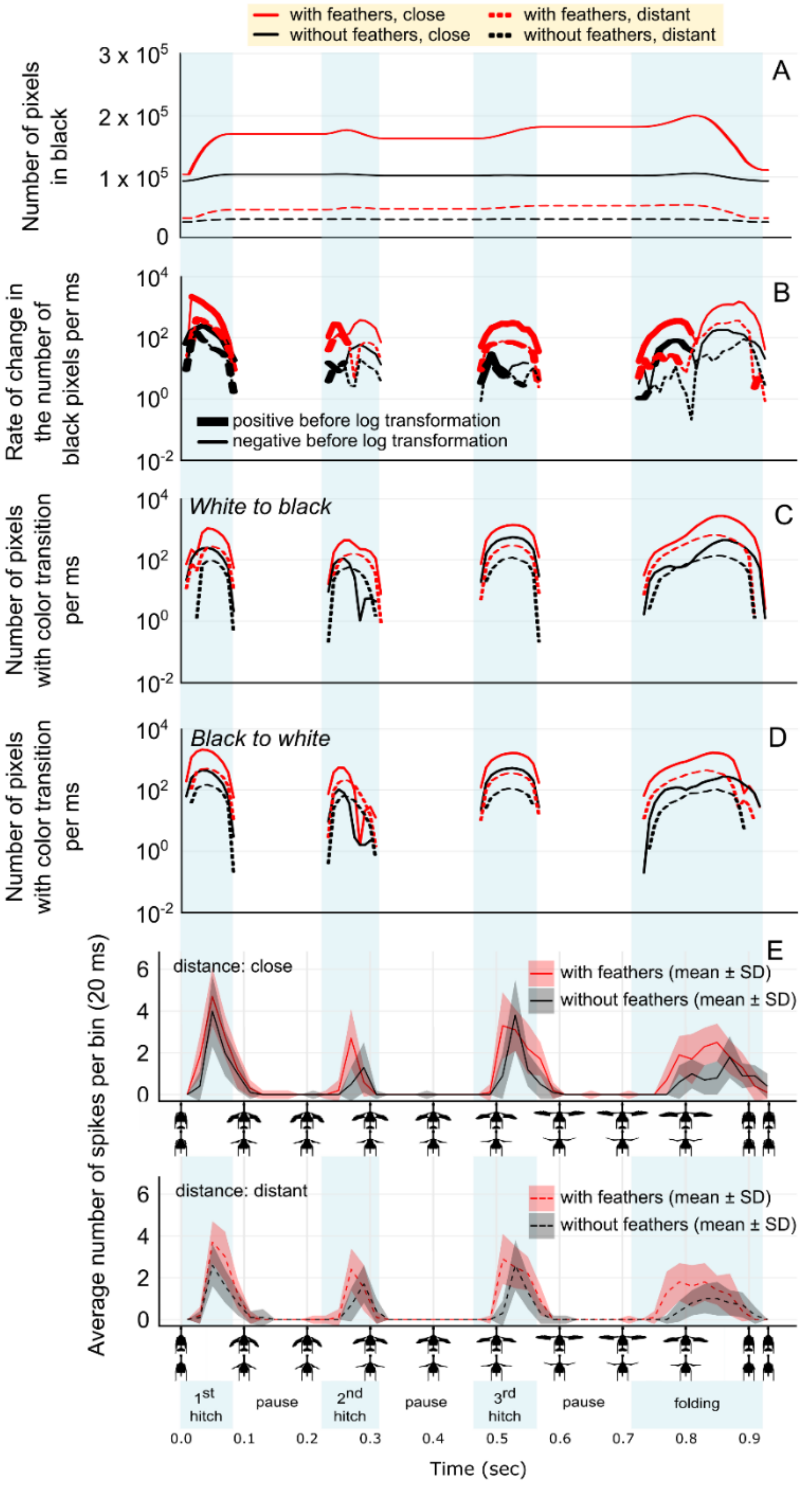


**Figure S11. Profiles of four variables concerning the visual expansion of the dinosaur silhouette and edge movements extracted from the animations (A–D), and DCMD responses (E, F) to animations imitating the display of *Mimus polyglottos* in Experiment 3 by dinosaurs "with" (red line) or "without" (black line) feathers at "close" (solid line) or "distant" (dashed line) locations.** (A) Number of pixels in black per frame. (B) Rate of change in the number of black pixels per ms based on change observed between consecutive frames. Positive values before applying the log scale to the Y-axis are shown as thick lines, while negative values are represented as thin lines. (C) Number of pixels with white-to-black color transition per ms based on the number of pixels with color transition observed between two consecutive frames during animation. (D) Number of pixels with black-to-white color transition per ms based on the number of pixels with color transition observed between two consecutive frames during animation. (E) Average spiking rate per bin (no. spikes/20 ms; mean ± SD; n = 24, i.e., four recordings from each of six individuals) of the locust's LGMD/DCMD escape pathway in response to animations "with" (red) and "without" feathers (black). Upper panel in E represents the imitation of a close situation (60 cm to prey), while lower panels represent a distant case (120 cm to prey). Time (sec) is plotted on the X-axis, with corresponding screenshots below. Display phases are indicated by color-shaded vertical boxes. For panels B, C, and D, a logarithmic scale was applied to the Y-axis for better visualization.

#### 3D) Text – commentary to Extended Data Figs. 6–9 and Figs S6–S11 and S14–S15

In all experiments, the timing of neural response peaks roughly coincided with increases in the angular velocity of expansion or contraction of the stimulus measured between the forelimb tips, and in experiment 2, also the tail tip corners (panel B in Figs. S6–S8, S14). Increases in spiking rate leading to these peaks approximately coincided with increases in the rate of change in the number of black pixels (panel B in Figs. S9–S11, S15) and the rate of black-to-white pixel transitions (panels C and D in Figs. S9–S11, S15). These variables represent changes in silhouette area and edge movement speed in the animations. The strongest alignments in both timing and magnitude of neural responses were found with the acceleration (viewed on a log scale) of white-to-black pixel transitions (Fig. 2; Figs. Extended Data Figs. 6–9), representing the edge acceleration in the moving, expanding, or contracting stimulus observed by the locust. The results support the idea that the presence of feathered surfaces on forelimbs and tail performing accelerated movements, even within a limited motion range typical for early pennaraptorans[1,4,5], could enhance stimulation of sensory-neural circuits in prey receivers and could benefit flush-pursuing dinosaurs (see SI Part 6a, b). This finding is consistent with the association between visual stimulus acceleration and DCMD responses[25,26]. Looming-sensitive fast-escape circuits with properties generally similar to the LGMD/DCMD pathway are known across a wide range of animal taxa[27–31]. Therefore, our results could be expanded to the evolution of proto-wing and tail surfaces in flush-pursuing pennaraptorans that performed flush-displays toward diverse prey taxa, confirming the feasibility of this component of visual display hypothesis.

### Part 4: Results of Experiment 4 with walking before display

Experiment 4 added a 'walking' stimulus prior to the flush display (Extended Data Fig. 5; Video S6; display from Experiment 1). While the presence of feathers still increased the peak spiking rates (Extended Data Fig. 5a, b, c; Table S9), the added walking stimulus caused an overall decrease in the peak spiking rates (Extended Data Fig. 5c; Table S9). Consequently, the threshold for the initiation of jump preparation was not reached in the walk present treatment, regardless of the presence of feathers. However, for the walk absent treatment, the presence of feathers did increase the probability of reaching the threshold, similar to Experiment 1.

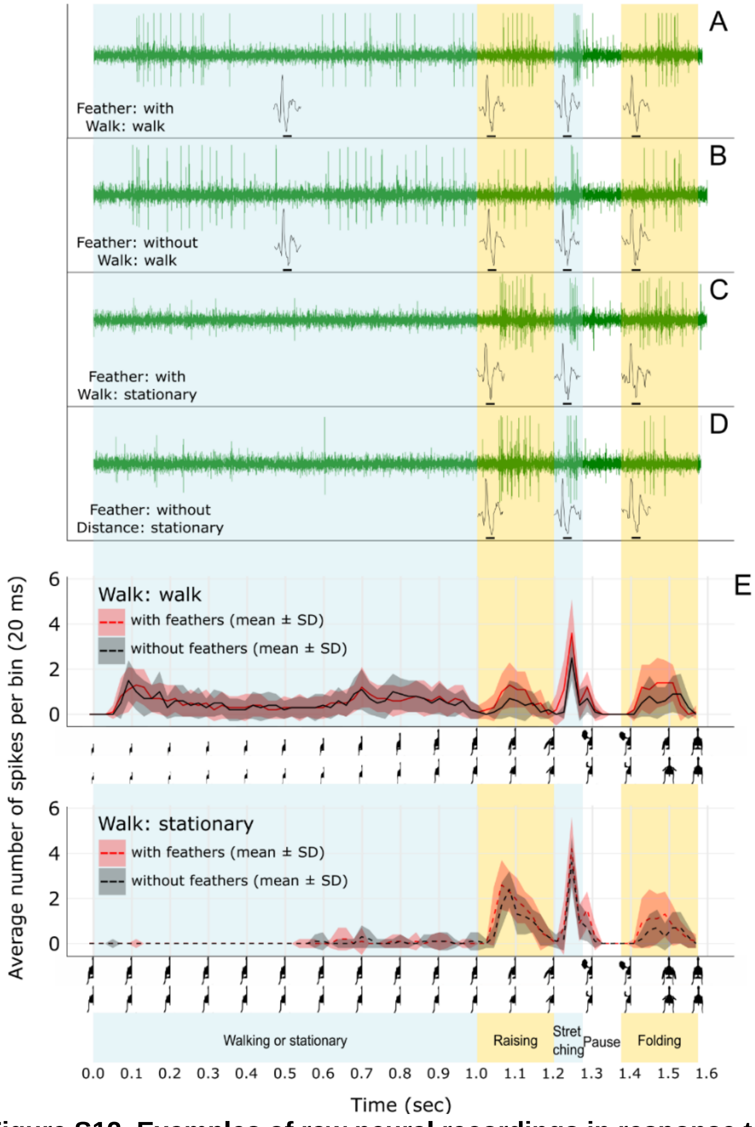


**Figure S12. Examples of raw neural recordings in response to animations imitating the display and walking (1 m/sec) of *Geococcyx californianus* in Experiment 4** "with" (A, C) or "without" (B, D) feathers in animations with "walk" present before display (A, B) or in animations of "stationary" dinosaurs (C, D). Single spike shapes within each colored box represent an example from the recording corresponding to that particular display phase. The four recordings in A–D) are from the same individual. (E) Average spiking rate of the locust's LGMD/DCMD escape pathway (no. spikes/20 ms; mean ± SD; n = 24, i.e., four recordings from each of six individuals) in response to animations "with" (red) and "without" (black) feathers. The upper panel in E represent the case where walking stimulation (1 m for 1 sec) is included, while lower panels represent the case where the dinosaur model remains stationary at 70 cm to prey until it begins the forelimb display. Time (sec) is plotted on the X-axis, with corresponding half-screenshots below. Display phases are indicated by semitransparent vertical boxes.

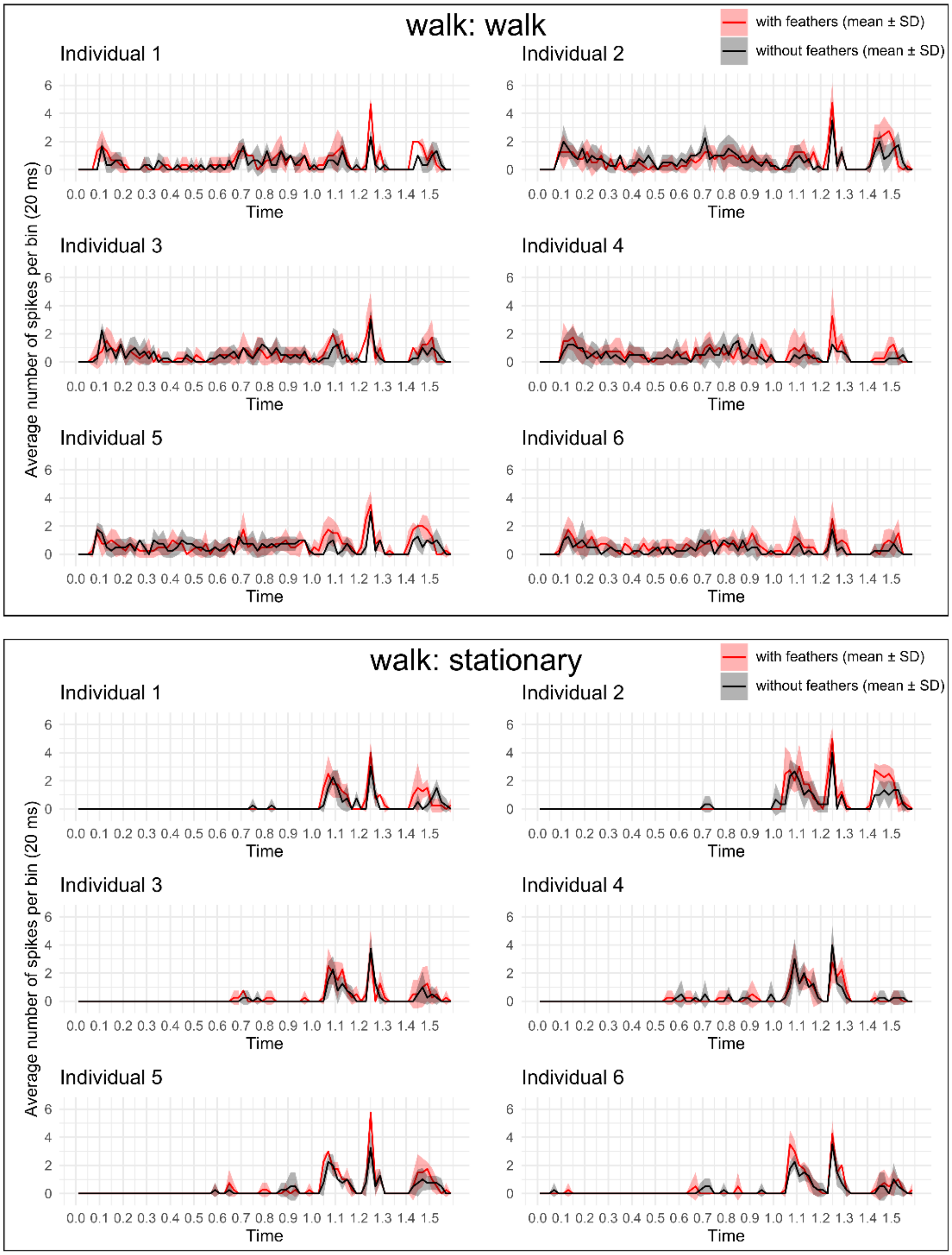


**Figure S13. Average spiking rate per bin (no. spikes/20 ms; mean ± SD) of the locust's LGMD/DCMD escape pathway for each individual, responding to animations imitating the display of *Geococcyx californianus* and a walking stimulus of 1 m/sec in Experiment 4.** Each small panel represents the spiking rate (n = 4 records) in response to animations "with" (red line) or "without" (black line) feathers. Shading represents 95% confidence intervals. The X-axis represents time. The upper box illustrates the spiking rate for animations with the 'walk' phase (1 m for 1 sec), while lower panels represent the case where the dinosaur model remains stationary until it begins the forelimb display at 70 cm to prey.

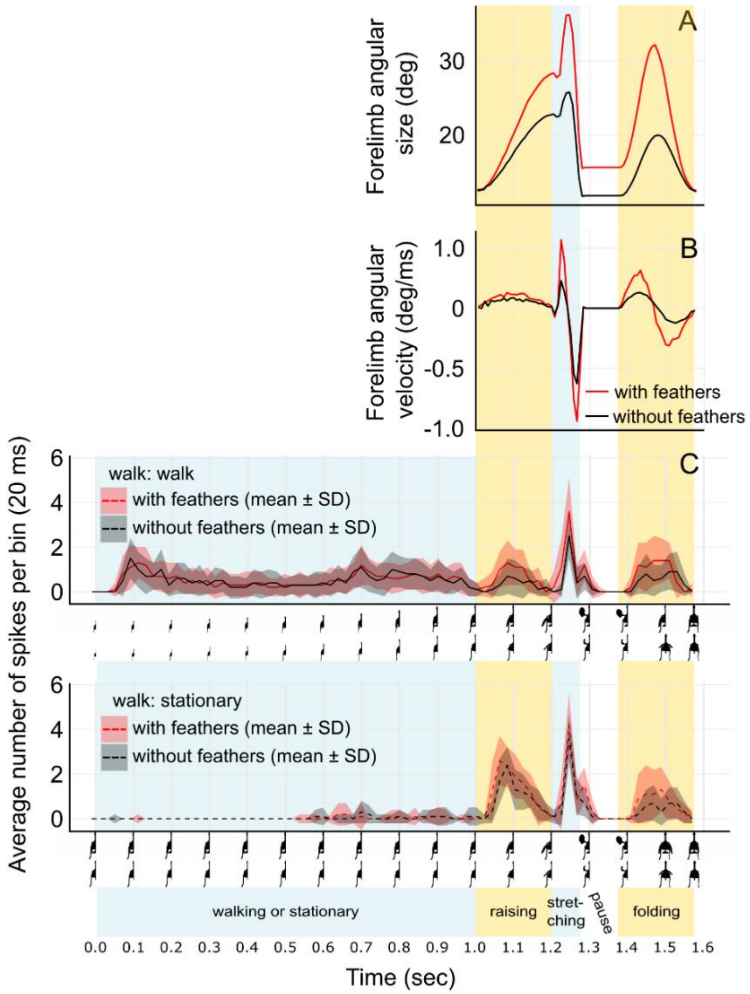


**Figure S14. Profiles of angular size and angular velocity based on forelimb tips in animations, and DCMD responses of locusts to those animations, imitating the display of *Geococcyx californianus* in Experiment 4 by dinosaurs "with" (red line) or "without" (black line) feathers in animations imitating a "walk" before display or a "stationary" display.** The forelimb display phases remain identical after the walking or stationary phase. (A) Angular size (deg) of the distance between the forelimb tips. (B) Angular velocity (deg/ms) based on (A). (C) Average spiking rate of the locust's LGMD/DCMD escape pathway (no. spikes/20 ms; mean ± SD; n = 24, i.e., four recordings from each of six individuals) in response to animations with (red) and without feathers (black). Upper panel in C represent the case where walking stimulation (1 m for 1 sec) is included, while lower panels represent the case where the dinosaur model remains stationary until it begins the forelimb display at 70 cm to prey. Time (sec) is plotted on the X-axis, with corresponding half-screenshots below. Display phases are indicated by semitransparent vertical boxes.

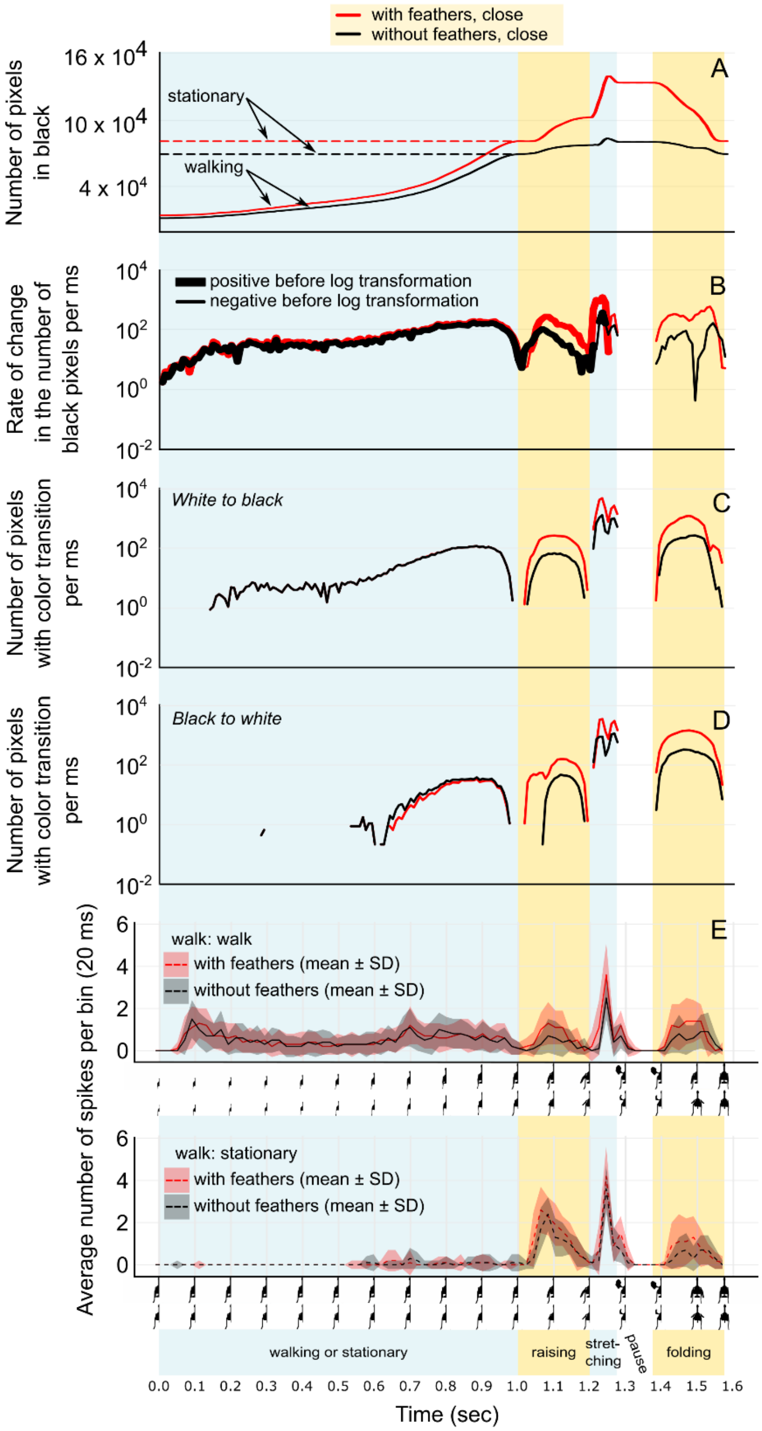


**Figure S15. Profiles of four variables concerning visual expansion of dinosaur silhouette and edge movements extracted from the animations (A–D), and DCMD responses (E) to those animations imitating the display of *Geococcyx californianus* in Experiment 4 by dinosaurs "with" (red line) or "without" (black line) feathers in animations imitating a "walk" before display or just a "stationary" display.** (A) Number of pixels in black per frame. (B) Rate of change in the number of black pixels per ms based on change observed between consecutive frames. Positive values before applying the log scale to the Y-axis are shown as thick lines, while negative values are represented as thin lines. (C) Rate of white-to-black transition in the stimulus represented by the number of pixels with white-to-black color transition per ms based on the number of pixels with color transition observed between two consecutive frames during animation. (D) Rate of black-to-white transition in the stimulus represented by the number of pixels with black-to-white color transition per ms based on the number of pixels with color transition observed between two consecutive frames during animation. (E) Average spiking rate per bin (no. spikes/20 ms; mean ± SD; n = 24, i.e., four recordings from each of six individuals) of the locust's LGMD/DCMD escape pathway in response to animations "with" (red) and "without" feathers (black). The upper panel in E represents the imitation including walking, while lower panels represent the stationary dinosaur animation. Time (sec) is plotted on the X-axis, with corresponding half-screenshots below. Display phases are indicated by semitransparent vertical boxes. For panels B, C, and D, a logarithmic scale was applied to the Y-axis for better visualization.

### PART 5: Supplementary Tables with results (Tables S5–S9)

**Table S5. Number of neural recordings that reached the threshold of 5 spikes/20 ms in the experiment involving the display of *Geococcyx californianus* in Experiment 1.** The numbers in each box represent the count of records reaching the threshold value, with a maximum value of 24 in response to four animation types. This table concerns Fig. 1e.

| Display phase | Threshold reaching | Animation type (distance option, feather option) | | | |
|---|---|---|---|---|---|
| | | Close, With | Close, Without | Distant, With | Distant, Without |
| Raising | Reached | 14 | 6 | 6 | 2 |
| | Not reached | 10 | 17 | 17 | 21 |
| Stretching | Reached | 7 | 4 | 10 | 5 |
| | Not reached | 14 | 19 | 13 | 18 |
| Folding | Reached | 0 | 0 | 0 | 0 |
| | Not reached | 24 | 23 | 23 | 23 |

**Table S6. Number of neural recordings reaching the threshold of five spikes/20 ms in the experiment involving the display of *Cercotrichas galactotes* in Experiment 2.** The numbers in each box represent the count of records reaching the threshold value, with a maximum value of 24 in response to four animation types. This table concerns Fig. 1l.

| Display phase | Threshold reaching | Animation type (distance option, feather option) | | | |
|---|---|---|---|---|---|
| | | Close, With | Close, Without | Distant, With | Distant, Without |
| Third tail hitch | Reached | 18 | 13 | 0 | 0 |
| | Not reached | 6 | 11 | 23 | 21 |
| Fourth tail hitch | Reached | 7 | 4 | 0 | 2 |
| | Not reached | 17 | 20 | 23 | 19 |
| Lowering tail & spreading forelimbs | Reached | 20 | 2 | 14 | 0 |
| | Not reached | 4 | 22 | 9 | 21 |
| Folding forelimbs | Reached | 1 | 0 | 0 | 0 |
| | Not reached | 23 | 24 | 23 | 21 |

**Table S7. Number of neural recordings reaching the threshold of five spikes/20 ms in the experiment involving the display of *Mimus polyglottos* in Experiment 3.** The numbers in each box represent the count of records reaching the threshold value, with a maximum value of 24 in response to four animation types. This table concerns Fig. 1s.

| Display phase | Threshold reaching | Animation type (distance option, feather option) | | | |
|---|---|---|---|---|---|
| | | Close, With | Close, Without | Distant, With | Distant, Without |
| First hitch | Reached | 13 | 10 | 6 | 1 |
| | Not reached | 11 | 14 | 18 | 23 |
| Second hitch | Reached | 3 | 0 | 1 | 0 |
| | Not reached | 21 | 24 | 23 | 24 |
| Third hitch | Reached | 8 | 10 | 2 | 1 |
| | Not reached | 16 | 14 | 22 | 23 |
| Folding | Reached | 1 | 0 | 0 | 0 |
| | Not reached | 23 | 24 | 24 | 24 |

**Table S8. Number of neural recordings reaching the threshold of 5 spikes/20 ms in the experiment involving the display of *Geococcyx californianus* and a walking stimulus of 1 m/sec in Experiment 4.** The numbers in each box represent the count of records reaching the threshold value, with a maximum value of 24 in response to four animation types. This table concerns Extended Data Fig. 5e.

| Display phase | Threshold reaching | Animation type (walk option, feather option) | | | |
|---|---|---|---|---|---|
| | | Stationary, With | Stationary, Without | Walk, With | Walk, Without |
| Raising | Reached | 3 | 0 | 0 | 0 |
| | Not reached | 21 | 23 | 23 | 23 |
| Stretching | Reached | 12 | 4 | 7 | 0 |
| | Not reached | 12 | 19 | 16 | 23 |
| Folding | Reached | 0 | 0 | 0 | 0 |
| | Not reached | 24 | 23 | 23 | 23 |

**Table S9. Generalized mixed-effects model for analyzing peak size and threshold variations in neural responses to dinosaur animations imitating the display and walking of *Geococcyx californianus* in Experiment 4.** For the under-dispersed count of peak size (no. spikes/20 ms), we utilized the Conway-Maxwell poisson family with a log link function. For the binary threshold (reached = 1, unreached = 0), we used the Binomial distribution with a logit link function. The fixed effects include feather (with or without), walk (walk or stationary), the interaction term of feather and distance, and display phase (raising, stretching, or folding). Random effects include individual ID and record ID nested within individual ID. In response to the 'raising' display phase, only three records reached the threshold (5 spikes/20 ms[16]) for jump preparation, all of which responded to animations featuring feathers and no walking (Table S8). Moreover, none of the responses to the 'walking-non-feathered' animation reached the threshold in the 'folding' display phase (Table S8). Consequently, when analyzing threshold variation, we only considered records responding to the 'stretching' display phase without walking. The table values correspond to the effect estimate, standard error, Z value, and P-value for each variable. P-values below 0.05 are denoted in bold, except for the intercept. This table concerns Extended Data Fig. 5.

| Response variable | Fixed effect | Estimate | Standard Error | Z value | P-value |
|---|---|---|---|---|---|
| Peak size | Intercept | 1.046 | 0.084 | 12.498 | < 0.001 |
| | **Feather** *'without'* | -0.214 | 0.073 | -2.940 | **0.003** |
| | **Walk** *'walk'* | -0.177 | 0.072 | -2.457 | **0.014** |
| | Feather *'without'*:walk *'walk'* | -0.179 | 0.109 | -1.638 | 0.101 |
| | **Display phase** *'stretching'* | 0.427 | 0.049 | 8.661 | **< 0.001** |
| | **Display phase** *'folding'* | -0.347 | 0.061 | -5.658 | **< 0.001** |
| Threshold | Intercept | 0.009 | 0.586 | 0.015 | 0.988 |
| | **Feather** *'without'* | -1.781 | 0.774 | -2.303 | **0.021** |

### PART 6. Details of visual display hypothesis and review of sauropsid visual displays

#### 6A) Hypothetical flush-displays in Pennaraptora

The results support the idea that the presence of feathered surfaces on forelimbs and tail performing accelerated movements, even within the anatomically limited motion range for typical early pennaraptorans[1,4,5], could enhance stimulation of sensory-neural circuits in various prey receivers, which would lead to higher frequency of prey escapes and fuel the natural selection for flush-displays. The neural response patterns approximately matched the acceleration pattern in the rate of white-to-black pixel transitions in the stimulus, representing accelerated edge movements in looming or translation of the stimulus. It appears that those variables have higher values in the relatively simple displays (Fig. 2a, b) compared to the relatively complex and stereotyped mockingbird-based display performed within the narrow range allowed by pennaraptoran anatomy (Fig. 2c). Additionally, when the walking stimulus was included prior to a display (SI Part 4), the peaks of neural responses were generally reduced compared to the non-walking animations (possibly due to habituation with prolonged exposure to walking visual stimuli[32]). Therefore, it would have been beneficial that flush-displays comprise relatively simple extensive movements after a period of near-motionless posture. As those efficient flush-movements seem to loosely resemble prey-grabbing attempts with forelimbs or breaking of moving body with both proto-wings spread outward, this brings an idea that the of flush-display component of the visual display hypothesis may have started with the benefit from accidental flushing of focal prey by forward/outward moving forelimbs followed by fast neck/head movements to intercept the prey’s trajectory with mandibles to capture it orally (similar to the roadrunner behavior; Video S1).

#### 6B) Reinforcing natural selection mechanisms of the flush-display component of the visual display hypothesis

Unlike other components of the visual display hypothesis (Fig. 3d), the flush-display component uniquely incorporates a reinforcing cycle of natural selection for flush-display effectiveness (Fig. 3b, c) and for pursuit success (multiple mechanisms in Fig. 3a). Once a prey-pursuer employs flush-displays (Fig. 3b), the frequency of prey-pursuit events may increase, which could drive stronger natural selection for improved pursuit and capture efficiency (functions 1–7 in Fig. 3a). This selection pressure may lead to the evolution of larger and stiffer proto-wings and tail feathers, aiding functions 1–7. These enlarged feather areas are likely to increase flush frequency (prey flush function; Fig. 3b) due to the prey escape pathways’ sensitivity to enhanced visual stimuli (Fig. 3c). As the same traits may be beneficial for signaling (Fig. 3b) and non-signaling (fig. 3a) functions, we hypothesize that this relationship between flush and pursue functions may create a reinforcing cycle of natural selection for traits (e.g., proto-wing size and forelimb anatomy) that enhance flush-display efficiency (Fig. 3b) as well as pursuit and capture efficiency[1] (Fig. 3a).

Some of the functions in Fig. 3a (functions 3, 4, and 7) invoke hypothetical aerodynamic benefits from flapping proto-wings. However, they probably did not have a crucial influence on the acquisition of pennaceous feathers[33–35] because integuments and pectoral girdle anatomy in non-paravian pennaraptorans are inconsistent with flapping-based functions of proto-wings[36]. Similarly, wing-assisted incline running[37,38] (function 7 in Fig. 3a) is more plausible in Ornithothoraces[33,34] with greater shoulder mobility than earlier-diverging pennaraptorans. Hence, it was proposed[33,35] that non-locomotory or less stringent locomotory behaviors—such as balancing or braking—may have played a role in the origin of pennaceous proto-wings and tail, which is expected to happen once at the stage of basal Pennaraptora[15]. These include drag-based (functions 1 and 2) and mechanical (functions 5 and 6) functions of pennaceous feathers on limbs and tail. The original "insect netting" function[39], and its modified form of "insect cupping"[40] (function 5 in Fig. 3a), should not be ignored because some birds use wings to capture food items[40]. The criticism suggesting that early pennaceous feathers would not allow sufficient air passage to function as efficient capturing/knocking down net-like devices[41] has not been empirically tested.

Hence, not all functions in Fig. 3a pose the same potential for shaping the function and the initial evolution of pennaceous proto-wings and tail in early pennaraptorans. Regardless of particular non-signaling functions involved, the hypothetical reinforcing process between natural selection for non-signaling and for visual-signaling functions (Fig. 3b, c) is consistent with elevated rates of the relevant anatomical evolution along the bird stem lineage[42,43]. Flush-display component might have been indeed used, in addition to other components, by the ancestors of early Pennaraptora because these ancestors along the bird-stem lineage were omnivorous[27] just like many modern avian flush-pursuers.

#### 6C) Visual display hypothesis

The visual display hypothesis is artistically depicted in Fig. S16. The hypothesis proposes that in each species the multiple signaling functions (Fig. 3d) interact with one another in shaping the feathered display surfaces and forelimb/tail movements, leading to the observed diversity[33] and inferred multifunctionality of the exaggerated motion-based visual displays by proto-wings and tail observed in Pennaraptora. This hypothesis is consistent with the widespread use of motion-based visual signals across a range of contexts in diverse animal taxa, as illustrated for extant Sauropsida in a brief summary (Table S10), and in examples presented in SI Parts 6E, F. The review illustrates that in principle the motion-based visual displays could be based on motions of various body parts: head, neck, back, tail, forelimbs, and hindlimbs (SI Part 6E). However, the most extensive movements are possible with forelimbs and tail owing to their length and therefore forelimbs and tail motions are most likely to deliver strong motion-based signals.

The multiple components of the visual display hypothesis are schematically shown in Fig. 3d. Based on the observations in extant Sauropsida (SI Part 6D, E, F), these components represent various motion-based visual signaling contexts and different functions within each signaling context. In the context of signaling towards prey, visual displays may function as flushing, prey-luring, and deflection during attacks on dangerous prey (e.g., poisonous snakes). In the context of signaling towards predators, visual displays may function as startle/deimatic, pursuit deterrence, aposematic, mobbing, parental distraction displays that lead predators away from offspring, and deflection displays during attacks by predators. In the context of signaling towards social group members, visual displays may function to establish dominance and resolve aggressive conflicts with conspecific and heterospecific members of a social group. In the context of signaling towards parents and offspring caretakers in general, visual displays may aid in begging for food. In the context of signaling towards mating competitors, visual displays serve a function of resolving within-sex contests in competition for mates (e.g., competition among males to establish territories and to obtain mates) and may resemble the aggressive/dominance displays in social groups. Finally, in the context of signaling towards prospective and actual mates, visual displays could be a part of courtship behavior. Begging between mates can also be accompanied by motion-based displays.

The review (SI Part 6D, E, F) shows that within each species, the same surfaces—such as forelimbs and tail in reptiles or wings and tail feathers in birds—can be used in different signaling contexts, while movements employed may differ among different signaling contexts. For example, lizards use various arm waving or tail movements in socio-sexual (toward mates and toward competitors) and antipredatory contexts (e.g., paper 19 in SI Part 6F). Among birds, roadrunners use flashes of white spots on open wings in flush-displays, wing flapping while attacking rattlesnakes, wing fluttering during in distractions displays to predators, tail wagging from side to side and wing popping during male-male interactions, displays by males to females of lifted wings and tail followed by lowering the wings and bringing them close to the body with a "pop" sound, to which females respond through rapid flicks in vertical plane[14]. Generally, the diverse avian passerines' visual motion-based displays involving tail spreading, gliding with tail spreading and shallow wingbeats, body-pivoting with spread tail, wing drooping, wings-out (horizontally lifted), and wing/tail spreading and quivering are typically used in multiple different signaling contexts within each species: courtship, mate competition, flush-displays, and displays towards predators (e.g., papers 32, 35 in SI Part 6F).

Based on these examples and the evidence from the literature review of visual displays in Sauropsida (SI Part 6D–F), it is likely that the visual display hypothesis comprising multiple functions of early pennaceous feathers (Fig. 3d) may have shaped the pennaceous proto-wings and tail in early Pennaraptora.

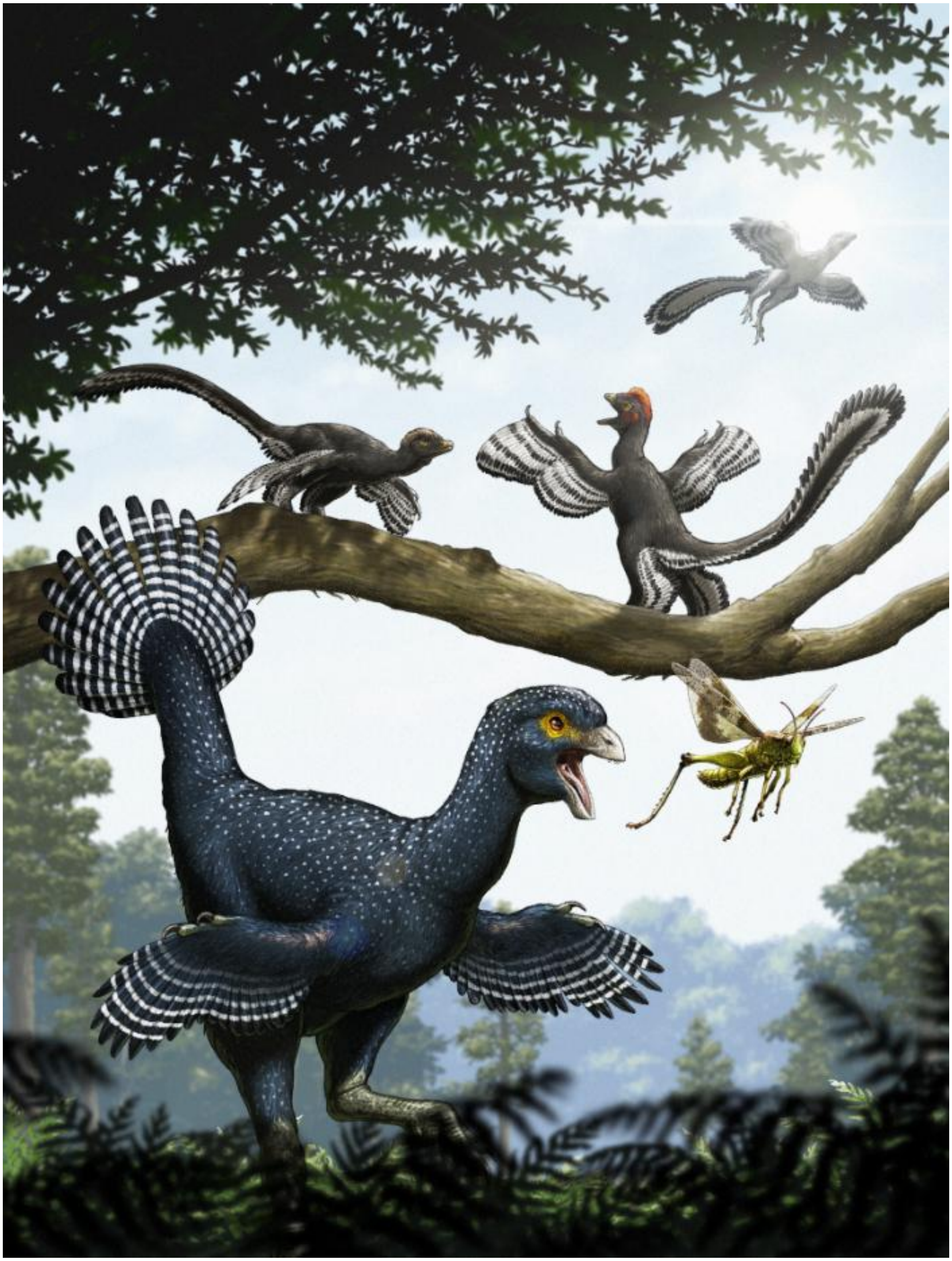

**Figure S16. Artistic reconstruction of displaying dinosaurs with pennaceous feathers.**
© Choi Yu-sik.

#### 6D) Brief overview of motion-based signals in Sauropsida

**Table. S10. Brief overview of motion-based visual signals, focusing on movements with limbs and tail in extant Sauropsida.** The table is based on general knowledge from animal ethology as well as information contained in scientific literature, some of which are listed below. The table focuses on movements of limbs and tail, but movements of head/head ornaments, neck (dewlap), body movements and shape/size changes are also common.

| **Visual displays in Sauropsida** | Signal categories | tuatara, geckos, lizards, and snakes (Squamata) | turtles & tortoises (Testudines) | crocodilians (Crocodilia) | birds (Aves) |
|---|---|---|---|---|---|
| Properties of visual signals | increase in surface | + | - | + | + |
| | color contrast | + | + head | - | + |
| | visual motion | + | + head | + | + |
| Functions of visual signals | flush-pursue (towards prey) | ? tail movements in one species | - | - | + |
| | antipredator | + | + plastron colors | - | + |
| | social signals | + | + | + | + |
| | sexual signals | + | + | + | + |
| | offspring-parent (e.g., begging) | - | - | - | + |
| Limb and tail displays | tail: apparent size increase and/or color | + | - | + | + |
| | tail motion | + | - | + | + |
| | limb: apparent size increase and/or color | + | + foreclaw display | - | + |
| | limb motion | + | + foreclaw display | + | + |

### 6E) Examples of various descriptions of display motions in Sauropsida, excluding birds

This information is extracted from the “List of papers” in SI Part 6E

**Head**
bobbing of the head vertically
crest erection
crest raising
head-shuddering
lateral headshaking
head nods
opening of head flaps
lip smacking
vertical vibration, horizontal swinging
rising head and exposing dorsum,

**Neck**
lateral compression and extension of dewlap

**Back**
back arching
crest erection
crest raising

**Body lateral surfaces**
body inflation
body compression/inflation laterally
inflated posture

**Body movements**
Pushups
forward body thrust
body raising
body tilting
body-rock display

**Forelimb and hindlimb**
limb-waving
backward arm-wave, forward arm-wave,
limb extension and retraction
leg flexing
arm waving
foreclaw display (in turtles)
upward extension of either the front two or all four limbs

**Tail**
tail-flicking
tail-coiling/raising
tail-lashing,
tail trashing
tail luring (to deflect attack)
tail waving
tail wagging
lateral tail undulation
elevation of the straightened tail + moving it back and forth laterally = lateral wave
sinusoidal motion = sinusoidal wave
tail curl involves slowly raising and curling the distal portion of the tail
tail raise involves rapidly elevating the tail and holding it stiffly
undulation of the tail from side to side in gentle sweeping or rapid twitching motions
tail inflating
tail arched posture,

#### 6F) Examples of papers on Sauropsida with descriptions of visual displays

**Examples of papers on Sauropsida containing:**

- **the use of movements in visual displays, with a focus on wing and tail movements (marked bold underline)**
- **the determination of the visual signaling context**
- **estimates of the speed and acceleration of visual displays**
- **the effects of environment on motion-based visual displays**

The short descriptions below provide most relevant information

##### Tuatara, Geckos, lizards, and snakes (Order: Squamata)

***1)*** *Baird T.A. 2013. Lizards and other reptiles as model systems for the study of contest behaviour. CH 12 in "Animal Contests" eds. I.C.W. Hardy and M. Briffa. Cambridge University Press.* – Review of aggressive behaviors in reptiles, with a focus on lizards. Lists the following visual motion-based displays: lateral compression and extension of dewlap (both increase in apparent size), bobbing of the head vertically, pushups, **thrashing the tail**, "**circumduction"** of the **front limbs** **(i.e., "arm-waving")**. Many species show colors during those movements.

***2)*** *Bian X., Pnilla A., Chandler T., Peters R. 2021. Simulations with Australian dragon lizards suggest movement-based signal effectiveness is dependent on display structure and environmental conditions. Scientific Reports (2021) 11:6383, https://doi.org/10.1038/s41598-021-85793-3* – This is a study of motion-based displays of four species of agamid lizards evaluated through an artificial neural network that estimates the dynamics of relative saliency of display recreated on the computer screen in 3D animation against different habitats' background. The following distinct visual display elements were considered: **tail-flicking**, **limb-waving**, push-ups, head bobbing, forward body thrust, and **tail-coiling/raising**.

***3)*** *Bian X., Zhao W., Qi W., Peters R. 2025. Tail tales: how ecological context mediates signal effectiveness in a lizard. Integrative Zoology, 0, 1-16, https://doi.org/10.1111/1749-4877.12943.* – The effectiveness of two elements of visual **socio-sexual** displays were compared, **tail-coiling** and **tail-lashing**, depending on the features of the **visual background** of the environment.

***4)*** *Cooper W.E., Perez-Mellado V., Baird T.A., Caldwell J.P., Vitt L.J. 2004. Pursuit deterrent signaling by the Bonaire whiptail lizard Cnemidophorus murinus. Behaviour 141, 297-311.* – The paper contains a description of **arm-waving towards a potential predator** (human observer) and suggests a pursuit-deterrence function.

***5)*** *Fleishman L.J. 1988. Sensory influences on physical design of a visual display. Animal Behaviour 36, 1420-1424.* – Long-range displays especially contained movements of extremely **high acceleration and velocity**.

***6)*** *Gillingham J.C., Carmichael C., Miller T. 1995. Social behavior of the tuatara sphenodon punctatus. Herpetological Monographs 9, 5-16.* – Description of **socio-sexual** visual displays including body inflation and **crest erection**, head-shuddering, **lateral headshaking**, and head nods. The radio-controlled life-sized tuatara model revealed the importance of nuchal and dorsal crest displays. Although no arm or tail waving was observed, the crest erection may be viewed as an equivalent of the hypothetical proto-wings/tail spreading in pennaraptorans, and lateral head shaking may produce visual stimuli generally similar to later tail movements.

***7)*** *Hagman M., Phillips B.K., Shine R. 2008/ Tails of enticement: caudal luring by an ambush-forging snake (Acanthopis praelongus, Elapidae). –* Visual **tail movements** are used as signals **towards prey** to lure the prey within striking distance.

***8)*** *Jenssen T.A. 1977. Evolution of anoline lizard display behavior. American Zoologist 17, 203-215. –* The paper contains hypothetical evolutionary scenarios from the origin of simple visual motion-based displays to the development of displays with multiple elements in anoline lizards.

***9)*** *Miranda R.B., Klaczko J., Tonini J.F.R., Brandao R.A. 2022. Escaping from predators: a review of Neotropical lizards defense traits. Ethology Ecology & Evolution [https://doi.org/10.1080/03949370.2022.2082538](https://doi.org/10.1080/03949370.2022.2082538) –* Review and phylogenetic analyses based on data from 206 papers indicate multiple origins of various antipredatory behaviors, including visual displays. From among the various visual displays, the following have been mentioned as potentially serving antipredatory functions and involve tails or forelimbs displayed **towards predators**: **tail luring** (with colored tail) to deflect the predatory attack, **limb extension and retraction**, **tail waving**, **leg flexing**, possibly as pursuit deterrent signals.

***10)*** *Ord T.J., Blumstein D.T., Evans C.S. 2001. Intrasexual selection predicts the evolution of signal complexity in lizards. Proceedings of the Royal Sciety London B 268 737-744. –* Review and phylogenetic analyses show an association between the evolution of sexual dimorphism and the visual **socio-sexual display** richness in agamid and iguanid lizards. The review included the following elements+ of visual displays: head-nod, push-up display, dewlap extensions, **tail wagging**, **arm waving**, crest raising, body compression/inflation, back arching, body raising/tilting, and changes in color.

***11)*** *Ord T.J., Blumstein D.T., Evans C.S. 2002. Ecology and signal evolution in lizards. Biological Journal of the Linnean Society22, 127-148. –* Review and phylogenetic analyses for agamid and iguanid lizards from 141 literature sources. The paper lists the following elements of visual displays: head-nod display, push-up display, back arching, **arm waving**, body compression/inflation, body raising, body tilting, body color changes, lip smacking, crest raising, **tail displays**, throat displays (dewlap extensions).

***12)*** *Ord, T.J., Peters, R.A., Clucas, B. & Stamps, J.A. 2007. Lizards speed up visual displays in noisy motion habitats. Proc. R. Soc. B Biol. Sci. 274: 1057–1062. –* The paper illustrates how **environmental conditions** affect the design of motion-based **visual displays** in a **socio-sexual** context.

***13)*** *Ossip-Drahos A.G., Berry N.J., King C.M., Martins E.P. 2018. Information-gathering as a response to manipulated signals in the eastern fence lizards, Sceloporus undulatus. Ethology, 124, 684-690. –* **Experimental manipulation of color patch size** that is used in visual displays during agonistic/territorial interactions demonstrated that the larger surface area of the patch affects the reaction of the competitor in a manner consistent with the **larger patch** representing a **clearer signal of dominance**.

***14)*** *Peters R.A., Cliffords C.W.G., Evans C.S. 2002. Measuring the structure of dynamic visual signals. Animal Behaviour 64, 131-146. –* Video sequences of socio-sexual displays by the Jacky dragon have been analyzed using computational motion analysis to estimate optic flow and produce optic flow plots. The following elements of the lizard's **socio-sexual** visual displays have been observed: **tail-flick**, **backward arm-wave**, **forward arm-wave**, push-ups, and body-rock display, which features movement up and to the side as the lizard raises its body from the substrate, and down as it returns.

***15)*** *Peters R.A., Evans C.S. 2003. Design of the Jacky dragon visual display: signal and noise characteristics in a complex moving environment. Journal of Comparative Physiology A, 189, 447-459. –* This paper contains **speed-time profiles** for some of the visual displays. The speed time profiles (e.g. Fig. 8 in this paper) illustrate that the visual displays in a **socio-sexual** context comprise movements of **high speed and acceleration** that **stand out from the movements present in the environment**.

***16)*** *Peters R.A., Ord T.J. 2003. Display response of the jacky dragon, Amphibolurus muricatus (Lacertilia: Agamidae), to intruders: a semi-Markovian process. Austral Ecology 28, 499-506. –* Results illustrate social visual display to conspecifics and include backward **arm-wave**, forward **arm-wave**, **tail-flicks** as well as push-ups, and 'body-rock" display.

***17)*** *Ramos J.A., Peters R.A. 2017. Motion-based signaling in sympatric species of Australian agamid lizards. Journal of Comparative Physiology A, 203, 661-671. –* Examples of speed-time plots and angular speed distributions of visual displays illustrate that displays contain **high-speed** and **high-acceleration** movements.

***18)*** *Regaldo R. 2003. Roles of visual, acoustic, and chemical signals in social interactions of the tropiclal house gecko (Hemidactylus mabouia). Caribbean Journal of Science, 39: 307-320. –* Results show that tropical house geckos use arched-back and body-raised behavior for increasing visual size towards same-sex or opposite-sex individuals. **Tail wave** and **tail trashing** are also used in **social interactions** by both sexes.

***19)*** *Senter P.J. 2024. Antipredator displays by the Cuban groundlizard (Pholidoscelis auberi) on Little San Salvador Island (Half Moon Cay), the Bahamas, and a review of similar behavior in other lizards. Reptiles & Amphibians 31, e22527. –* **Arm waving** and **lateral undulation of tails** displays were observed in the focal species in the contexts indicating **antipredatory function**. A **review of the literature** revealed that **arm waving** display was described in **socio-sexual contexts** in agamid, iguanid, lacertid, liolaemid, phrynosomatid, and skink lizards as well as in geckos. Specific details of the arm waving vary from species to species. The review also revealed that **lateral tail undulation** displays in **response to predators** or humans were observed in teiids, lacertids, varanids, crotaphytids, phrynosomatids, sphaerodactylids, and skinks. It was observed as part of **socio-sexual displays** in gekkonids, lacertids, and phrynosomatids. It was also used in **displays towards prey** (prey luring displays).

***20)*** *Whiting M.J., Noble D.W.A., Qi Y. 2022. A potential deimatic display revealed in a lizard. Biological Journal of the Linnean Society, 136: 455-465. –* Results show that opening of **head flaps** in response **to predators** serves as a deimatic/deterrence signal to **predators** in agamid lizards. This may be viewed as **equivalent to sudden proto-wing expansion**/tail opening in the hypothetical pennaraptoran.

***21)*** *Woo K.L., Rieucau G. 2011. Aggressive signal design in the jacky dragon (Amphibolurus muricatus): display duration affects efficacy. Ethology 118 157168. –* This is an experimental study with video playbacks to study push-up and **tail-flick** responses in lizards in the context of **socio-sexual** displays.

***22)*** *York J.R., Baird T.A. 2016. Juvenile collared lizards adjust tail display frequency in response to variable predatory threat. Ethology. 122, 37-44. –* Tail motion used **towards predators** apparently as a pursuit deterrent signal. Part of Results: "Juvenile collared lizards performed four variations of displays using their tails. The most common display involved the conspicuous **elevation of the straightened tail** above the substrate and **moving it back and forth laterally (= lateral wave**). Lizards are also frequently displayed by **elevating the tail** and moving it using a stereotypical slow and deliberate **sinusoidal motion (= sinusoidal wave**, Fig. 2b). Less frequently, lizards **raised and held the tail stiffly** in two different positions. **Tail curl** (shown just beginning in Fig. 2c) involved **slowly raising and curling the distal portion of the tail** until it was directly above the head and pointed anteriorly. Tail raise involved **rapidly elevating the tail** and holding it stiffly above the substrate (same as in Fig. 2a, but without lateral movement)".

#### Turtles & tortoises (Order: Testudines)

***23)*** *Liu Y. X., Davy C. M., Shi H. T. Murphy R. W. 2013. Sex in the half-shell: a review of the functions and evolution of courtship behavior in freshwater turtles. Chelonian Conservation and Biology, 12: 84-100. –* Contains review information on **foreclaw display** and **head movements** as part of **socio-sexual** signaling repertoire in multiple species of turtles from several families. While these behaviors produce visual stimuli—such as fast limb movements, temporary increases in apparent body size, and resulting posture changes—they are less dramatic than those observed in some lizards or birds.

***24)*** *Brejcha J,. Kleisner K. 2015. Turtles are not just walking stones: conspicuous coloration and sexual selection in freshwater turtles. Biosemiotics. DOI 10.1007/s12304-015-9249-9 –* Contains descriptions of the association between color on head and use of the head in signaling in a sexual and social context. Describes the use of **head movements** (vertical vibration, horizontal swinging) and **forelimb movements** ("foreclaw display") in **socio-sexual signaling**.

***25)*** *Britson C.A. 1998. Predatory responses of largemouth bass (Micropterus salmoides) to conspicuous and cryptic hatchling turtles: a comparative experiment. Copeia 2: 385-390. –* Contains experimental determination of **aposematic** signaling function of **colors on plastron** as signals **towards predators**.

#### Crocodilians (Order: Crocodilia)

***26)*** *Garrick L.D., Lang J.W., Herzong H.A. 1978. Social signals of adult American alligators. Bulletin of the American Museum of Natural History 160, 153-192. –* Detailed description of behaviors in **socio-sexual** context, which include **rising head** and exposing dorsum, **tail raised** upward out of water, **tail and head lifting**, body compressed laterally, and **tail wagging.**

***27)*** *Garrick L.D., Lang J.W. 1977. Social signals and behaviors of adult alligators and crocodiles. American Zoologist 17, 225-239. –* This paper contains results of solid descriptions of signaling in 3 species. Among the various elements of visual displays in a **socio-sexual** context the following may be relevant to the "visual displays" hypothesis discussed in our paper: **tail arched** posture, inflated posture (increase in apparent size), **tail wagging**.

***28)*** *Brien M.L., Lang J.W., Webb J.W., Webb H.J. Stevenson C., Christian K.A. Brien ML, Lang JW, Webb GJ, Stevenson C, Christian KA (2013) The Good, the Bad, and the Ugly: Agonistic Behaviour in Juvenile Crocodilians. PLoS ONE 8(12): e80872. doi:10.1371/journal.pone.0080872 –* Observations of 7 species confirmed the following elements of **social** visual displays that involve tail and forelimbs: head and **tail raising**, **tail wagging** (undulation of the tail from side to side in either a **gentle sweeping** motion or **rapid twitching**, often repeated several times), **inflated tail,** inflated posture with **upward extension** of either the front two or all four **limbs**.

***29)*** *Vliet K.A. 1989. Social displays of the American alligator (Alligator mississipiensis). American Zoologist 29, 1019-1031. –* observations reveal visual displays that include the following relevant to our question: head oblique **tail arched** posture, inflated posture (to increase apparent size), **tail wag** used in **socio-sexual signaling**.

#### Birds (Class: Aves)

***30)*** *Andrew R.J. 1961. The displays given by passerines in courtship and reproductive fighting: a review. Ibis 103a, 549-579. –* An old classic review of displays organized by avian families documenting the wide-spread presence **wing and tail displays** across many families of birds.

***31)*** *Benedict L., Jones H. Robinson S., McEntee J. P. 2024. Phylogenetic analyses support flush-pursuit foraging and flocking behaviors as evolutionary drivers of flash plumage signals in North American passerines –* This analysis establishes that plumage patches that are flashed during movements of the bird and its tail and wing are **under natural selection** for **flush-pursue displays** and under selection for **social visual signaling** among birds in groups of **conspecifics and heterospecifics** (winter flocks).

***32)*** *Billerman, S. M., B. K. Keeney, G. M. Kirwan, F. Medrano, N. D. Sly, and M. G. Smith, Editors (2025). Birds of the World. Cornell Laboratory of Ornithology, Ithaca, NY, USA. https://birdsoftheworld.org/bow/home. –* Contains descriptions of the biology of all species of birds of the world. Each description includes a summary of communication. **Visual displays with tails and wings are known in almost all species**. They include a variety of opening/closing, sidewise, and upward/downward movements at various speeds. From reading the species accounts, and reaching references cited there, it is clear that **movement-based wing and tail displays** function as visual signals within the same species **in different multiple contexts**: **towards**

**mates, towards mate competitors, towards social group members (conspecifics and heterospecific) and predators as well as prey**. While the detailed characteristics of the movements may vary between contexts in a species-specific manner, an increase in conspicuousness/size of the tail or wing will affect signal effectiveness in all contexts.

*33) Dhondt A.A., Kemink K. 2008. Wing-flashing in Northern mockingbirds: antipredator defense? Journal of Ethology 26: 361-365 –* This study compiles observational evidence for **wing-flash** displays used in the presence/towards **predators** in 9 species in the family Mimidae.

*34) Dhondt, A.A. and K.M. Kemink. 2008. Wing-flashing in Northern Mockingbirds: Antipredator defense? Journal of Ethology 26:361–365 –* The study confirms that the flush-pursuing species used **wing-flashing** not only for **flush-pursue** foraging but also as a visual display **towards predators**.

*35) Ficken M.S. 1962. Agonistic behavior and territory in the American redstart. Auk 79, 607-632. –* A classic example of ornithological papers describing visual displays in birds and illustrating the use of visual motion-based displays with wings and tail across different contexts: **courtship**, **mate competition**, **flush-pursue display**, and **display towards predators**. The visual displays described here involve **tail spreading**, gliding with tail spreading and **shallow wingbeats**, **body-pivoting with tail spread**, head-forward with **wings slightly drooped**, **wings-out** (horizontally lifted), distraction display with **tail spread and wings outspread** and **quivered**.

*36) Fitzpatrick S. 1998. Birds' tails as signaling devices: markings, shape, length, and feather quality. American Naturalist 151, 157–173. –* This is a comparative study of the entire avifauna of the Western Palearctic. **Tail displays** are present in 80% of species with elongated tails. The study concludes that tail-displays in a socio-sexual context may act as **handicaps** in the sense proposed by *Zahavi, A. 1975. Mate selection—a selection for a handicap. Journal of Theoretical Biology 53: 205-214*. This means that **socio-sexual selection** is responsible for the evolution of **tail displays** in birds.

*37) Holberton R.L., Able K.P., Wingfield J.C. 1989. Status signaling in dark-eyed juncos, Junco hyemalis: plumage manipulation and hormonal correlates of dominance. Animal Behaviour 37, 681-689. –* Experimental increase in the white patch on the tail that is flicked in social dominance interactions helped in attaining higher dominance status. This illustrates that an **experimental increase** in the **size** of motion-based **visual display** in the **social signaling** context **increases the signal efficiency** in the **dominance interactions** with conspecifics.

*38) Humphreys R.K., Ruxton G.D. 2020. Avian distraction displays: a review. –* The review shows that distraction displays **towards predators** are widespread among birds and might have evolved multiple times independently. The following forms of visual displays that involved tail or wings were mentioned in the review: **tail flagging**, **wing-out** run, **broken-wing** display, and erratic **fluttering**.

*39) Kenyon HL, Martin PR. 2022. Aggressive signaling among competing species of birds. PeerJ 10:e13431 DOI 10.7717/peerj.13431 –* Literature review and phylogenetic analyses across 164 species from 50 families and 24 orders. Results fully support the widespread use of plumage colors and motion-based visual displays in socio-sexual signals towards conspecifics and social visual displays to heterospecifics. **Opening of wings and tails** is listed among the visual displays analyzed in the paper and includes, among others: **wing outward-spreading, wing upward-spreading, wing raising upward, partial-spreading, tail trailing-fanned, tail raised-not fanned, tail raised-fanned, tail down-fanned, tail partly raised-fanned, tail partly raised-not fanned, tail side oriented fanned, and tail side oriented not fanned**.

*40) Martin S.G. 1970. The agonistic behario of varied thrushes (Ixoreus naevius) in winter assemblages. Condor 72, 452-459. –* An example of the good old classic descriptive papers with photos and illustrations representing visual displays of the thrushes in social interactions with conspecifics and heterospecific in winter flocks: **tail-up** display, head-forward display with **spread wings** and **spread tail**, **wing-quivering** display.

***41)** Peltier S.K., Wilson C.M. Godard R.D. 2019.Wing-flashing by northern mockingbirds while foraging and in response to a predator model. Northeastern Naturalist. 26, 251-260 –* Observational proof that **wing-flashing** is used for **flush-pursue** foraging and as a visual display **towards predators** in the mockingbird and in literature review of the antipredatory function of **wing-displays** in several other species from the same family (Mimidae).

***42)** Ramesh D., Lima S.L. 2019. Tail-flashing as an anti-predator signal in small wintering birds. Behavioral Ecology and Sociobiology (2019) 73:67 –* Results indicate that the Dark-eye junco's (*Junco hyemalis*) **tail-flash/tail-flick** displays are directed **towards predators** and may have antipredatory function (pursuit deterrence).

***43)** Randler C. 2016. Tail movements in birds – current evidence and new concepts. Ornithological Science 15, 1-14. –* This is a review paper. It considers different contexts of visual signaling **toward predators**, **towards prey**, **socio-sexual** signals towards conspecifics, and **alarm signals** to conspecifics about predators. It classifies and attempts to define or redefine tail-movement displays into several types: **upward-downward tail-flicking**, **side-to-side movements**, **tail-spreading**, **tail-fanning**, **side-to-side wagging**, **tail pumping**, and **tail wagging**. Tail movements are often associated with **wing-spreading**, and **wing-drooping**.

#### 6G) Examples of terrestrial agonistic contests associated with visual-displays in lekking birds that also use of wings for non-display functions inextricably linked to the visual-display behavior

Underlined **bold font**:
It marks descriptions that indicate behaviors for which enhancement of protowings area and forelimb movement range for visual display, would have been also likely beneficial for the hypothetical use of **wings in drag-based and lift-based fast forward, backward and other maneuvers and movements/leaps/braking of a movement toward or away from the opponent**.

Underlined normal font:
It contains examples of contests where forelimbs/wings are used to produce sound or to physically strike opponents during agonistic contests.

**** Examples of expressions describing behaviors that may involve the use of wings for functions other than the visual-display function in terrestrial lekking birds in the context if inter-sexual visual displays to opponents. The expressions are extracted from the papers listed below in the "List of examples of papers…"***

*Non-visual-signaling use of wings without physically hitting opponents:*
**- Chases**
**- Wing strokes during flutter jump**
- **Flutter-flight display**
**- Wing-Beats**
**- Jumping a few centimeters in the air**
**- Forward rushes**
**- Leap into the air**

*Use of wings to produce sound:*
- Wing-clapping
- Flutter-flight display

*Physical use of wings to hit the opponent:*
- smashing their wings down on the other male
- leaping/jumping at opponent
- lash out at each other with their wings,
- strike their opponent with feet, wings, and/or beak
- wing-beat fighting

**** List of examples of papers with excerpts from them describing the use of wings for non-display functions in modern bird in the context of agonistic interactions using visual displays for the signaling function:***

------------------------------------------------------------------------------------------------------------------------------------
**Lesser prairie chicken** (*Tympanuchus cupido*)

"Threat behavior during aggressive encounters between territorial males includes elevating tail, spreading and drooping primaries, exposing superciliary eye-combs, and elevating pinnae. **Chases are common** when males intrude into another male's territory"
"Males display by exposing and enlarging the superciliary eye-combs, elevating tail to highest extent, erecting pinnae and positioning them forward and parallel to the ground, drooping wings and spreading primaries, extending neck and head in forward position, stamping feet on ground and moving forward, and expanding esophageal air sacs and producing Booming vocalization. **Flutter Jump, or Wing-Beat**, appears to be stimulated by hens flying or walking onto arena; males may rise 2–3 m and often rotate 180° when alighting, sometimes onto shrubs. Usually **only a short burst of wing strokes during Flutter Jump**; birds keep their bodies fairly horizontal when landing."

*Hagen, C. A. and K. M. Giesen (2020). Lesser Prairie-Chicken (Tympanuchus pallidicinctus), version 1.0. In Birds of the World (A. F. Poole, Editor). Cornell Lab of Ornithology, Ithaca, NY, USA. https://doi.org/10.2173/bow.lepchi.01*

Movie clip:
https://www.youtube.com/watch?v=jYw42E9rBsk

---

**Greater Sage grouse** (*Centrocercus urophasianus*), **Gunnison sage grouse** (*Centrocercus minimus*)

**"**Many physical interactions occur on leks. Males may move back and forth in front of each other and settle down about 0.5 m apart, head to tail, in Face-Past, or Parallel Reversed, Display. If either male moves, the other moves with him, maintaining spacing and position. Males may remain in this position for >1 h, particularly at end of morning display period. Face-Past Display and other activities may lead to Wing Fights: Males crouch forward with their bodies parallel to the ground, lower their tails, and lash out at each other with their wings, occasionally **jumping a few centimeters in the air** and smashing their wings down on the other male. Males rarely peck at each other, but may grab other males with their bills and attempt to drag them"

*Schroeder, M. A., J. R. Young, and C. E. Braun (2020). Greater Sage-Grouse (Centrocercus urophasianus), version 1.0. In Birds of the World (A. F. Poole and F. B. Gill, Editors). Cornell Lab of Ornithology, Ithaca, NY, USA. https://doi.org/10.2173/bow.saggro.01*
*Young, J. R., C. E. Braun, S. J. Oyler-McCance, C. L. Aldridge, P. A. Magee, and M. A. Schroeder (2020). Gunnison Sage-Grouse (Centrocercus minimus), version 1.0. In Birds of the World (P. G. Rodewald, Editor). Cornell Lab of Ornithology, Ithaca, NY, USA. https://doi.org/10.2173/bow.gusgro.01*

Movie clips:
https://www.google.com/url?sa=t&source=web&rct=j&opi=89978449&url=https://www.youtube.com/watch%3Fv%3D5rJgfm_eLS0&ved=2ahUKEwi0qe_k2pmRAxXrh68BHSdGBpYQ3aoNegQIExAN&usg=AOvVaw29y6KgvJrBCQrAtXonaDWG

https://www.google.com/url?sa=t&source=web&rct=j&opi=89978449&url=https://www.youtube.com/watch%3Fv%3DeR8jJdntt_0&ved=2ahUKEwi0qe_k2pmRAxXrh68BHSdGBpYQ3aoNegQIGhAN&usg=AOvVaw3NY44SskqPs-W_ZSDoFcIY

https://www.google.com/url?sa=t&source=web&rct=j&opi=89978449&url=https://www.youtube.com/shorts/m0NDpi2moaw&ved=2ahUKEwi0qe_k2pmRAxXrh68BHSdGBpYQ3aoNegQIFBAN&usg=AOvVaw0UvOFlf2aAGTAvf1wZcJl4

https://www.google.com/url?sa=t&source=web&rct=j&opi=89978449&url=https://www.youtube.com/watch%3Fv%3DK-gPdZSmwU4&ved=2ahUKEwi0qe_k2pmRAxXrh68BHSdGBpYQ3aoNegQIGRAN&usg=AOvVaw1wPTVrvJHqXUXRUc6zlgB0

---

**Sharp-tailed grouse** (*Tympanuchus phasianellus*)

"Fights involve pecking and feather-pulling with bill, wing beating, and jabbing and scratching with nails of toes while 0-1 m in air."
"Fights rarely end in bloodshed, but some males are mortally wounded (MWG). Fights typically are preceded by **Forward Rushes** and associated with Face-offs, Cackles, Whines, and Dances"

*Connelly, J. W., M. W. Gratson, and K. P. Reese (2024). Sharp-tailed Grouse (Tympanuchus phasianellus), version 1.1. In Birds of the World (A. F. Poole, F. B. Gill, and M. G. Smith, Editors). Cornell Lab of Ornithology, Ithaca, NY, USA. https://doi.org/10.2173/bow.shtgro.01.1*

Movie clips:
https://www.google.com/url?sa=t&source=web&rct=j&opi=89978449&url=https://www.youtube.com/watch%3Fv%3DvUicA4gRC24&ved=2ahUKEwjMof_h3JmRAxWgqVYBHX3xFEIQ3aoNegQIGRAN&usg=AOvVaw3_rWJkWXb35lSWQtOXB7MR

https://www.youtube.com/watch?v=xLluaoSEtpA

---

**Greater prairie chicken** (*Tympanuchus cupido*)
'During aggressive encounters with other males, males lower their pinnae feathers, deflate their air sacs, **leap into the air**, and strike their opponent with feet, wings, and/or beak.
Males display by extending eye combs, lowering head, erecting pinnae feathers on neck, pointing tail slightly forward, stamping feet on the ground, tail clicking, stiffening, shaking, and dropping wings until the tips of the primaries touch the ground, expanding esophageal air sacs, and producing a booming vocalization"

*Johnson, J. A., M. A. Schroeder, and L. A. Robb (2020). Greater Prairie-Chicken (Tympanuchus cupido), version 1.0. In Birds of the World (A. F. Poole, Editor). Cornell Lab of Ornithology, Ithaca, NY, USA.* *https://doi.org/10.2173/bow.grpchi.01*

Movie clips:
https://www.youtube.com/watch?v=mAgLsHxcb8Y

https://www.youtube.com/watch?v=8vTfQsrkGdo

-------------------------------------------------------------------------------------------------------------------------------

**Spruce grouse** (*Canachites canadensis*)

"… most agonistic interactions occur between members of the same sex. Males typically respond to the presence of other males by partially erecting their tail and breast feathers, often by pecking aggressively at inanimate objects, and/or uttering the trill call. The territory occupant may perform any or all of the following displays: the Tail-swish Display, **Flutter-flight Display** and, in Franklin's Spruce Grouse, the **Wing-clap Display** in response to an intruder."

"When fights occur between males or when a territorial male is presented with a mirror image of itself, it attacks the head, neck, and back of the intruding male or image with feet, bill, and wings.

*Schroeder, M. A., E. J. Blomberg, D. A. Boag, P. Pyle, and M. A. Patten (2021). Spruce Grouse (Canachites canadensis), version 1.1. In Birds of the World (P. G. Rodewald, Editor). Cornell Lab of Ornithology, Ithaca, NY, USA. https://doi.org/10.2173/bow.sprgro.01.1*

-------------------------------------------------------------------------------------------------------------------------------

**Ring-necked pheasant** (*Phasianus colchicus*)

"Most aggressive encounters occur during breeding season, especially during territory establishment. An escalating series of threat displays (see below) may culminate in physical interaction if one bird does not flee. Fights involve fluttering up together, breast to breast, biting at each other's wattles, or making alternate **high leaps forward at opponent**, using bill, claws, and spurs. Fatalities are rare; one bird usually withdraws at early stage; the victor usually chases loser away after combat, running with head low."

Ring-necked pheasant clip:
https://www.google.com/url?sa=t&source=web&rct=j&opi=89978449&url=https://www.youtube.com/watch%3Fv%3DjfdqM7zhMkQ&ved=2ahUKEwiAmavWxJqRAxVeklYBHaOrOCIQ3aoNegQIEhAN&usg=AOvVaw2EzAeGJfUaDb60WjpSDS6N

*Giudice, J. H., J. T. Ratti, and S. G. Mlodinow (2022). Ring-necked Pheasant (Phasianus colchicus), version 1.1. In Birds of the World (N. D. Sly, Editor). Cornell Lab of Ornithology, Ithaca, NY, USA.* *https://doi.org/10.2173/bow.rinphe1.01.1*

-------------------------------------------------------------------------------------------------------------------------------

**Black grouse** (*Lyrurus tetrix*)

"Flutter-jumping (Fig. 2). The introductory part of flutter-jumping is identical to crowing in the erect posture, but instead of flapping the wings weakly, the **male flies up to a height of 1 to 2 meters** and over a distance of up to 10 or 15 meters. Simultaneously, the crowing sound is produced and repeated. During flutter-jumping the white underparts of the wings are very conspicuous."

"Males frequently threaten each other along the boundaries of the display sites. Threat consists of **tentative approaches and withdrawals**, performed in postures that vary between the horizontal display posture and a modification of this posture which contains elements seen in the erect posture'

"During fighting, males **jump up at each other**, often crowing simultaneously. Although the legs are used to kick the opponent, the bill is used more often; vicious pecking occurs often directed especially at the head of the opponent. In addition the fighting birds hit each other by beating their wings."

*Kruijt J.P. Hogan J.A. 1967. Social behavior on the lek in the black grous, Lyrurus tetrix tetrix (L.). Ardea. 55: 204-240.*

Black grouse clips:
https://www.google.com/url?sa=t&source=web&rct=j&opi=89978449&url=https://www.youtube.com/watch%3Fv%3DbM3rOHxrlQc&ved=2ahUKEwjMof_h3JmRAxWgqVYBHX3xFEIQ3aoNegQIExAN&usg=AOvVaw1iYQ8GG-D_0IbpKAlVoAX-

https://www.youtube.com/watch?v=P7osXtZEDRM

------------------------------------------------------------------------------------------------------------------------------------

**Capercaillie** (*Tetrao*)

"Fighting with wing beats and/or pecking... is initiated and occasionally terminated with deep bowing by both rivals/ The birds often try to start beak pecking or wing-beat fighting"

*Porkert J., Solheim R., Flor A. 1997. Behaviour of hybrid male Tetrao tetrix x Tetrao urogallus on black grouse leks. Wild. Biol. 3: 169-176.*

Movie clip of capercaillie:
https://www.youtube.com/watch?v=Comh75ZS7eY

------------------------------------------------------------------------------------------------------------------------------------

**Rock ptarmigan** (*Lagopus muta*)
"Males interact aggressively when one intrudes on another's territory. Contests usually highly ritualized (e. g., Parallel Walking; when opponents are territorial neighbors; rarely involving physical contact (Attacks)."
"Attacking males strike opponent with wings and beak.'

*Montgomerie, R. and K. Holder (2020). Rock Ptarmigan (Lagopus muta), version 1.0. In Birds of the World (S. M. Billerman, B. K. Keeney, P. G. Rodewald, and T. S. Schulenberg, Editors). Cornell Lab of Ornithology, Ithaca, NY, USA. https://doi.org/10.2173/bow.rocpta1.01*

------------------------------------------------------------------------------------------------------------------------------------

**Ruffed grouse** (*Bonasa umbellus*)

"Fights rarely observed in wild, but common in captivity. Fighting Ruffed Grouse holds feathers close to body, tail folded and dropped, head and neck lowered and outstretched. May attempt to peck opponent. Sometimes both individuals stand erect, peck, and claw opponent; may beat each other with wings."

*Rusch, D. H., S. Destefano, M. C. Reynolds, and D. Lauten (2020). Ruffed Grouse (Bonasa umbellus), version 1.0. In Birds of the World (A. F. Poole and F. B. Gill, Editors). Cornell Lab of Ornithology, Ithaca, NY, USA. https://doi.org/10.2173/bow.rufgro.01*

------------------------------------------------------------------------------------------------------------------------------------

**Wild turkey** (*Meleagris gallopavo*)

"Fighting begins with mutual threat and progresses to striking with wings and kicking. Eventually one bird grabs the other's beak or snood and birds entwine necks, pushing against each other with breasts. Fights usually end by one bird gaining advantage and getting beak hold on skin of back of opponent's neck. With this hold, winning bird forces opponent's head to ground, until loser is able to twist free"

*McRoberts, J. T., M. C. Wallace, and S. W. Eaton (2020). Wild Turkey (Meleagris gallopavo), version 1.0. In Birds of the World (A. F. Poole, Editor). Cornell Lab of Ornithology, Ithaca, NY, USA. https://doi.org/10.2173/bow.wiltur.01*

Movie clips of fight/display:
https://www.google.com/url?sa=t&source=web&rct=j&opi=89978449&url=https://www.youtube.com/watch%3Fv%3DmH8e8Gx_3E4&ved=2ahUKEwjLuOHbhJqRAxVX1zQHHTlrIt8Q3aoNegQIEhAN&usg=AOvVaw1I26OwILIDssG8TYjDPKW-

------------------------------------------------------------------------------------------------------------------------------------

#### PART 7: Description of Supplementary Videos

**Video S1.** The greater roadrunner visually flushing a lizard.

**Video S2.** Pilot animations loosely based on the display of *Cercotrichas galactotes*.

**Video S3.** Hypothetical flushing dinosaur animations imitating the display of *Geococcyx californianus*, used in experiment 1.

**Video S4.** Hypothetical flushing dinosaur animations imitating the display of *Cercotrichas galactotes*, used in experiment 2.

**Video S5.** Hypothetical flushing dinosaur animations imitating the display of *Mimus polyglottos*, used in experiment 3.

**Video S6.** Hypothetical flushing dinosaur animations imitating the display of *Geococcyx californianus* and a walking stimulus of 1 m/sec in Experiment 4

### PART 8: Supplementary References


1. Park, J. *et al.* Escape behaviors in prey and the evolution of pennaceous plumage in dinosaurs. *Sci. Rep.* **14**, 549 (2024).
2. Senter, P. & Robins, J. H. Resting orientations of dinosaur scapulae and forelimbs: A numerical analysis, with implications for reconstructions and museum mounts. *PLoS One* **10**, 1–21 (2015).
3. Witmer, L. M. The extant phylogenetic bracket and the importance of reconstructing soft tissues in fossils. in *Functional morphology in vertebrate paleontology* (ed. Thomason, J. J.) 19–33 (Cambridge University Press, Cambridge, 1995).
4. Senter, P. Comparison of forelimb function between *Deinonychus* and *Bambiraptor* (Theropoda: Dromaeosauridae). *J. Vertebr. Paleontol.* **26**, 897–906 (2006).
5. Senter, P. & Robins, J. H. Range of motion in the forelimb of the theropod dinosaur *Acrocanthosaurus atokensis*, and implications for predatory behaviour. *J. Zool.* **266**, 307–318 (2005).
6. Zanno, L. E., Gillette, D. D., Albright, L. B. & Titus, A. L. A new North American therizinosaurid and the role of herbivory in 'predatory'dinosaur evolution. *Proc. R. Soc. B Biol. Sci.* **276**, 3505–3511 (2009).
7. Ji, Q., Currie, P. J., Norell, M. A. & Ji, S.-A. Two feathered dinosaurs from northeastern China. *Nature* **393**, 753–761 (1998).
8. Wings, O. A review of gastrolith function with implications for fossil vertebrates and a revised classification. *Acta Palaeontol. Pol.* **52**, 1–16 (2007).
9. O'Connor, J. K. & Zhou, Z. The evolution of the modern avian digestive system: insights from paravian fossils from the Yanliao and Jehol biotas. *Palaeontology* **63**, 13–27 (2020).
10. Johnson, D. R. Diet and estimated energy assimilation of three Colorado lizards. *Am. Midl. Nat.* **76**, 504–509 (1966).
11. Sokol, O. M. . Lithophagy and geophagy in reptiles. *J. Herpetol.* **5**, 69–71 (1971).
12. Downs, C. T., Bredin, I. P. & Wragg, P. D. More than eating dirt: a review of avian geophagy. *African Zool.* **54**, 1–19 (2019).
13. Ma, W., Pittman, M., Lautenschlager, S., Meade, L. E. & Xu, X. Functional morphology of the oviraptorosaurian and scansoriopterygid skull. in *Bull. Am. Mus. Nat. Hist* vol. 440 229–250 (2020).
14. Hughes, J. M. Greater Roadrunner (*Geococcyx californianus*), version 1.0. In Birds of the World (A. F. Poole, Editor). *Cornell Lab of Ornithology, Ithaca, NY, USA* (2020) doi:10.2173/bow.greroa.01.
15. Farnsworth, G., Londono, G. A., Martin, J. U., Derrickson, K. C. & Breitwisch, R. Northern Mockingbird (*Mimus polyglottos*), version 1.0. In Birds of the World (A. F. Poole, Editor). *Cornell Lab of Ornithology, Ithaca, NY, USA* (2020) doi:10.2173/bow.normoc.01.
16. Fotowat, H., Harrison, R. R. & Gabbiani, F. Multiplexing of motor information in the discharge of a collision detecting neuron during escape behaviors. *Neuron* **69**, 147–158 (2011).
17. Gabbiani, F., Krapp, H. G. & Laurent, G. Computation of object approach by a wide-field, motion-sensitive neuron. *J. Neurosci.* **19**, 1122–1141 (1999).
18. Judge, S. J. & Rind, F. C. The locust DCMD, a movement-detecting neurone tightly tuned to collision trajectories. *J. Exp. Biol.* **200**, 2209–2216 (1997).
19. Santer, R. D., Yamawaki, Y., Rind, F. C. & Simmons, P. J. Preparing for escape: An examination of the role of the DCMD neuron in locust escape jumps. *J. Comp. Physiol. A Neuroethol. Sensory, Neural, Behav. Physiol.* **194**, 69–77 (2008).
20. Simmons, P. J., Rind, F. C. & Santer, R. D. Escapes with and without preparation: the neuroethology of visual startle in locusts. *J. Insect Physiol.* **56**, 876–883 (2010).
21. Hailman, J. P. A field study of the mockingbird's wing-flashing behavior and its association with foraging. *Wilson Ornithol. Soc.* **72**, 346–357 (1960).
22. Moon, J.-Y. Field Study of Orthopteran Escape Pathway's Response to Robotic Simulation of Visual Displays of Flush-Pursue Predators. (Seoul National University, 2013).
23. Hancock, J. A., Stevens, N. J. & Biknevicius, A. R. Elegant-crested Tinamous *Eudromia elegans* do not synchronize head and leg movements during head-bobbing. *Ibis (Lond. 1859).* **156**, 198–208 (2014).
24. Miall, R. C. The flicker fusion frequencies of six laboratory insects, and the response of the compound eye to mains fluorescent 'ripple'. *Physiol. Entomol.* **3**, 99–106 (1978).
25. Rind, F. C. & Simmons, P. J. Orthopteran DCMD neuron: A reevaluation of responses to moving objects. I. Selective responses to approaching objects. *J. Neurophysiol.* **68**, 1654–1666 (1992).
26. Rind, F. C. & Bramwell, D. I. Neural network based on the input organization of an identified neuron

signaling impending collision. *J. Neurophysiol.* **75**, 967–985 (1996).
27. Branco, T. & Redgrave, P. The neural basis of escape behavior in vertebrates. *Annu. Rev. Neurosci.* **43**, 417–439 (2020).
28. Card, G. M. Escape behaviors in insects. *Curr. Opin. Neurobiol.* **22**, 180–186 (2012).
29. Hemmi, J. M. & Tomsic, D. The neuroethology of escape in crabs: from sensory ecology to neurons and back. *Curr. Opin. Neurobiol.* **22**, 194–200 (2012).
30. Peek, M. Y. & Card, G. M. Comparative approaches to escape. *Curr. Opin. Neurobiol.* **41**, 167–173 (2016).
31. Rind, F. C. Recent advances in insect vision in a 3D world: looming stimuli and escape behaviour. *Curr. Opin. Insect Sci.* **63**, 101180 (2024).
32. Gray, J. R. Habituated visual neurons in locusts remain sensitive to novel looming objects. *J. Exp. Biol.* **208**, 2515–2532 (2005).
33. Dececchi, T. A., Larsson, H. C. E. & Habib, M. B. The wings before the bird: An evaluation of flapping-based locomotory hypotheses in bird antecedents. *PeerJ* **2016**, (2016).
34. Kuznetsov, A. N. & Panyutina, A. A. Where was WAIR in avian flight evolution? *Biol. J. Linn. Soc.* 1–12 (2022) doi:10.1093/biolinnean/blac019.
35. Talori, Y. S. *et al.* Winged forelimbs of the small theropod dinosaur *Caudipteryx* could have generated small aerodynamic forces during rapid terrestrial locomotion. *Sci. Rep.* **8**, 1–14 (2018).
36. Novas, F. E., Agnolin, F., Brissón Egli, F. & Lo Coco, G. E. Pectoral girdle morphology in early-diverging paravians and living ratites: implications for the origin of flight. in (ed. History, A. M. of N. H. B. of the A. M. of N.) (American Museum of Natural History, 2020).
37. Dial, K. P. Wing-assisted incline running and the evolution of flight. *Science (80-. ).* **299**, 402–404 (2003).
38. Dial, K. P., Jackson, B. E. & Segre, P. A fundamental avian wing-stroke provides a new perspective on the evolution of flight. *Nature* **451**, 985–989 (2008).
39. Ostrom, J. H. Archaeopteryx and the origin of flight. *Q. Rev. Biol.* **49**, 27–47 (1974).
40. Reynolds, P. S. & Lima, S. L. Direct use of wings by foraging woodpeckers. *Wilson Bull.* **106**, 408–411 (1994).
41. Padian, K. *The Origin of Birds and the Evolution of Flight*. (California academy of sciences, 1986).
42. Lee, M. S. Y., Cau, A., Naish, D. & Dyke, G. J. Sustained miniaturization and anatomical innovation in the dinosaurian ancestors of birds. *Science (80-. ).* **345**, 562–566 (2014).
43. Brusatte, S. L., Lloyd, G. T., Wang, S. C. & Norell, M. A. Gradual assembly of avian body plan culminated in rapid rates of evolution across the dinosaur-bird transition. *Curr. Biol.* **24**, 2386–2392 (2014).
